# Elucidating biosynthetic pathways related to the synthesis of small halogenated peptidic natural products in marine sponge microbiomes

**DOI:** 10.64898/2026.08.03.742642

**Authors:** Catarina Loureiro, Michelle Schorn, Mohammad Alanjary, Ben Kuipers, Joris Louwen, John van der Oost, Marnix H. Medema, Detmer Sipkema

**Affiliations:** Laboratory of Microbiology, Wageningen University, Stippeneng 4, 6708WE Wageningen, The Netherlands; Bioinformatics Group, Wageningen University, Droevendaalsesteeg 1, 6708PB Wageningen, The Netherlands

## Abstract

Marine sponges are known sources of bioactive natural products (NPs), many of which are produced by associated bacterial symbionts via encoded biosynthetic gene clusters (BGCs). A particularly interesting subclass of sponge-derived NPs is comprised of small, brominated alkaloids, which are recovered from diverse habitats and host sponge taxonomies. Despite having been described decades ago, most of these NPs do not have an elucidated biosynthetic origin. We queried metagenomes of several sponge species by making use of a minimal set of core enzymes that we postulate to be necessary to produce these small peptidic NPs: an FADH2-dependent halogenase and an AMP-binding adenylation enzyme. This revealed a variety of novel BGC architectures, many of which showed conservation among sponge host phylogenies and were encoded in the genomes of diverse sponge-associated bacteria. Furthermore, we identified a BGC in the sponge *G. barretti* that is potentially linked to the production of the iconic barettins, given its enzymatic machinery and specific acidobacterial origin. The present work contributes to the challenging quest to link orphan brominated NPs to their parent BGCs in the sponge holobiont and beyond.

## Introduction

Marine sponges are ancient multicellular organisms that inhabit varied aquatic habitats as heterotrophic sessile filter feeders (1–3). These organisms host rich and diverse communities of microbial symbionts in their tissues (4–6), in a symbiosis where the microbes provide organic nutrients and protection against pathogens (2,7,8), in return for shelter and nutrition from the sponge host (9). Marine sponge holobionts are known as rich sources of diverse bioactive natural products (NPs), and their bacterial symbionts are believed to be responsible for the production of many of these specialized metabolites (10–14).

A particular feature of the marine environment and the sponge holobiont is the enrichment in bromide-halogenated NPs (15,16). Halogenated molecules show increased stability and membrane permeability, and exhibit a wide range of pharmaceutically interesting biological activities (17,18). Within this diverse repertoire are the bromotyrosine alkaloids, which constitute one of the most abundant, diverse and ubiquitous groups of marine-sponge-derived NPs (14,19,20). These molecules are reported across sponge genera and geographic locations and are thought to function as chemical defenses. Well known examples include the aerophobins, fistularins and psammplins, which undergo chemical transformations that release hydrogen cyanide from these brominated alkaloids in response to sponge tissue damage (21–24). These molecules are found in, among others, the Demosponge genus *Aplysina,* which has been reported to produce numerous brominated alkaloid NPs that can make up to 12% of the sponge dry weight (5,25–27). Another interesting class of abundant and diverse sponge NPs are (brominated) tryptophan-derived indole alkaloids, which have shown potent and diverse bioactivities (20,28,29). A well-known producer of this class of NPs is the Demosponge *Geodia barretti,* which is known to produce several bromotryptophan derivatives, including the bioactive diketopiperazine (DKP) containing barettins (30–33). DKPs are “head-to-tail” cyclic dipeptides, featuring a six-membered piperazine ring, that shows a wide variety of bioactivities, such as antimicrobial and anti-inflamatory (34,35)).

Even though for the large majority of marine halogenated NPs their biosynthetic origin remains unresolved, several have been associated to specific bacterial sponge symbionts (36–41). Recent studies on the presence and distribution of brominated alkaloid NPs in sponges have shown that despite the widespread presence of these NPs, there are significant correlations between microbiota and metabolomes of several phylogenetically distant sponges (14,42). The authors also point to specific symbionts that are likely responsible for the production of these NPs in *G. barretti* (42)). Nevertheless, there is no consensus on the origin of brominated alkaloids and a bacterial symbiont has yet to be associated with the production of NPs, which are widely distributed across sponge host phylogeny (14). In *Aplysina aerophoba* sponges, bromotyrosines are located in spherulous cells in the sponge mesohyl, which in turn points towards the sponge itself as their producer (27,43).

The enzymatic machinery for the biosynthesis of bacterial NPs is in its majority genetically encoded in Biosynthetic Gene Clusters (BGCs) (44) but despite recent advances in technologies and methodologies for generating, annotating, and mining metagenomic data from complex environmental communities (45), the characterization of the sponge holobiont biosynthetic landscape remains challenging with (i) only a few studies performing untargeted sponge metagenome mining, (ii) large sequence diversity gaps between sponge-derived genes/proteins/BGCs and those in public databases and (iii) a general lack of sponge symbiont cultured representatives. In this way, we are left with scarce examples of chemically characterized sponge-derived enzymes (46–48). A testament to this challenge is the fact that the extensive repertoire of sponge-derived brominated alkaloid NPs has been described for decades, with only a few molecules having a known biosynthetic origin, such as the *bmp* BGC, which directs the production of several brominated phenols/pyrroles (37). It is intuitive to start the efforts to decipher the biosynthesis of a chemical compound by examining its structure and generating a hypothetical BGC architecture which can then be used to mine metagenomic data. However, this approach has its caveats: the hypothetical BGC is thought out based on biosynthetic strategies likely described for distant biomes and bacterial hosts, and evolutionary distance of genes/proteins might lead to erroneous or missing functional annotations. As an example, a recent study showed that a large fraction of halogenase sequences derived from amplicon sequencing in *Aplysina* species did not produce quality hits (*e*-value < 0.001) to reference sequences (48).

In this study, we aimed to explore the diversity of sponge bacterial symbiont BGCs that are likely related to the production of small halogenated peptidic NPs and/or their precursors. Peptidic NPs are generally thought to be nonribosomal peptides (NRPs) or ribosomally synthesized and post-translationally modified peptide (RiPPs), encoded by canonical BGC architectures. However, recent studies that explored the diversity of sponge-derived RiPP BGCs featuring halogenases have mostly reported an expanded diversity of (sponge-derived) proteusin BGCs (46,47). Additionally, RiPP BGCs are often responsible for the production of large peptides, whilst sponge-derived brominated alkaloids generally feature a small number of amino acids (14,49,50). Thus, we have chosen not to consider RiPP BGCs in this study. Following the biosynthetic logic behind NRP biosynthesis, we hypothesize that the first domain in the enzymatic cascade, an adenylation (A) domain, would be necessary to select and activate the amino acid substrates (51). We decided to not include any further NRPS core domains in the search to avoid missing novel BGC architectures that do not comply with the currently described biosynthetic routes. In fact, deemphasizing the presence of core domains in the search for novel BGCs has already proven fruitful, and is likely to drive a next wave of discovery in NP biosynthesis (52–55). A prime example of a BGC lacking a canonical architecture, which constitutes perhaps one of the few successes in elucidating the biosynthesis of sponge symbiont derived brominated NPs is the *bmp* BGC, which directs the production of several brominated phenols/pyrroles (37,56,57). Analogous to our query, the *bmp* BGC features a standalone A domain, as well as two (FADH_2_)-dependent halogenases (FDHs). Other examples of BGCs leading to the production of brominated alkaloids and their precursors featuring these two enzymes include those responsible for the production of the bromoalterochromides, pyoluteorin, kutznerides and largimiycin (41,58–61). Finally, the presence of a halogen is a defining feature in this class of NPs and mining for halogenating enzymes in sponge metagenomes has shown to be a successful strategy to identify novel BGCs (37,46). More specifically, we have chosen to focus on bacterial FDHs, which halogenate a variety of tryptophan derivatives and aromatic substrates, and are often encoded as part of BGCs (37,60,62–66). Following this line of thinking, we explored sponge metagenomes through semi-untargeted mining for small halogenated peptidic NP producing BGCs, anchored in two main enzymatic principles: the presence of (1) a halogenase domain, and (2) an adenylation domain. The genes encoding for these domain-containing enzymes thus constitute the core genes, i.e. the genes responsible for the synthesis of the backbone of the compound, for the Halo_AMP BGC pseudoclass.

We queried the metagenomes of 15 host sponge species, sampled across diverse geographical locations and depths (Table S1) by leveraging recently developed tools to mine (meta)genomes specifically for candidate BGCs (67–71). We provide an overview of the novel BGC architectures featured within the Halo_AMP pseudo-class, as well as their distribution across the sponge and bacterial hosts. Furthermore, we identify a Gene Cluster Family (GCF) showing similarities to the *bmp* BGC, which includes a BGC variant recovered only from *G. barretti* metagenomes. A combination of enzymatic machinery and taxonomy of the predicted bacterial host indicate a potential link between this BGC variant and the well-known barettins.

## Materials and Methods

### Sponge collection

*Aciculites cribrophora* (DOM43), *Smenospongia* sp. (DOM10), *Caminus* sp. (DOM14), *Petrosia hartmani* (DOM40), and *Aplysina* sp. (DOM33) samples were collected and processed in our previous study (72), *Aplysina aerophoba* (Aply16-23) samples were collected and processed in another previous study (73); *Geodia barretti* (gb1-10), Seawater (sw7-9, gb1_f-gb10_f) and *Petrosia ficiformis* (Pf4-Pf12) samples were collected and processed in yet another previous study (47). *Geodia atlantica* (gb3) was collected during the same sampling effort and processed as described in our previous study (47). *Aplysina* sp. (DOM012A), *Siphonodictyon coralliphagum* (DOM005), *Svenzea zeai* (DOM015, DOM013), *Verongula* sp. (DOM026, DOM045), *Neopetrosia* spp. (DOM044, DOM011), *Xestospongia muta* (DOM049), and *Oceanapia bartschi* (DOM057, DOM016, DOM007A-B) were collected during the same sampling effort (Dominica, March 2016, Table S1). The sponges were morphologically identified by prof. Nicole De Voogd (Naturalis, The Netherlands) and processed as in our previous study study (72).

### Total DNA extraction and metagenomic sequencing

#### Illumina short reads

*A. aerophoba, G. barretti* and *P. ficiformis* sponge samples and seawater samples were processed in our previous study (47) and for the *G. atlantica* sample the same procedure was followed. *A. cribrophora* (DOM43)*, Smenospongia* sp. (DOM10), *Caminus* sp. (DOM14) and *Aplysina* sp. (DOM33) samples were processed in our previous study for generation of Illumina shotgun metagenomic sequencing data (72), and for the remaining ‘DOM’ samples the same procedure was followed.

#### Nanopore long reads

*A.cribrophora, Smenospongia* sp., *Caminus* sp. and *Aplysina* sp. (DOM33) samples were processed in our previous study (X) for generation of Nanopore long read sequencing data and for *P. hartmani the same procedure was followed*.

### Metagenomic sequence processing, assembly and binning

*A. aerophoba, G. barretti, P. ficiformis* and seawater Illumina reads were processed, assembled and binned in our previous study (47), and for the *G. atlantica* sample the same procedure was followed. *A.cribrophora*, *Smenospongia* sp., *P. hartmani*, *Caminus* sp. and *Aplysina* sp. (DOM33) Illumina reads were processed and assembled in our previous study (72) and for the remaining ‘DOM’ samples the same procedure was followed. *A.cribrophora*, *Smenospongia* sp., *P. hartmani*, *Caminus* sp. and *Aplysina* sp. (DOM33) reads were binned and classified following the same strategy used for *A. aerophoba* contigs (47) using hybrid long and short read data, and for the remaining ‘DOM’ samples the same strategy was used as for *P. ficiformis* contigs (47), using short read data only.

Metagenome assembled genome (MAG) reassembly was done with the MetaWRAP (74) reassembly module, and MAG relative abundance in host sample was predicted using GTDB-Toolkit (75) v1.1.0 (GTDB-Tk) quantify module. The obtained bins were dereplicated using dRep (76) v2.5.4 with default parameters for primary clustering and secondary clustering using parameters --S_algorithm gANI --S_ani 0.95 (77). MAG assembly features can be found in Table S8.

Querying of Acidobacterial amplicon sequence variants (ASVs) detected in *G. barretti* (42) was done using nucleotide BLAST (78) using the MAG gb6_2_bin.52 sequences as a database.

### BGC & GCF prediction and analysis

BGCs of all samples and reassembled MAGs were predicted using antiSMASH v6 (79), with a custom BCG detection rule ‘Halo_AMP’ featuring category: other, cutoff: 5, neighbourhood: 20, conditions: Trp_halogenase and AMP-binding. The bitscore cutoff value for detection of the tryptophan halogenase PFAM (PF04820) was set to 10. When the region predicted by antiSMASH featured hybrid, interleaved, or neighboring candidate clusters, only the Halo_AMP single cluster was considered. Presence of additional regulatory elements in manually selected BGCs was accessed with antiSMASH v7 beta (80). Predicted Halo_AMP BGCs were grouped into GCFs with BiG-SCAPE (67) using parameters --mix -v --mode auto -- mibig --cutoffs 0.3 --include_singletons. GCFs were further annotated by assessing diversity of BGC sponge hosts and bacterial taxonomic origins by making use of in-house Python scripts available at https://github.com/CatarinaCarolina/sponge_halo_amp.

#### Connected component (CC) A analysis and curation

BGCs present in the BiG-SCAPE connected component (CC) A (GCFs 2249 2410 2286 2173 2159), as well as those in related GCFs, i.e. GCFs harbouring the same architectural features as seen in CC A (10, 15, 2023) were singled out for further characterization. In order to generate the most complete version of these BCGs possible, we searched for the BGC homologues in the reassembled versions (reBGC) of their MAGs of origin, and kept these when length(BGC) < length(reBGC). An exception to this rule was made for the DOM43 sample, where the reBGC was present in a shorter contig but the BGC featured a likely assembly issue leading to an oversized core gene/CDS). The diversity of the curated set of CC A BGCs was analyzed using CORASON (67) v1, with parameters -g --cluster_radio 200, with gb8_2 contig 2489 gene 1 encoding the A domain as a query and gb10_c02489_haloAMP as the reference BGC.

#### Core and tailoring gene analysis

Prediction of functional class and substrate specificity of adenylation domain active-site-specificity-conferring residues was conducted with AdenylPred (81), using default settings. PARAS-residues v0.0.3 (82) was used to extract and align protein sequences of A-domain active site and stachelhaus residues. Geneious Prime v2022.2.2 (https://www.geneious.com) was used to compute average pairwise identity percentages of these alignments. Sequence logos were generated using WebLogo Version 2.8.2. Coding region sequences (CDS) of all genes annotated as a halogenase by antiSMASH (sec_met_domain=Trp_halogenase) were extracted and combined with an expanded public dataset (Fisher et al.’s dataset) of flavin-dependent halogenases (Table S2), including additional phenol and pyrrole halogenases not present in the Fisher’s dataset (65). These sequences were used to generate a limited sequence similarity network (SSN) using the EFI-EST webserver (83) with an E-value cut-off of 5 and an alignment score of 70 (∼35% minimal sequence identity). Sequences were clustered at 50% pairwise identity, and the network was visualized using Cytoscape v3.9.1 (84). CDSs of all genes annotated as radical SAM (rSAM) domain (sec_met_domain=PF04055) by antiSMASH were extracted and combined with a public dataset of chemically characterized rSAM sequences (85). These sequences were used to generate a limited sequence similarity network (SSN) using the EFI-EST webserver (83) with an E-value of 5 and an alignment score of 45 (∼30% minimal sequence identity). Sequences were clustered at 90% pairwise identity, and the network was visualized using Cytoscape v3.9.1 (84). SSN similarity and clustering parameters were designed to follow methodologies from the reference publications (65,85). CDSs of all genes annotated as a cytochrome p450 by antiSMASH (sec_met_domain= p450) were extracted and combined with the sequences of all single-domain p450-encoding genes (matching PF00067) in the SwissProt Database (86), as well as the sequences of bmp7 and bmp12 (38). The protein sequences were aligned using MAFFT (87) local iterative mode (-linsi option), and a phylogenetic tree was constructed using FastTree (88). The sequences were additionally characterized by assigning KEGG Orthologs (KO) (89) via HMMsearch and CYP family assignment was inferred via best Basic Local Alignment Search Tool (BLAST) hit to the cytochrome p450 reference set from CYPMiner (90). The tree and annotations were visualized with iTOL (91). All CDSs were compared via BLAST against cyclodipeptide synthase (CDPS) homologues and other genes in the *ank* BGC (52).

## Results and Discussion

### Halo_AMP pseudoclass gene clusters are enriched in biosynthetic domains

In total, 840 BGCs of the Halo_AMP pseudoclass were detected in sponge and seawater samples, which were clustered into 267 GCFs and 153 singletons. Of the recovered GCFs, 89% are exclusively sponge-derived, 10% are exclusively seawater-derived, 1% includes sponge- and seawater-derived BGCs and none included experimentally characterized reference BGCs from the Minimum Information about a Biosynthetic Gene cluster (MIBiG) database (92). The marine sponge holobiont thus continues to be a rich source of novel biosynthetic potential. We saw that, in agreement with our previous study on diversity and conservation of GCFs across host sponge phylogeny (47), most Halo_AMP GCFs detected are host-specific, with 72% of GCFs originating from a single host sponge species. We also aimed to decipher the bacterial origin of these GCFs by investigating the presence of their parent contigs in the generated MAGs. We found that 23% of GCFs are predicted to originate from Proteobacteria, 16% from Acidobacteriota, 9% from Latescibacterota, and 8% from Poribacteria (Table S3). These taxa are all known symbionts of sponges and their biosynthetic potential has also been previously noted (47). For 30% of GCFs a taxonomic origin could not be assigned with this approach as the parent contigs were not present in MAGs. To provide an overview of the representative BGC architectural features detected here, we considered a less stringent BGC similarity cutoff than the GCF, i.e., the BiG-SCAPE-generated connected component (CC), and have selected 20 CCs for a deeper analysis (Table S3.2). For CCs, we see higher levels of conservation among host sponge species (Fig1.a), with several CCs featuring BGCs detected in multiple host species, contrarily to what is seen at GCF level. This suggests a functional redundancy and conservation of key characteristics of the Halo_AMP biosynthetic strategy, paired with divergent evolution taking place in each holobiont.

The BCGs detected here were enriched in tailoring enzymes commonly featured in biosynthetic regions, an indication that diversifying the BGC search rules and steering away from the canonical core enzymes can provide access to a novel reservoir of biosynthetic routes. Detected genes encoding non-core biosynthetic enzymes (Fig. 1.b, Table S4) include several types of oxidoreductases (93) such as cytochrome P450 monooxygenases (P450s) (94) and radical S-adenosylmethionine enzymes (rSAMs) (95), as well as group transfer enzymes (96), pyridoxal phosphate (PLP)-dependent enzymes (97), thioesterases, peptidases and other lyases/hydrolases. Furthermore, also characteristic of biosynthetic regions is the presence of genes encoding several transporters, including the MATE (Multidrug And Toxic Compound Extrusion) family (98), ABC transporters, and TonB-dependent transporters (TBDT) (99). Although TBDTs are generally associated with metallophore BGCs, and thus surprising to detect in BGCs without other metallophore characteristics, they have also been found as potential antimicrobial resistance genes in lassopeptide BGCs (100,101) and could therefore also have a non-canonical role here. Another class of detected genes that are less frequently reported in BGCs and could be fulfilling a non-canonical role are those coding for toxin-antitoxin (TA) and nucleotide binding domains, as these have also been linked to regulators identified in the context of antimicrobial (resistance) and quorum sensing systems (102–104). We also observe the presence of the poorly characterized PQQ-like domain in our detected gene clusters at high frequency. This domain is not commonly featured in BGCs (not present in MIBiG (92) BGCs, and found only in two antiSMASH-DB BGCs (105)). Finally, ORFs designated as hypothetical proteins, with domains of unknown function (DUFs) or with no hits to databases were common in these BGCs, an indication of potentially novel enzymatic function.

**Figure 1.**
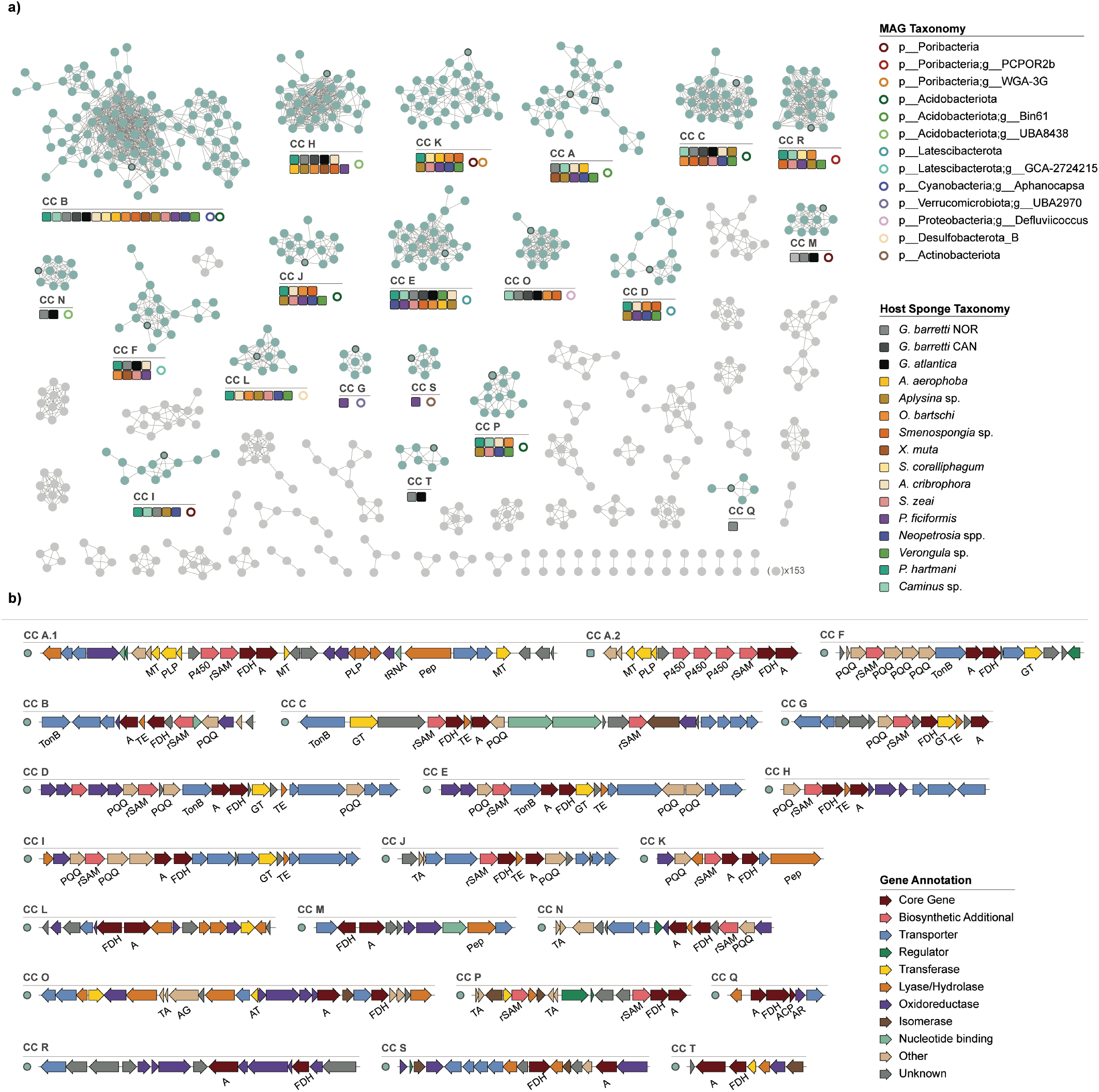
Diversity of identified gene clusters encoding both halogenases and AMP-binding domains. **a)** BGCs represented in a BiG-SCAPE generated similarity network. Colored nodes represent the connected components (CC) described. Each CC is annotated with sponge host taxonomy and bacterial taxonomy of its BGCs. **b)** Representative BGCs are depicted and annotated with predicted gene functions (A: AMP-binding domain, FDH: flavin dependent halogenase, PQQ: Pyrrolo-quinoline quinone, MT: methyl transferase, Pep: peptidase, GT: glycotransferase, TE: thioesterase, rSAM: radical S-adenosylmethionine enzyme, P450: cytochrome P450 monooxygenase, TonB: TonB-dependent transporter, TA: toxin-antitoxin, ACP: acyl carrier protein, AR: ACP reductase, AG: ATP-grasp domain, AT: acetyltransferase).

Although no complete MiBIG BGCs showed similarity to the Halo_AMP BGCs detected, antiSMASH 6 now performs a gene-by-gene comparison of query BGCs to the MiBIG database, which provides another way to infer the possible function of the encoded proteins. Genes coding for proteins with A-domains returned frequent hits (11 of the 20 CC representatives) (Table S4), and the most common best hits include BagE encoded in the BGC responsible for the production of the trans-coumaric-acid-derived antibiotics bagremycins (106), and YtkN encoded in the yatakemycin BGC, a spirocyclopropane antibiotic (107). Both these genes are proposed to encode enzymes that function as a phenylacetate-CoA ligase (PCL), which carry out adenylation of the aryl/acyl acid for either direct amide bond formation, or through a CoA-linked thioester intermediate (107). Another frequently detected hit was BrtI, encoded in the BGC responsible for the production of the halogenated glycolipids bartolosides, with functions reported as a part of a glycolipid exporter cassette as well as a transcriptional regulator (108–110). The genes returning the most frequent hits are those encoding FDHs (in 12 of the 20 CCs). The hits include the *orf12*-encoded FDH in the BGC responsible for producing the chloroanthrabenzoxocinones antibiotics zunyimicins (111,112), and most frequently the pyrrole brominase Bmp2 encoded in the *bmp* BCG responsible for producing pentabromopseudilin and associated polybrominated aromatic organic compounds (37). Finally, 4 of the 20 representative BGCs (CCs A, E, H, Q, Table S5) were also predicted, by the new antiSMASH v7 (beta) (80), to encode regulators for antibiotic production, which further supports that these sequences are linked to the production of bioactive NPs.

Although more work is undoubtedly needed to characterize these BGCs, elucidate their ecological roles, and link them to their respective output molecules, we believe the Halo_AMP pseudoclass proposed here comprises a valuable addition to the diverse repertoire of sponge-holobiont-encoded NPs.

### CC A: a putative link to barettin?

Despite the general lack of similarity to characterized BGCs seen in this dataset, BGCs represented by CC A showed similarity to the known *bmp* cluster. Namely with CC A’s FDH showing homology to the pyrrole brominase *bmp2*, with both clusters featuring P450 enzymes, an adenylation domain, and additional oxidoreductases (37). Additionally, CC A features antibiotic production regulatory elements (80), indicating potential involvement in the production of a bioactive specialized metabolite.

Additionally, we observed that CC A features two BGC variants (Fig.1.b): a 1xP450 variant (CC A.1) present in 8 host sponge species, and a 3xP450 variant (CC A.2) present exclusively in *G. barretti* NOR samples. BGCs detected in metagenomic assemblies are often found in truncated contigs and are therefore incomplete, and repeat regions are noteworthy for being difficult to correctly assemble. In order to gain proof of the existence of these two variants, we searched in the reassembled versions of the recovered MAGs for longer contigs and thus more complete versions of the BGCs. This strategy showed some success, with 11 cases of BGCs that were either found on longer contigs or had not been assembled in the original metagenome assembly at all. A CORASON-based phylogeny of this set of BGCs (Fig.2) showed that this type of BCG is still challenging to assemble, with several extremely truncated examples (Fig.2). Nevertheless, we can observe that this BGC architecture features a set of core genes (A, FDH, rSAM, P450) with largely conserved regions both up- and downstream of this core, as well a consistent and exclusive detection of the 3xP450 variant in *G. barretti* NOR samples

**Figure 2.**
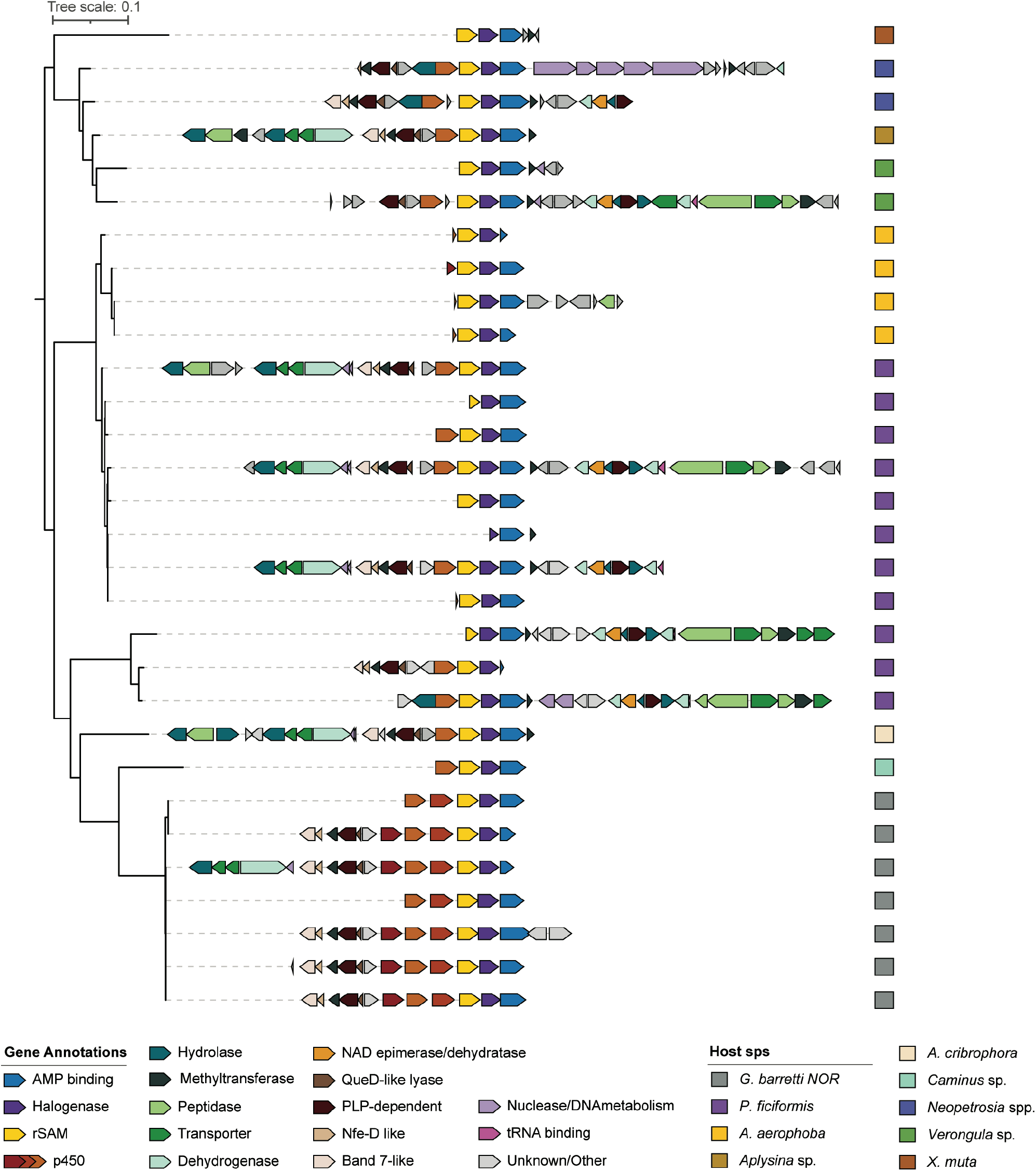
Characterization of CC A BGCs. CORASON generated phylogeny of CC A BGCs. Genes are colored based on predicted function, and each BGC is annotated by host sponge species origin.

*G. barretti* NOR (Norway) samples were captured up to approximately 1000 m depth, in contrast to *G. barretti* CAN (Canada) samples which originate from depths below 1000 m (Table S1). The presence of this *G. barretti* NOR-specific variant is of particular interest since it has been recently shown that the presence of several *G. barretti*-derived bioactive compounds is detected exclusively in individuals shallower than 1000 m (42). More specifically, these depth-dependent metabolites comprise the well-known brominated diketopiperazine (DKP) barettins, as well as the cyclo Pro-Arg DKP. The majority of non-DKP peptides and indole-derived NPs also isolated from *G. barretti* do not display this depth dependence (42). Furthermore, the same study has linked Acidobacteriota to the production of the majority of the DKP tryptophan derivatives detected, and indicated several acidobacteriotal ASVs detected in *G. barretti* showing strong positive correlations with the abundance of these metabolites (42). Similarly, BGCs in CC A are derived from MAGs with predicted acidobacteriotal origin, namely belonging to the genus Bin61 (Fig.1, Table S3). The phylogenetic diversity of the Acidobacteriota Bin61 MAGs recovered here showed a highly conserved species-level clade (MAG ANI > 99%) in *G. barretti* NOR (Fig. S1, Fig. S2). The 16S rRNA gene recovered from the representative MAG of this clade also returned a perfect match (100% coverage and identity, bit-score: 127, E-value: 6e-31) with one of the ASVs mentioned in the study above, ASV 144. Interestingly, all BGCs detected in the Acidobacteriota Bin61 MAGs recovered fall within CC A, with one exception, and there is a clear phylogenetic separation of Acidobacteriota Bin61 MAGs that encode Halo_AMP BGCs and those that do not (Supplemental Fig.1). Furthermore, as this separation does not seem to be correlated with predicted abundance of MAGs in host samples or with the quality of the MAG, this is thus a potential indication of the evolutionary moment where this BGC is gained/lost.

In this way, we believe that CC A provides a promising lead to decipher the biosynthetic origin of the barettins in *G. barretti*, as well as additional brominated peptidic NPs in several other host sponge species. We have thus decided to focus on the bioinformatic characterization of this CC for the remaining of this study.

### CC A’s core enzymes show high levels of sequence divergence from databases

To gain further insight into the potential roles of the BCGs in CC A, we set out to further characterized CC A’s core enzymes by leveraging specific annotation tools and sequence data of characterized members of each enzyme class obtained from publicly available databases. Given their central character in the context of this CC, rSAM and P450 encoding genes are also considered to be core genes, in addition to those encoding for an A-domain and FDH.

Analysis of the CC A’s adenylation domain active site specificity-conferring residues (AdenylPred (81), Table S6) predicted an aryl-CoA ligase functional class with substrate specificity for cinnamate and succinylbenzoate derivatives. However, prediction scores are low, undoubtedly a consequence of the low number of sponge-derived sequences in public databases and training datasets for prediction tools, as well as inherent broad substrate specificities of this class of enzymes (81). In this way, although without providing further detail on the specificity of these A domains, this prediction supports the expectation that these enzymes adenylate aromatic/phenolic amino acids. Additionally, active site specificity-conferring Stachelhaus residues and extended active site residues (PARAS (82)) within CC A A-domains showed extreme conservation, with an average pairwise identity of 100% for Stachelhaus residues and 96% for extended active site residues (Table S7), an indication of conserved substrate specificity within this CC (113).

To gain more insight into the relationship of CC A’s FDHs with known FDHs, and consequently into their substrate specificity, we generated a sequence similarity network (SSN) consisting of all FDHs recovered here, and a curated list of characterized FDH sequences including those compiled by Fisher et al. (65) as well as additional phenol and pyrrole halogenases (Fig 3.a, Table S2). We observed very limited clustering of Halo_AMP FDHs with those from the Fisher et al. (65) dataset, and no clustering with the additional curated sequences (Fig3.a). CC A’s FDHs clustered mostly with Halo_AMP FDHs, and with a small number of FDHs that belong to the larger phenol halogenase subnetwork described in Fisher et al. (65). Seeing that the barettins are brominated in the indole benzene/phenol ring, this placement does not exclude that CC A’s FDHs can act on tryptophan/indole substrates, and consequently does not invalidate the link between CC A and the barettins.

**Figure 3.**
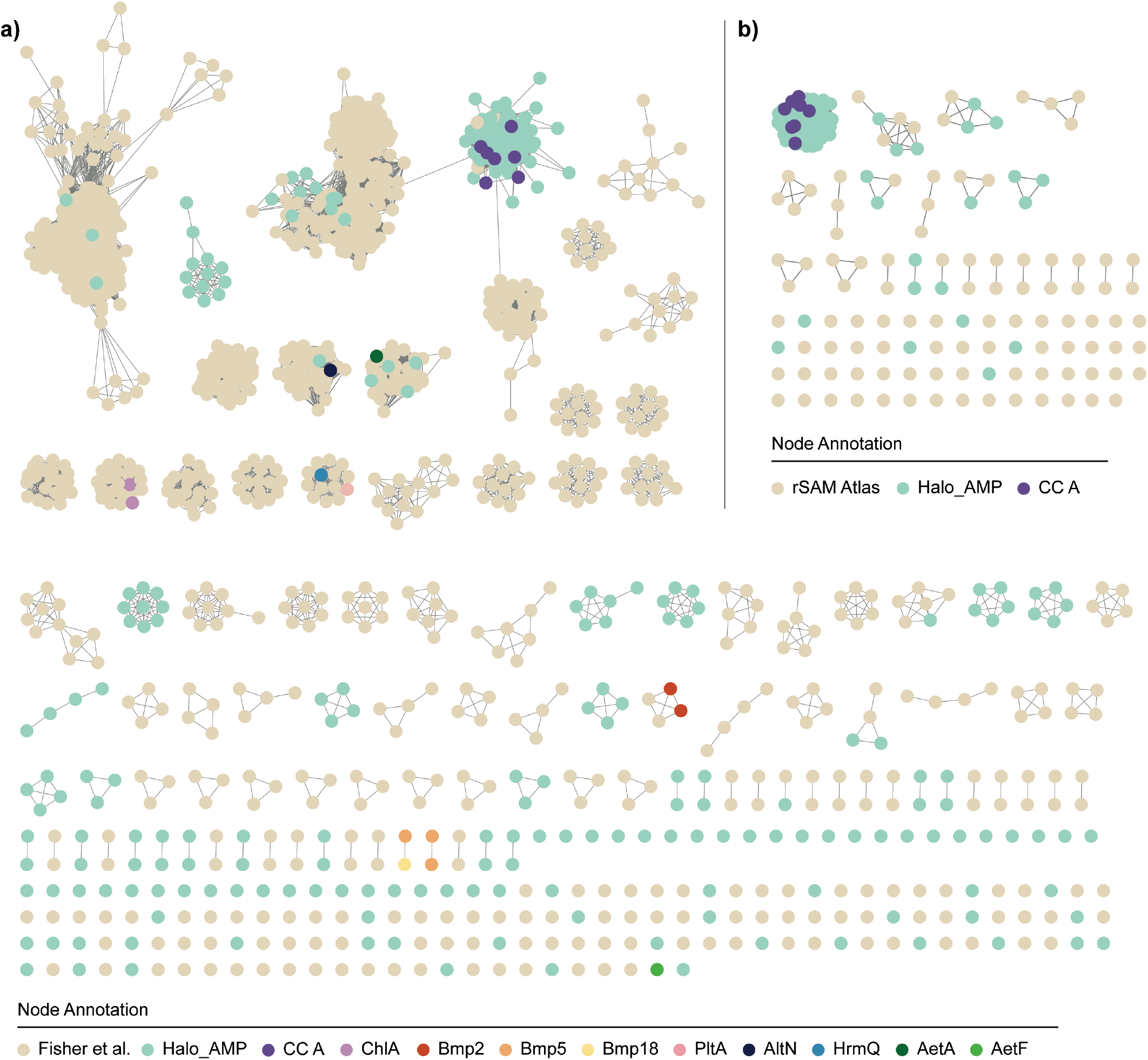
Sequence diversity of recovered enzymes and their characterized counterparts. a) Sequence similarity network of FDHs recovered in this study, those derived from Fisher et al. (65), and a curated set of characterized FDHs (Table S2). b) Sequence similarity network of rSAMs recovered here and representative characterized sequences derived from Holiday et al. (85). Nodes are colored by study origin and protein name.

In a showcase of how sequence similarity comparisons are poor predictors of FDH substrate specificity, we saw that a number of the additional characterized sequences also clustered poorly, or not at all, with Fisher et al.’s dataset. Particularly interesting here are the cases of the pyrrole brominase Bmp2, phenol brominases Bmp5 and Bmp18, and the unique indole/tryptophan brominase AetF (37,38,64). AetF halogenates the indole benzene ring in positions 5 and 7, and is the first example of a single-component (i.e. does not require a separate flavin reductase as a partner) flavin-dependent tryptophan halogenase, an enzymatic strategy that would fit CC A’s FDH role in the production of barettins (64). AetA, AetF’s sister FDH, halogenates the pyrrole ring of the tryptophan indole in positions 3 and 2, and clustered here in the subnetwork populated by Fisher et al.’s histidine halogenases (65). AltN, a tyrosine brominase, appeared in small halogenase subnetwork with and no other annotated sequences (61)). This indicates that the placement of a FDH sequence outside of Fisher et al.’s indole subnetwork does not exclude that this FHD acts on indole substrates. The pyrrole (PltA,HrmQ) and phenol (ChlA) chlorinases are placed in subnetworks of their own with sequences of each of their respective classes (60,114,115), which could indicate that characterizing brominating FDHs based on sequence similarity is more challenging than for chlorinating FDHs. It is known that FDHs tolerate a wide range of substrate scaffolds, and sponge-derived FDH sequences in particular show significant low identity to characterized halogenases, which can further point to diversity in chemical mechanism beyond substrate specificity (48,116). In yet another display of this diversity, even though a large fraction of marine NPs feature halotryptophans, only three Halo_AMP FDHs nodes (two are seawater-derived and one is a sponge-derived Fig 1 singleton) clustered with the largest Fisher et al. subnetwork, the indole halogenases (65,117). This illustrates how challenging it is to predict functionality and substrate specificity based on these enzymes’ protein sequence similarities.

An even more striking example of protein sequence dissimilarity between the enzyme-encoding genes detected here and those in characterized reference enzymes is seen for rSAM enzymes detected in CC A and Halo_AMP BGCs. The large majority of Halo_AMP rSAM sequences, including all of CC A’s rSAMs, did not cluster with any characterized rSAM Atlas homologs (Fig3.b) (85). rSAM enzymes are known for their remarkable ability to catalyze an extremely broad range of reactions, and for the poor correlation between sequence similarity-based clustering and functional specificity (85). Therefore, it is difficult to speculate about the putative function of these enzymes based on this analysis and it showcases the dimension of novelty encoded in the marine sponge holobiont ecosystem.

Finally, we analyzed perhaps the most striking family within CC A’s core enzymes, the cytochrome P450s. These heme-binding oxygenase enzymes are ubiquitous in nature and display a remarkable diversity in substrate specificity and reaction chemistry (93). As it is the presence of two additional P450s in *G. barretti* NOR samples that sets their CC A BGCs apart from the variants observed in other samples, we performed an extensive characterization of their sequences: a placement in a global phylogeny of chemically characterized single-domain homologues, including the P450s present in the *bmp* BGC, as well as annotation with CYP family and KEGG Orthology (KO) (Table S9) (37,38,64,86,89,90). Nevertheless, functional characterization is challenging. As an example, Bmp7 is known to catalyze the coupling of two brominated phenolic rings, but its KO/CYP classifications remain shallow, at the CYP1 superfamily level (37,118).

We observed that the CC A’s P450s form a clade of their own, which shows the least similarity to any characterized P450 (Fig 4). Within this clade we can observe an interesting distribution of the sequences: P450s in CC A.1 formed a clade with one of the three CC A.2 P450s, while a separate clade is seen with the two remaining P450s from CC A.2. This hints towards conserved function of the shared P450 across all BGCs, and additional, distinct, functions of the two extra copies of this enzyme. Despite this distribution, all CC A P450s are functionally annotated as K21164 and CYP family CYP253, showcasing once more the challenge in functional prediction and annotation of these diverse enzymes. These K21164 annotations point towards a role as P450 hydroxylase, and P450s belonging to the family CYP253 have shown to participate, with unclear roles, in the biosynthesis of enediyne antibiotics, as well as microsclerodermins and pyrodomycin (119–123). The characterized homologues that are closest to CC A’s sequences are also known to function as P450 hydroxylases (K22997/8, K15907), including Bmp12 (38,124–126).

**Figure 4.**
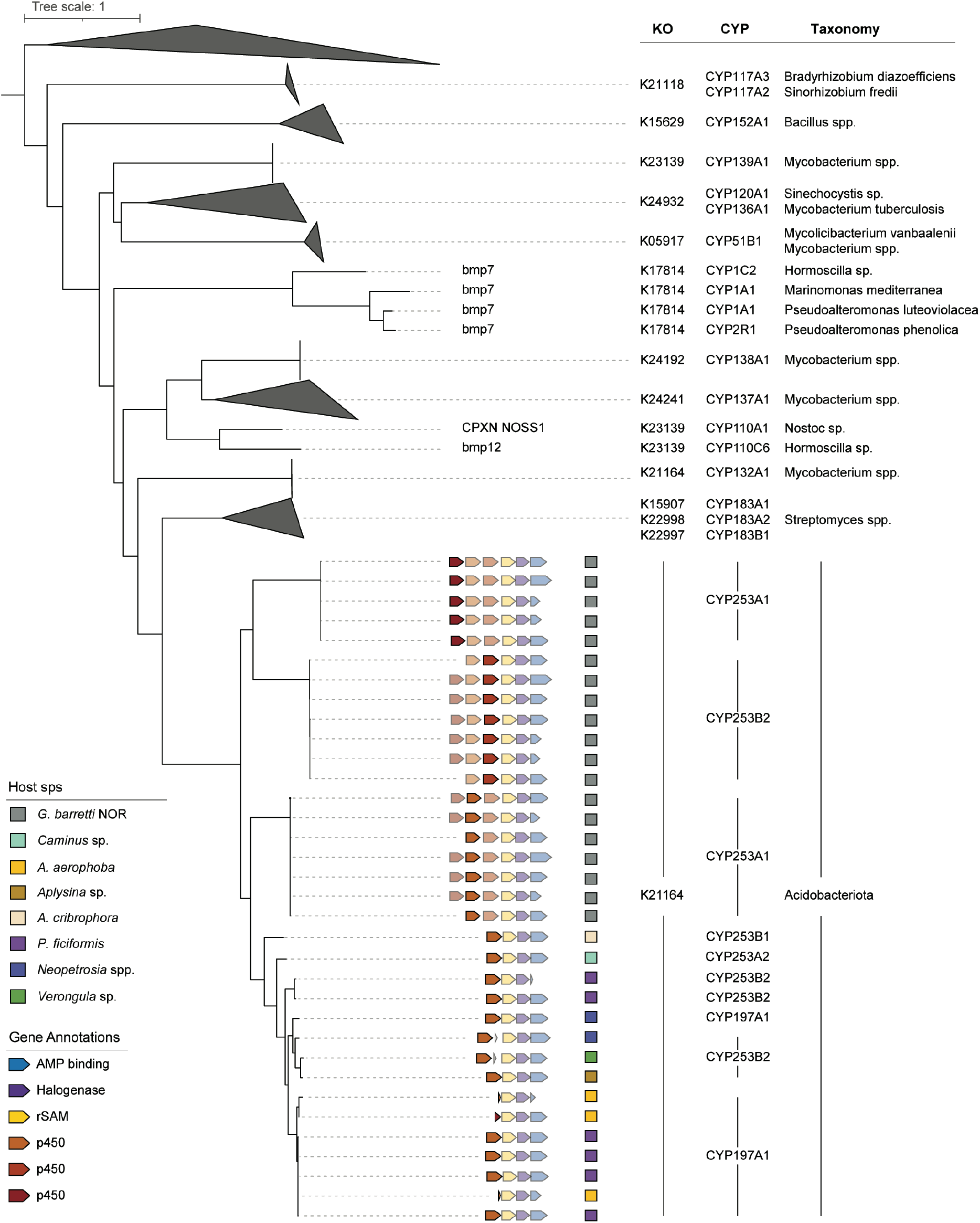
Evolutionary relationships between CC A cytochrome P450s and characterized P450s. Phylogenetic tree constructed with FastTree of P450 sequences recovered from CC A and single domain entries from the SwissProt database, rooted at midpoint. All sequences are annotated with predicted function (KO and CYP family), and bacterial host taxonomy (Table S9). For P450s identified here, the genomic context within their parent BGC is shown where genes are colored by predicted function. P450 encoding genes are coloured in bold in their respective leaf BGC representation. As such, 3xP450 variant BGCs appear in triplicate in the tree, with relevant P450 gene in bold in each leaf.

### The barettins remain orphan compounds

The exclusive presence of CC A.2 (3xP450 variant) in the ‘shallow’ *G. barretti* NOR, as well as in a MAG, which is strongly correlated with the presence of barettins, are indications that link CC A.2 to these metabolites. Barettin consists of a cyclo-Arg-Trp (cRW) diketopiperazine, brominated in position 5 of the dehydrogenated Trp residue. Additionally, most depth-correlated barettin derivatives feature further modifications to the Trp residue, such as hydroxylations and (de)saturations (42). In this way, computational predictions of the core A domains and FDHs encoded within the CC A.2 gene clusters indicate a phenolic substrate specificity that further support a link to these tryptophan-derived molecules.

However, a key element is still missing in the retro-biosynthesis hypothesis: the generation of the diketopiperazine (DKP) core. DKPs are cyclic dipeptides and derive from “head-to-tail” cyclization of two amino acids, with the two nitrogen atoms of the six-membered ring forming amide bonds (35). DKP scaffolds are known to be synthesized by two enzyme families: nonribosomal peptide synthetases (NRPSs), characterized mostly in fungi, and cyclodipeptide synthases (CDPSs), which are more often seen in bacterial gene clusters (51,127–130). Considering the absence of canonical NRPSs in the metagenome of *G. barretti* NOR, it is likely that if barettin is produced by a sponge symbiont, a CDPS would be involved in its biosynthesis. CDPSs feature a structurally conserved Rossman-fold domain and use aminoacyl-tRNAs for the formation of the two peptide bonds of cyclodipeptide products via a covalent acyl-enzyme intermediate (131–133). This is a poorly characterized enzyme family, with most characterized bacterial CDPSs originating from Actinobacteriota, Proteobacteria and Firmicutes and sharing low amino acid sequence identity across the family (average of 30%) (131,134,135). Characterized CDPSs that install Arg and Trp residues are particularly scarce, with a single characterized cRW CDPS (AvaA) recently discovered from the fungus *Aspergillus versicolor* (52,130,136). However, no CDPS was detected in Halo_AMP BGCs when searching for both the PFAM CDPS PF16715 pHMM, as well as the CDPS sequences newly reported by Yee et al. (52). Nevertheless, it is plausible that the sequence identity of a putative CC A acidobacterial CDPS is too low to currently detect via sequence similarity methods. In a clear showcase of novelty, Yee et al. have identified an entire family of novel CDPSs which were previously annotated as hypothetical proteins and showed no sequence similarity to fungal DKP-producing NRPSs or bacterial CDPSs (52). Whether this is a result of a limitation of the sequence similarity searching algorithms, or whether these enzymes constitute a structurally novel family that has evolved independently/convergently, has yet to be elucidated, as no structural data is available for these novel CDPSs.

Furthermore, we have seen that CC A.1 (1xP450 variant) is widespread across host sponge phylogeny, including sponge species for which cRW DKPs have not been reported. As such, CC A.1 is unlikely to be involved in the production of these molecules. Consequently, since what distinguishes CC A.1 from CC A.2 is the presence of the additional two copies of the P450 enzyme, we must consider either the possibility that there may be a new strategy being employed here to generate the DKP involving these additional P450s, or that CC A.2 is not at all responsible for producing barettins. To confirm either hypothesis, it is ultimately necessary to carry out experimental work to functionally characterize these BGCs, ideally through heterologous expression of the BGC as well as individual enzymes which can then be assessed *in vivo* and/or *in vitro* for both enzymatic activity and structural elucidation.

## Conclusion

Marine sponge holobionts continue to be a promising source of novel biosynthetic pathways of unique halogenated NPs. Here we have contributed to the expansion of this repertoire by exploring BGC architectures that fall outside the canonical biosynthetic classes, which has proven to be a valuable strategy to access otherwise hidden diversity. We show that the presence of putative BGCs coding for the production of peptidic halogenated NPs is widespread across host sponge taxonomy and geographical location, and highlight Acidobacteriota as important community members harboring these BGCs. Furthermore, we identified a particularly interesting BGC architecture that might be related to the production of the known DKP barettins, given its enzymatic characteristics and predicted taxonomic origin. There is, nevertheless, an ultimate need to pair these bioinformatic predictions with experimental work to functionally characterize these enzymes and eventually link BGCs to chemical products.

## Acknowledgments

The authors wish to thank Ellen Kenchington for the Canadian *G. barretti* samples, the late Hans Tore Rapp for his repeated help to sample the Norwegian fjord *G. barretti* specimens, Adriaan Schrier for offering his submarine to collect sponges in Dominica, and dr. Vasilis Gerovasileiou and HCMR (Hellenic Centre for Marine Research) for the collaboration in sampling the *P. ficiformis* samples.

This research was financially supported by the VLAG NWO PhD project “*Mare incognita*” to CL, the European Commission through Horizon2020 project SponGES (Grant agreement ID: 679849) to DS, and AG and the Marie Sklodowska-Curie Individual Fellowship COSMos (Grant agreement ID: 897121) to MAS.

## Author Contributions

C.L: Research design. eDNA extraction, QC-filtering, assembly, binning. BGC identification and characterization. Downstream data analysis. Writing the manuscript.

M.A.S: Research design. DOM samples eDNA extraction, QC-filtering, assembly.

M.A: p450 sequence MSA, phylogenetic tree building and annotation (KO, CYP).

B.K, J.L: Research design, preliminary results.

J.vdO: Research design, manuscript reviewing/editing

M.H.M: Research design, manuscript reviewing/editing

D.S: Research design, manuscript reviewing/editing

## Data Availability

The data for this study have been deposited in the European Nucleotide Archive (ENA) at EMBL-EBI under accession numbers PRJEB51534 and PRJEB59408. Python scripts created for this analysis are available at https://github.com/CatarinaCarolina/sponge_halo_amp.

