## Supplementary Data for "Elucidating biosynthetic pathways related to the synthesis of small halogenated peptidic natural products in marine sponge microbiomes"

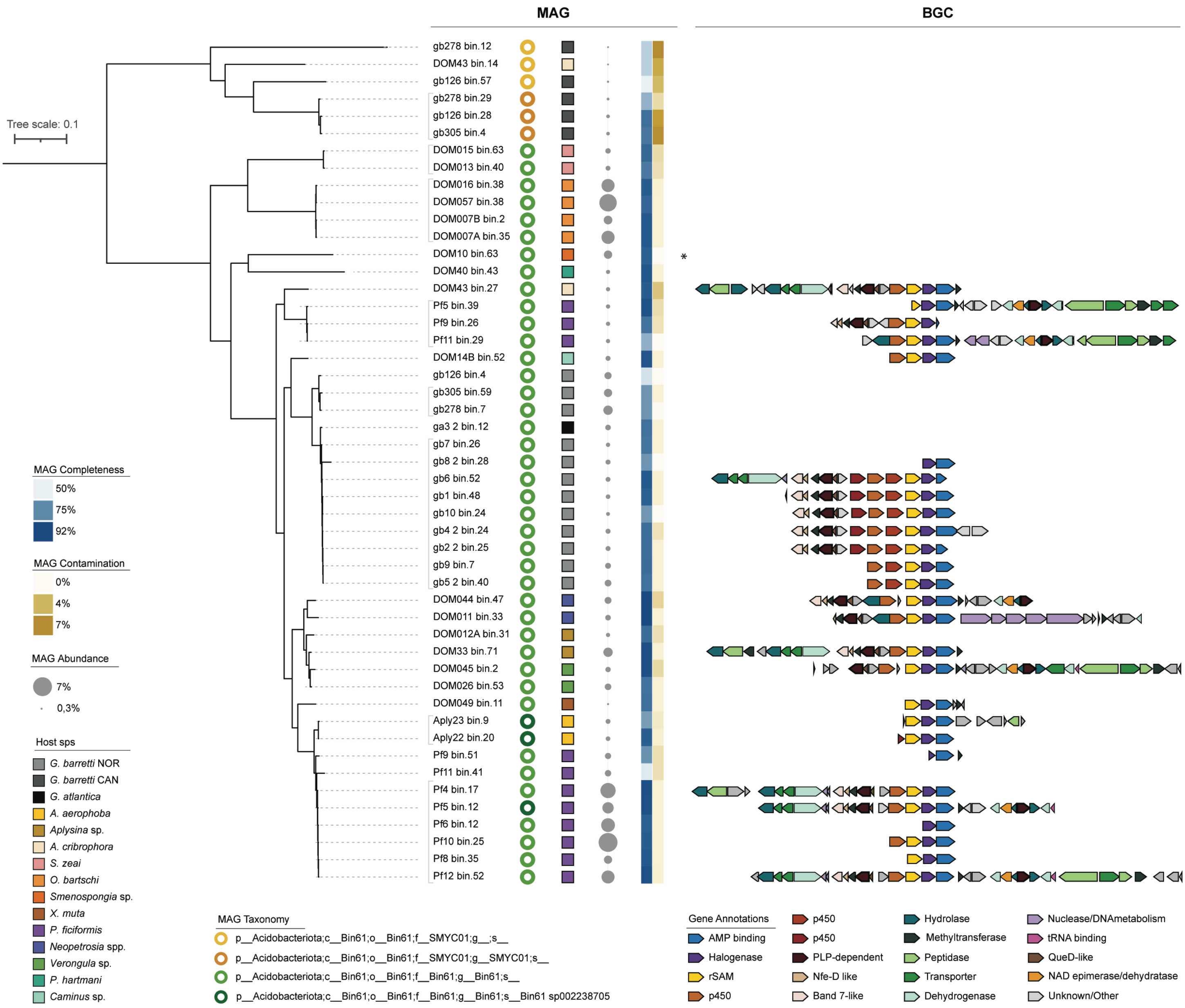

**Figure S1.** Diversity and biosynthetic content of Acidobacteriota Bin61. Cladogram based on GTDB classification of Acidobacteriota Bin61 MAGs recovered in this study, rooted at midpoint. MAGs are annotated with GTDB-tk predicted taxonomy, host sponge taxonomy, predicted abundance in host sponge sample, MAG completeness and contamination, and the encoded CC A BGC (*: DOM10_bin.63 encodes a HALO_AMP BGC not included in CC A). BGC genes are colored based on predicted function. Grey brackets encompass MAGs for which the average nucleotide identity (ANI) is above 99%.

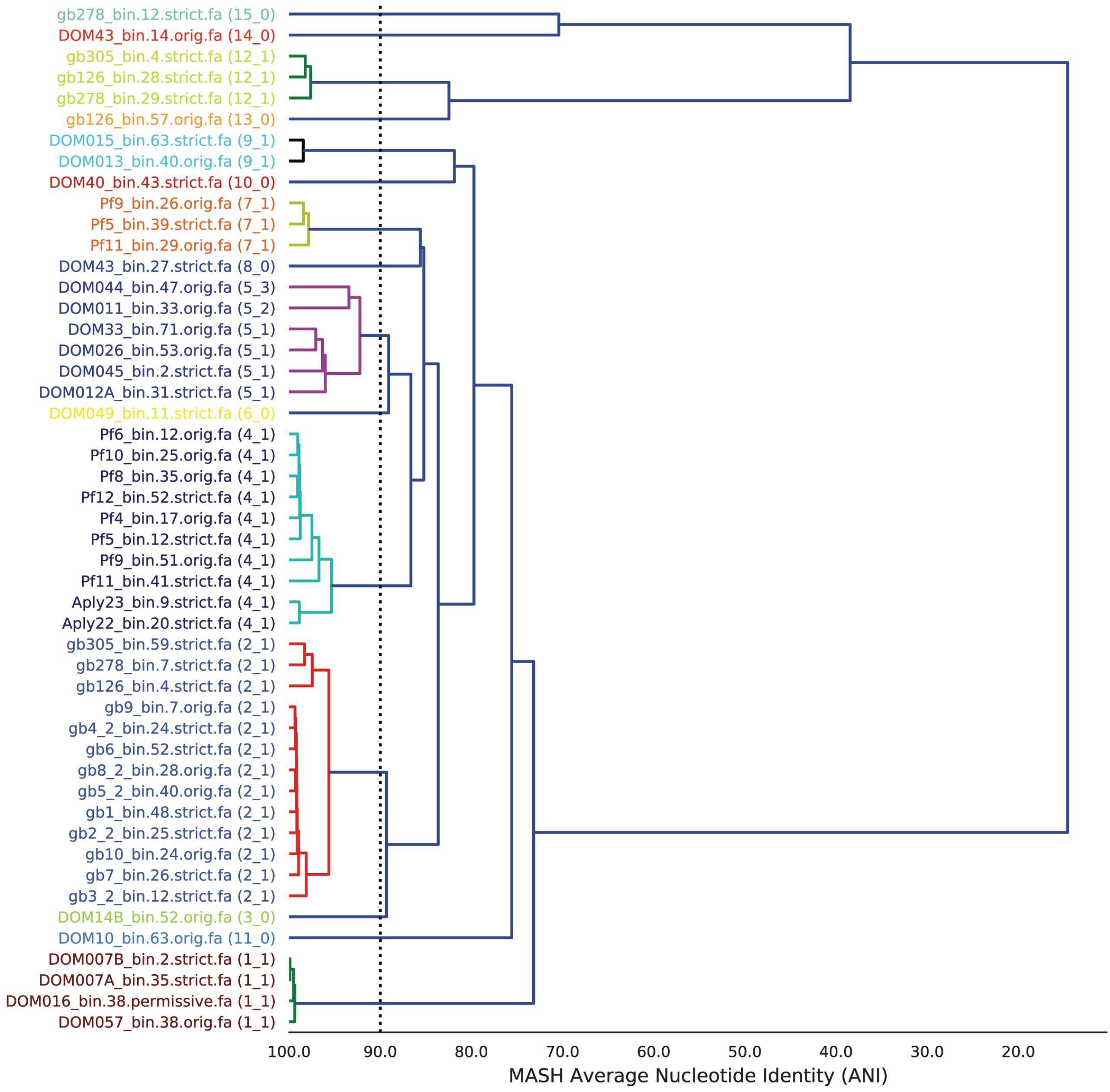

**Figure S2.** dREP clustering of all MAGs recovered and taxonomically classified as Acidobacteria Bin61**.**

**Table S1.** Sample Metadata

| Name | Type | Species | Depth (m) | Location | Coordinates | Country | Sampling Year | Sequencing Year | Sequencing technology |
|---|---|---|---|---|---|---|---|---|---|
| **Aply16** | Sponge Tissue | Aplysina aerophoba | 10,25 | Cala Montgo | N42°06'52.20", E3°10'06.52" | Spain | 2014 | 2014 | Hybrid Illumina Pacbio |
| **Aply21** | Sponge Tissue | Aplysina aerophoba | 10,25 | Cala Montgo | N42°06'52.20", E3°10'06.52" | Spain | 2014 | 2014 | Hybrid Illumina Pacbio |
| **Aply22** | Sponge Tissue | Aplysina aerophoba | 10,25 | Cala Montgo | N42°06'52.20", E3°10'06.52" | Spain | 2014 | 2014 | Hybrid Illumina Pacbio |
| **Aply23** | Sponge Tissue | Aplysina aerophoba | 10,25 | Cala Montgo | N42°06'52.20", E3°10'06.52" | Spain | 2014 | 2014 | Hybrid Illumina Pacbio |
| **Pf4** | Sponge Tissue | Petrocia ficiformis | 5,5 | Sfakia, Crete (Semi-dark zone) | N35°12'0.7", E24°7'09.8" | Greece | 2018 | 2018 | Illumina |
| **Pf5** | Sponge Tissue | Petrocia ficiformis | 5,5 | Sfakia, Crete (Semi-dark zone) | N35°12'0.7", E24°7'09.8" | Greece | 2018 | 2018 | Illumina |
| **Pf6** | Sponge Tissue | Petrocia ficiformis | 5,5 | Sfakia, Crete (Semi-dark zone) | N35°12'0.7", E24°7'09.8" | Greece | 2018 | 2018 | Illumina |
| **Pf7** | Sponge Tissue | Petrocia ficiformis | 5,5 | Sfakia, Crete (Dark zone) | N35°12'0.7", E24°7'09.8" | Greece | 2018 | 2018 | Illumina |
| **Pf8** | Sponge Tissue | Petrocia ficiformis | 5,5 | Sfakia, Crete (Dark zone) | N35°12'0.7", E24°7'09.8" | Greece | 2018 | 2018 | Illumina |
| **Pf9** | Sponge Tissue | Petrocia ficiformis | 5,5 | Sfakia, Crete (Dark zone) | N35°12'0.7", E24°7'09.8" | Greece | 2018 | 2018 | Illumina |
| **Pf10** | Sponge Tissue | Petrocia ficiformis | 5,5 | Sfakia, Crete (Entrance) | N35°12'0.7", E24°7'09.8" | Greece | 2018 | 2018 | Illumina |
| **Pf11** | Sponge Tissue | Petrocia ficiformis | 5,5 | Sfakia, Crete (Entrance) | N35°12'0.7", E24°7'09.8" | Greece | 2018 | 2018 | Illumina |
| **Pf12** | Sponge Tissue | Petrocia ficiformis | 5,5 | Sfakia, Crete (Entrance) | N35°12'0.7", E24°7'09.8" | Greece | 2018 | 2018 | Illumina |

| | | | | | | | | | |
|---|---|---|---|---|---|---|---|---|---|
| **ga3** | Sponge Tissue | Geodia atlantica | 150 | Scengsbukt-Korsfjord | N60°8'8", E5°6'42" | Norway | 2017 | 2018 | Illumina |
| **gb1** | Sponge Tissue | Geodia barretti | 450 | Scengsbukt-Korsfjord | N60°8'8", E5°6'42" | Norway | 2017 | 2018 | Illumina |
| **gb2_2** | Sponge Tissue | Geodia barretti | 150 | Scengsbukt-Korsfjord | N60°8'8", E5°6'42" | Norway | 2017 | 2018 | Illumina |
| **gb4_2** | Sponge Tissue | Geodia barretti | 150 | Scengsbukt-Korsfjord | N60°8'8", E5°6'42" | Norway | 2017 | 2018 | Illumina |
| **gb5_2** | Sponge Tissue | Geodia barretti | 150 | Scengsbukt-Korsfjord | N60°8'8", E5°6'42" | Norway | 2017 | 2018 | Illumina |
| **gb6** | Sponge Tissue | Geodia barretti | 150 | Scengsbukt-Korsfjord | N60°8'8", E5°6'42" | Norway | 2017 | 2018 | Illumina |
| **gb7** | Sponge Tissue | Geodia barretti | 450 | Scengsbukt-Korsfjord | N60°8'8", E5°6'42" | Norway | 2017 | 2018 | Illumina |
| **gb8_2** | Sponge Tissue | Geodia barretti | 450 | Scengsbukt-Korsfjord | N60°8'8", E5°6'42" | Norway | 2017 | 2018 | Illumina |
| **gb9** | Sponge Tissue | Geodia barretti | 450 | Scengsbukt-Korsfjord | N60°8'8", E5°6'42" | Norway | 2017 | 2018 | Illumina |
| **gb10** | Sponge Tissue | Geodia barretti | 450 | Scengsbukt-Korsfjord | N60°8'8", E5°6'42" | Norway | 2017 | 2018 | Illumina |
| **gb126** | Sponge Tissue | Geodia barretti | 1213 | Davis Strait | N62°52'15.1", W58°37'34.32" | Canada | 2015 | 2018 | Illumina |
| **gb278** | Sponge Tissue | Geodia barretti | 1335 | Davis Strait | N61°53'36.13", W60°7'57.612" | Canada | 2014 | 2018 | Illumina |
| **gb305** | Sponge Tissue | Geodia barretti | 1437 | Davis Strait | N62°31'6.24", W59°58'13.872" | Canada | 2014 | 2018 | Illumina |
| **gb1_f** | Filtered Sea Water | N.a. | 150 | Scengsbukt-Korsfjord | N60°8'8", E5°6'42" | Norway | 2017 | 2018 | Illumina |
| **gb2_f** | Filtered Sea Water | N.a. | 150 | Scengsbukt-Korsfjord | N60°8'8", E5°6'42" | Norway | 2017 | 2018 | Illumina |

| | | | | | | | | | |
|---|---|---|---|---|---|---|---|---|---|
| **gb3_f** | Filtered Sea Water | N.a. | 150 | Scengsbukt-Korsfjord | N60°8'8", E5°6'42" | Norway | 2017 | 2018 | Illumina |
| **gb5_6_f** | Filtered Sea Water | N.a. | 450 | Scengsbukt-Korsfjord | N60°8'8", E5°6'42" | Norway | 2017 | 2018 | Illumina |
| **gb9_f** | Filtered Sea Water | N.a. | 450 | Scengsbukt-Korsfjord | N60°8'8", E5°6'42" | Norway | 2017 | 2018 | Illumina |
| **gb10_f** | Filtered Sea Water | N.a. | 450 | Scengsbukt-Korsfjord | N60°8'8", E5°6'42" | Norway | 2017 | 2018 | Illumina |
| **sw_7** | Filtered Sea Water | N.a. | 10,25 | Cala Montgo | N42°06'52.20", E3°10'06.52" | Spain | 2014 | 2014 | Illumina |
| **sw_8** | Filtered Sea Water | N.a. | 10,25 | Cala Montgo | N42°06'52.20", E3°10'06.52" | Spain | 2014 | 2014 | Illumina |
| **sw_9** | Filtered Sea Water | N.a. | 10,25 | Cala Montgo | N42°06'52.20", E3°10'06.52" | Spain | 2014 | 2014 | Illumina |
| **DOM10** | Sponge Tissue | Smenospongia sp. | 101 | Portsmouth | N15°34’33.24”, W61°27’20.16” | Dominica | 2016 | 2020 | Illumina Novaseq6000 and Oxford Nanopore MinION |
| **DOM14B** | Sponge Tissue | Caminus sp. | 230 | Portsmouth | N15°34’33.24”, W61°27’20.16” | Dominica | 2016 | 2020 | Illumina Novaseq6000 and Oxford Nanopore MinION |
| **DOM33** | Sponge Tissue | Aplysina sp. | 106 | Cottage | N15°36’54.66”, W61°27’48.93” | Dominica | 2016 | 2020 | Illumina Novaseq6000 and Oxford Nanopore MinION |
| **DOM43** | Sponge Tissue | Aciculites cribrophora | 213 | Connor Bay | N15°38'14.13", W61°27'39.13" | Dominica | 2017 | 2020 | Illumina Novaseq6000 and Oxford Nanopore MinION |
| **DOM40** | Sponge Tissue | Petrosia hartmani | 146,304 | Connor bay | N15°38'14.13", W61°27'39.13" | Dominica | 2016 | 2022 | Illumina Novaseq6000 and Oxford Nanopore MinION |
| **DOM005** | Sponge Tissue | Siphonodictyon coralliphagum | 137 | Portsmouth | N15°34’33.24”, W61°27’20.16” | Dominica | 2016 | 2022 | Illumina Novaseq6000 |

| | | | | | | | | | |
|---|---|---|---|---|---|---|---|---|---|
| **DOM007A** | Sponge Tissue | Oceanapia bartschi | 82 | Portsmouth | N15°34’33.24”, W61°27’20.16” | Dominica | 2016 | 2022 | Illumina Novaseq6001 |
| **DOM007B** | Sponge Tissue | Oceanapia bartschi | 82 | Portsmouth | N15°34’33.24”, W61°27’20.16” | Dominica | 2016 | 2022 | Illumina Novaseq6002 |
| **DOM011** | Sponge Tissue | Neopetrosia spp. | 134 | Portsmouth | N15°34’33.24”, W61°27’20.16” | Dominica | 2016 | 2022 | Illumina Novaseq6003 |
| **DOM012A** | Sponge Tissue | Aplysina sp. | 134 | Portsmouth | N15°34’33.24”, W61°27’20.16” | Dominica | 2016 | 2022 | Illumina Novaseq6004 |
| **DOM013** | Sponge Tissue | Svenzea zeai | 223 | Portsmouth | N15°34’33.24”, W61°27’20.16” | Dominica | 2016 | 2022 | Illumina Novaseq6005 |
| **DOM015** | Sponge Tissue | Svenzea zeai | 107 | Portsmouth | N15°34’33.24”, W61°27’20.16” | Dominica | 2016 | 2022 | Illumina Novaseq6006 |
| **DOM016** | Sponge Tissue | Oceanapia bartschi | 108 | Portsmouth | N15°34’33.24”, W61°27’20.16” | Dominica | 2016 | 2022 | Illumina Novaseq6007 |
| **DOM026** | Sponge Tissue | Verongula sp | 144,78 | Cottage | N15°36’54.66”, W61°27’48.93” | Dominica | 2016 | 2022 | Illumina Novaseq6008 |
| **DOM044** | Sponge Tissue | Neopetrosia spp. | 144 | Connor bay | N15°38'14.13", W61°27'39.13" | Dominica | 2016 | 2022 | Illumina Novaseq6009 |
| **DOM045** | Sponge Tissue | Verongula sp | 149,352 | Connor bay | N15°38'14.13", W61°27'39.13" | Dominica | 2016 | 2022 | Illumina Novaseq6010 |
| **DOM049** | Sponge Tissue | Xestospongia sp. | 140 | Connor bay | N15°38'14.13", W61°27'39.13" | Dominica | 2016 | 2022 | Illumina Novaseq6011 |
| **DOM057** | Sponge Tissue | Oceanapia bartschi | 113 | Connor bay | N15°38'14.13", W61°27'39.13" | Dominica | 2016 | 2022 | Illumina Novaseq6012 |

**Table S2.** Nucleotide sequence of the additional characterized FDHs used to supplement Fisher et al.'s dataset**.**

>MARME_bmp2

MNTNANYDVVIIGSGPAGSLCGIECRKKGLSVLCIEKDEFPRFHIGESLTGNAGQIITDLGLYDKMEEAEFPNKLG
VNVIGSLSKNEFFIPILAKTWQVRRSSFDKMLKEKALEHGVEYQTGMVTDVLRDGEKVVGATYKTAAGSQNVN
SKVLVDASGQTTFLSRKGVAGKRKVEFFSQQIASFAHFENVERDLPPFSTNTTILYSKQYHWSWIIPISPTTDSLGV
VIPKDLYYKECNSPEAAIEWGMNNISPELKRRFKNAEQVEASQSMADFSYRIEPFVGDGWLCIGDSHRFLDPIFS
YGVSFGMKEGIRSAEAIATSIQTGDWKTPFYAFRDWSNKGQQIAADLIRYFWIYPIFFGYQMQNPDLRDEVIRLL
GGCCFDCEGWKAPSIFSNAIKEYDRKQMETKIAS*

>MARME_bmp5

MKKRIAIIGAGLSGIAAIKQLTDEGHHVVCYEKAESFGGVFAAKKIYEDLHLTISNYFMAYSDFLPTEQSLKFWSKQ
EYVQYLKRYLAHFDIEKHIVYNHKVVNAEQNGDKWTVKVQSGSGEETESEFDMVVVCSGHFQEPKTPDLEGLS
DFMGDIIHSNDYRDKMAFKGKRVMCVGLGESSADITSEISEVAEKCILSLRRYPAVAPRYMAFQEDPYFTIDTSW
LTSRIVNKLPFSYHRGITKNIFHKYVNSRNLHLRIRGEWLHKSGPSIHQAVTKNERLFKPIAEGKVLPNIGGIERFEG
NTVIFKDGTHEEIDAIVFCTGYKLSFPFLQHKIECMRDLYKQIFIPSVGSSLAFVGFVRPQQGGIPVIAEMQSRYLA
QLASGVKSLPSLEKQKEVIMEDANHWETEYHITPHVASLVNYCHYMDSMARLVGCMPKTPSLLKDPLLRVKLLH
NPQFAAQYRLEGPHPMSESSRDFLVNFPNISTWPRIIHFECALAMQKLLSFLSMDNLRELKK*

>bmp5_PLPH

MRKKIAVIGAGLSGIAAIKQLTDGGHEVVCFEKAESFGGVFADKKIYEDLHLTISNYFMAYSDYVPSQQKLKFWS
KKEYVNYLGEYLARFELGQYIHYDHEVRKVEKQAGKWQVTTKHGMAEQTDTFDMVAVCSGHFQKPKMPELA
GLDMFEGEIEHSNDYRDKHKYAGKRVLCVGLGESSADITSEISEVASKCILSLRRYPAVAPRYMAFQEDPYFTIDTS
WLTSRIVNKLPHRYHGGITKGIFTKYVNSRNDHVRIRGEWLKKSGPSHHQAVTKNERLFRPIADGKVVPNIGGIV
RFEKNAVVFQDGTREEIDAVVFCTGYQLSFPFLDVSISNMRDLYKQMFIPEMGDSLSFIGFVRPQQGGIPVIAEM
QCRYLSQLASGEKSLPPLSEMVDIIKYDTEHWQTEYKITPHVASLVNYCHYMDSMAKLVGCMPEIPSLFKDPLLR
VKLLHNPQFAAQYRLDGPNKMTHTARSFLLGFPNISSWPRIIHFELALITQKLLSRLRLDGLRELSK*

>PL2TA16_bmp5

MNKTIAVIGAGLSGIAAVKQLTDGGHQVTCFEKAESFGGVFADKKIYDDLHLTISNYFMAYSDYVPNHQKLKFW
SKKEYINYLGEYIERFDIAKHIHYDHEVCCVQKQGDKWLVTYKNADTEQTKEFDMVAVCSGHFQKPKLPDLPGL
DMYQGNIEHSNDYRDKHNYAGKRVLCVGLGESSADITSEISQVARKCILSLRRYPAVAPRYMAFQEDPYFTIDTS
WLTSRIVNKLPHRYHGGITKGIFNKYVTSRNDHVRIRGEWLKKSGPSHHQAVTKNERLFRPIADGKVTPNIGGIE
RFEKNAVVFKDGTREEIDAVVFCTGYQLSFPFLDVSIANMRDLYKQMFIPEMGHSLSFIGFVRPQQGGIPVIAEM
QCRYLSKLASGEAQLPTLSEMHDVIKYDTKHWQTEYKITPHVASLVNYCHYMDSVAKLVGCMPQIPSLFKDPML
RVKLLHNPQFAAQYRLDGPNNMTHTAREFLLSFPNISSWPRIIHFEVALAAQKLLSRLRLDGLREISK*

>PL2TA16_bmp2

MSEFKSYDVVIIGSGPAGSLCGIECRKKGLSVLCIEKDEFPRFHIGESLTGNAGQIIRDLGLAEEMDAAGFPDKPGV
NVIGSLSKNEFFIPILAPTWQVKRSDFDNMLKRKALEHGVEYQQGLVKDVIKHDGKVVGAIYKADDMEHQVRS
KVLVDASGQNTFLSRKGIAGKREIEFFSQQIASFAHYKGVERDLPPFSTNTTILYSKQYHWSWIIPISPDTDSLGIVI
PKDLYYKECKNPDDAIEWGMENISPEIRRRFQNAERIGDSQSMADFSYRIEPFVGDGWLCIGDAHRFLDPIFSYG
VSFAMKEGIRAADAIKQAIDGNDWKTPFYAYRDWSNGGQQIAADLIRYFWIYPIFFGYQMQNPDLRDEVIRLL
GGCCFDCEGWKAPTIFRNAIEEYDRKQMVG*

>PLSP_bmp2

MNGFTHYDVVIIGSGPAGSLCGIECRKKGLSVLCIEKEQFPRFHIGESLTGNAGQIIRDLGLAEDMDAAGFPDKP
GVNVIGSLSKNEFFIPILAPTWQVRRSDFDDMIKRKAVEHGVEYKLGMVTDVIKDGEKVVGALYKADGVEHQV
RSKVLVDASGQNTFLSRKGVAGKRQIEFFSQQIASFAHYKGVERDLPPFSTNTTILYSKQYHWSWIIPISPDTDSL
GVVIPKDLYYKECKNPDDAIAWGMDHISPELKRRFKNAERQGDSQSMADFSYRIEPFVGDGWMCIGDAHRFL
DPIFSYGVSFAMKEGIRAAEAIAQVVAGQDWKAPFYAYRDWSNGGQQIAADLIRYFWIYPIFFGYQMQNPDLR
DEVIRLLGGCCFDCEGWKAPAIFRNAIEEYDRKQMAS*

>PLPC_AltN

MAKENVDVLIIGAGPAGTMTAAKLIQAGLTVKIVERSHFPRYVIGESLLPQSMQHLEDAGFMDALVARNYQKKI
GANFKQDDFQEFFDFSKNYTDGWTWTWQVPRDDFDNVLAEEVQKMGAPIEFGTSVVDANMDLEEPIIKVVN
EAGDVQEIQAKFVVDSSGYGRVLANLLDLNEPSSAPTRATLFAQFKPADGIETGAENNKVTILVHEEDVWVWLI
PFSDGHVSVGFVGNPEYLESAEGDAKTRFLTYLARDQHASAMLDGLALCMEPKEIKGYSKGIKQLFGKNFVLTG
NASEFIDPVFSAGVMFAIESGARAADLIVAQLNGEEADWQQDYEAHMLKGIAVMRTYIEGWYDGRLRRLFFSD
NKPEKIKSQITSVLAGYVWDETNPFVNRSEKTLNLLSELHNNPEMA*

>GUM202_bmp5

MLDCIVIGAGSGGLVTTKELLEQGVGEVVCLEQAEDVGGVFTNTYDSLVLTSSATISMFSDFWIGDGNQHKFW
TKDEVVDYWKKYAEHFGVREHIRFGSKVVAVVEQGGEGWQVQLASGENLLTKRVALAIGNNAIPKYPEWKELL
TEVEYSHSQDYRNADRFAGKTVLAVGGGESGADIALEISRVASKCWVSLRNSAGWIVPRRRGINAADISTHRGV
WGLPRDYGAVLTEAVNQAELSQKDPVYDTVVKLNQKVEAKKGIWGIFATKNFSLPKAIVNHGCKVVGEIVKVE
DGGRTLHTADGECLTNVDAVVFSTGYKNAVSFLPEELKQTDPRGLYKHMFHPKYQDKLVWIGWARPNFGSQF
PIMEMQARLFALICKGELALPAPVEMEKVACIDRATYLEQFENNAHQVRSLVDYHRYIDDMASLIGCEPPLWQY
FFLHPRIWLRMVYGAIQSTQFRLRGPGNKESLARELLMKLPVSKPTHIVKAGLKGRVIYAFKALIPKIIFVGLEGSRK
SSSSLAVSRVSS*

>GUM202_bmp18

MLDCIVIGAGPGGLVCTKELIEQGLQEVVCLEQAKDAGGVFANAYDSLVLTSSATISMFSDFWIGDGNQHEFW
TKDEAVDYWKRYAEHFGVLERIRFNSKVVAVVEQGREGWQIQLESGDILLSKRLALATGNNAIPNYPEWKNLLT
HVEYFHSQEYRNADMFHDKNVLAVGGGESGADIALELSSVASQCWVSIRNTMGWVATRKRSLEGNVVAADV
ALNRLVWAMSGKTITKVIQQADLIEQDPVFDAAVELNKKIKGSNGIWGIFGTKNFSFPKAIVYHGCKVVGEIVKV
EDGGRTLHTAEGECLENVDAVVFSTGYKNAVSFLPEELKQTDPRSLYKHMFHPKYRDKIVWIGWARPNYGSQF
PVMEMQARLFALICKGELALPATAEMERVACIDRVANLEQFDHHAYRVRSLVDYHHYMDDMASLIGCKPSLW
KYLFSAPRICLPLVFAAIQGTQFRLQGPGNKESLARKILIKLPIIVPTPIVKGLLRQSLADALSLSRLGR*

**Table S3.** GCF metadata. Features each GCF's samples, host sponge species, MAGs and MAG predicted taxonomy.

| GCF | Samples | Host_taxa | dRep representative MAGs | MAG taxonomy |
|---|---|---|---|---|
| **1896** | Aply16,Aply22,DOM005,DOM007A,DOM007B,DOM012A,DOM015,DOM026,DOM044,Pf11 | Oceanapia bartschi,Neopetrosia sp.,Aplysina aerophoba,Svenzea zeai,Aplysina sp.,Petrocia ficiformis,Verongula sp.,Siphonodictyon coralliphagum | -,Aply23_bin.19.fa,Aply16_bin.2.fa,DOM005_bin.9.fa,DOM012A_bin.2.fa,DOM016_bin.19.fa,DOM044_bin.14.fa,Pf11_bin.65.fa | -,p__Poribacteria;g__WGA-3G;s__,p__Poribacteria;g__;s__ |
| **1** | Aply16,Aply22 | Aplysina aerophoba | Aply22_bin.11.fa | p__Poribacteria;g__MSPOR6;s__ |
| **2002** | Aply16,DOM013,DOM015,DOM044,DOM045,DOM10,DOM33,DOM40,DOM43,Pf12,Pf6,gb2_2,gb9 | Neopetrosia sp.,Petrosia hartmani,Aplysina aerophoba,Aciculites cribrophora,Svenzea zeai,Aplysina sp.,Petrocia ficiformis,Verongula sp.,Geodia barretti NOR,Smenospongia sp. | Aply16_bin.4.fa,-,DOM045_bin.26.fa | p__Acidobacteriota;g__;s__,- |
| **2249** | Aply16,Aply23,DOM14B,Pf10,Pf11,Pf6,Pf7,Pf8,Pf9 | Caminus sp.,Aplysina aerophoba,Petrocia ficiformis | -,Pf4_bin.17.fa,DOM14B_bin.52.fa | -,p__Acidobacteriota;g__Bin61;s__ |
| **5** | Aply16 | Aplysina aerophoba | - | - |
| **2364** | Aply21,Aply23,DOM43,Pf11,Pf12,Pf5,Pf6,Pf7,Pf9 | Aplysina aerophoba,Petrocia ficiformis,Aciculites cribrophora | Aply21_bin.12.fa,DOM43_bin.49.fa,-,Pf4_bin.24.fa | p__Acidobacteriota;g__;s__,- |
| **7** | Aply21 | Aplysina aerophoba | Pf12_bin.33.fa | p__Poribacteria;g__WGA-3G;s__WGA-3G sp000406005 |
| **2043** | Aply21,DOM007A,DOM007B,DOM013,DOM016,DOM057,DOM10,Pf12,Pf4,Pf9 | Oceanapia bartschi,Aplysina aerophoba,Svenzea zeai,Petrocia ficiformis,Smenospongia sp. | -,DOM016_bin.19.fa,DOM10_bin.83.fa,Pf9_bin.14.fa,Pf9_bin.10.fa | -,p__Poribacteria;g__;s__ |
| **2281** | Aply21,Pf10,Pf11,Pf12,Pf4,Pf6,Pf8 | Aplysina aerophoba,Petrocia ficiformis | Aply23_bin.19.fa,Pf9_bin.24.fa,- | p__Poribacteria;g__WGA-3G;s__,- |
| **10** | Aply21 | Aplysina aerophoba | - | - |
| **2361** | Aply22,Pf9 | Aplysina aerophoba,Petrocia ficiformis | Aply22_bin.45.fa,Pf9_bin.57.fa | p__Proteobacteria;g__Bin95;s__ |
| **2040** | Aply22,DOM057 | Aplysina aerophoba,Oceanapia bartschi | Aply22_bin.44.fa,DOM007B_bin.12.fa | p__Proteobacteria;g__;s__ |

| | | | | |
|---|---|---|---|---|
| **15** | Aply22 | Aplysina aerophoba | Pf4_bin.17.fa | p__Acidobacteriota;g__Bin61;s__ |
| **2559** | Aply22,DOM012A,DOM10,DOM40,DOM43,Pf10,Pf11,Pf12,Pf4,Pf5,Pf7,gb10,gb1,gb278,gb2_2,gb305,gb3_2,gb4_2,gb6,gb7,gb8_2 | Petrosia hartmani,Aplysina aerophoba,Aciculites cribrophora,Aplysina sp.,Petrocia ficiformis,Geodia barretti CAN,Geodia barretti NOR,Smenospongia sp.,Geodia atlantica | -,DOM10_bin.73.fa,Pf11_bin.39.fa,gb7_bin.52.fa,gb305_bin.22.fa | -,p__Acidobacteriota;g__UBA8438;s__ |
| **2402** | Aply22,DOM007A,DOM007B,DOM011,DOM016,DOM10,DOM33,Pf10,Pf11,Pf6,Pf7,gb126,gb305,gb3_2,gb5_2,gb7,gb8_2,gb9 | Oceanapia bartschi,Neopetrosia sp.,Aplysina aerophoba,Aplysina sp.,Petrocia ficiformis,Geodia barretti CAN,Geodia barretti NOR,Smenospongia sp.,Geodia atlantica | Pf7_bin.67.fa,DOM007A_bin.13.fa,DOM33_bin.49.fa,-,gb305_bin.13.fa | p__Latescibacterota;g__;s__,- |
| **19** | Aply23 | Aplysina aerophoba | Aply23_bin.14.fa | p__Proteobacteria;g__Bin36;s__Bin36 sp002239085 |
| **20** | Aply23 | Aplysina aerophoba | - | - |
| **21** | Aply23 | Aplysina aerophoba | Aply21_bin.6.fa | p__Proteobacteria;g__;s__ |
| **23** | Aply23 | Aplysina aerophoba | - | - |
| **1912** | DOM005,DOM013,DOM015,DOM026,DOM045,DOM10,DOM33,DOM40,Pf5 | Petrosia hartmani,Svenzea zeai,Aplysina sp.,Petrocia ficiformis,Verongula sp.,Smenospongia sp.,Siphonodictyon coralliphagum | DOM005_bin.1.fa,DOM013_bin.35.fa,DOM40_bin.23.fa,DOM026_bin.32.fa,DOM10_bin.14.fa,Pf5_bin.13.fa | p__Poribacteria;g__PCPOR2b;s__ |
| **2027** | DOM007A,DOM007B,DOM016,DOM057 | Oceanapia bartschi | DOM057_bin.59.fa | p__Poribacteria;g__PCPOR2b;s__ |
| **1823** | DOM007A,DOM007B,DOM016,DOM057,DOM40 | Petrosia hartmani,Oceanapia bartschi | DOM057_bin.37.fa,DOM40_bin.4.fa | p__Latescibacterota;g__GCA-2724215;s__ |
| **1853** | DOM007A,DOM007B,DOM013,DOM016,DOM057,DOM33,DOM40,gb4_2 | Oceanapia bartschi,Petrosia hartmani,Svenzea zeai,Aplysina sp.,Geodia barretti NOR | DOM007B_bin.31.fa,- | p__Poribacteria;g__;s__,- |

| | | | | |
|---|---|---|---|---|
| **2039** | DOM007A,DOM007B,DOM011,DOM016,DOM044,DOM045,DOM057,DOM10,DOM14B,gb1,gb5_2,gb6 | Oceanapia bartschi,Neopetrosia sp.,Caminus sp.,Verongula sp.,Geodia barretti NOR,Smenospongia sp. | -,DOM057_bin.11.fa,DOM011_bin.51.fa,DOM044_bin.2.fa,DOM045_bin.56.fa,DOM14B_bin.10.fa | -,p__Acidobacteriota;g__;s__ |
| **1854** | DOM007A,DOM007B,DOM013,DOM015,DOM016,DOM057 | Svenzea zeai,Oceanapia bartschi | DOM015_bin.57.fa,- | p__Acidobacteriota;g__;s__,- |
| **2506** | DOM007A,DOM007B,DOM33,Pf8,gb4_2,gb8_2 | Aplysina sp.,Oceanapia bartschi,Petrocia ficiformis,Geodia barretti NOR | DOM007B_bin.3.fa,- | p__Acidobacteriota;g__;s__,- |
| **1948** | DOM007A,DOM007B,DOM011,DOM015,DOM016,DOM026,DOM057,DOM10,DOM40 | Oceanapia bartschi,Neopetrosia sp.,Petrosia hartmani,Svenzea zeai,Verongula sp.,Smenospongia sp. | DOM057_bin.24.fa,DOM40_bin.82.fa,DOM015_bin.24.fa,DOM10_bin.71.fa | p__Latescibacterota;g__UBA8231;s__ |
| **1951** | DOM007A,DOM007B,DOM013,DOM015,DOM016,DOM057,DOM33,Pf4 | Svenzea zeai,Aplysina sp.,Oceanapia bartschi,Petrocia ficiformis | DOM015_bin.57.fa,-,DOM33_bin.26.fa,Pf10_bin.11.fa | p__Acidobacteriota;g__;s__,- |
| **2010** | DOM007A,DOM007B,DOM012A,DOM013,DOM015,DOM016,DOM026,DOM044,DOM045,DOM057,DOM40,DOM43 | Oceanapia bartschi,Neopetrosia sp.,Petrosia hartmani,Aciculites cribrophora,Svenzea zeai,Aplysina sp.,Verongula sp. | DOM016_bin.44.fa,DOM015_bin.62.fa,DOM011_bin.25.fa,DOM40_bin.61.fa,DOM43_bin.30.fa | p__Desulfobacterota_B;g__;s__ |
| **1892** | DOM007A,DOM007B,DOM011,DOM012A,DOM026,DOM045,DOM057,DOM10,DOM14B,DOM33 | Oceanapia bartschi,Neopetrosia sp.,Caminus sp.,Aplysina sp.,Verongula sp.,Smenospongia sp. | DOM007A_bin.28.fa,DOM011_bin.60.fa,DOM10_bin.80.fa,- | p__Acidobacteriota;g__;s__,- |
| **1834** | DOM007A,DOM007B | Oceanapia bartschi | DOM007A_bin.57.fa | p__Proteobacteria;g__;s__ |
| **2042** | DOM007A,DOM007B,DOM016,DOM026,DOM057,DOM10,DOM14B,DOM33 | Oceanapia bartschi,Caminus sp.,Aplysina sp.,Verongula sp.,Smenospongia sp. | DOM007B_bin.45.fa,- | p__Poribacteria;g__;s__,- |

| | | | | |
|---|---|---|---|---|
| **1983** | DOM007A,DOM007B,DOM016,DOM044,DOM057,DOM43 | Oceanapia bartschi,Aciculites cribrophora,Neopetrosia sp. | DOM057_bin.52.fa,-,DOM43_bin.102.fa | p__Latescibacterota;g__;s__,- |
| **1859** | DOM007A,DOM007B | Oceanapia bartschi | DOM007B_bin.34.fa | p__Poribacteria;g__MSPOR6;s__ |
| **1838** | DOM007A,DOM007B,DOM016,DOM057 | Oceanapia bartschi | DOM007A_bin.65.fa | p__Latescibacterota;g__;s__ |
| **1839** | DOM007A,DOM007B,DOM016,DOM057 | Oceanapia bartschi | DOM007B_bin.47.fa | p__Proteobacteria;g__;s__ |
| **2033** | DOM007A,DOM016,DOM057 | Oceanapia bartschi | -,DOM057_bin.20.fa | -,p__Proteobacteria;g__;s__ |
| **1959** | DOM007A,DOM016,DOM057 | Oceanapia bartschi | - | - |
| **1843** | DOM007A,DOM012A,DOM016,DOM057 | Aplysina sp.,Oceanapia bartschi | DOM057_bin.66.fa,DOM012A_bin.13.fa | p__Actinobacteriota;g__;s__ |
| **2044** | DOM007A,DOM007B,DOM016,DOM049,DOM057,Pf5,Pf6,gb5_2,gb9 | Geodia barretti NOR,Oceanapia bartschi,Petrocia ficiformis,Xestospongia muta | DOM057_bin.42.fa,-,DOM049_bin.6.fa,Pf11_bin.39.fa,gb7_bin.52.fa | p__Acidobacteriota;g__UBA8438;s__,- |
| **1845** | DOM007A | Oceanapia bartschi | DOM007A_bin.11.fa | p__Proteobacteria;g__;s__ |
| **1846** | DOM007A,DOM33 | Aplysina sp.,Oceanapia bartschi | -,DOM33_bin.5.fa | -,p__Acidobacteriota;g__;s__ |
| **2045** | DOM007A,DOM011,DOM016,DOM044,DOM057 | Oceanapia bartschi,Neopetrosia sp. | DOM40_bin.36.fa,- | p__Acidobacteriota;g__;s__,- |
| **1851** | DOM007B,DOM057 | Oceanapia bartschi | DOM007B_bin.47.fa | p__Proteobacteria;g__;s__ |
| **1869** | DOM007B | Oceanapia bartschi | DOM057_bin.66.fa | p__Actinobacteriota;g__;s__ |
| **1870** | DOM007B | Oceanapia bartschi | - | - |

| | | | | |
|---|---|---|---|---|
| **2009** | DOM011,DOM012A,DOM045,DOM10,DOM33,DOM40 | Neopetrosia sp.,Petrosia hartmani,Aplysina sp.,Verongula sp.,Smenospongia sp. | DOM011_bin.47.fa,DOM33_bin.36.fa,DOM33_bin.22.fa,DOM40_bin.51.fa | p__Latescibacterota;g__;s__ |
| **2410** | DOM011,gb10,gb1,gb5_2,gb9 | Geodia barretti NOR,Neopetrosia sp. | DOM011_bin.33.fa,gb6_bin.52.fa | p__Acidobacteriota;g__Bin61;s__ |
| **1876** | DOM011 | Neopetrosia sp. | - | - |
| **2487** | DOM011,gb10,gb1,gb2_2,gb3_2,gb4_2,gb9 | Geodia barretti NOR,Neopetrosia sp.,Geodia atlantica | - | - |
| **1879** | DOM011,DOM013,DOM015,DOM026,DOM10,DOM40 | Neopetrosia sp.,Petrosia hartmani,Svenzea zeai,Verongula sp.,Smenospongia sp. | DOM007A_bin.65.fa,DOM10_bin.10.fa,- | p__Latescibacterota;g__;s__,- |
| **1880** | DOM011 | Neopetrosia sp. | DOM011_bin.50.fa | p__Proteobacteria;g__;s__ |
| **1881** | DOM011 | Neopetrosia sp. | DOM011_bin.50.fa | p__Proteobacteria;g__;s__ |
| **1998** | DOM011,DOM044,DOM10,DOM33 | Aplysina sp.,Neopetrosia sp.,Smenospongia sp. | DOM33_bin.9.fa | p__Latescibacterota;g__;s__ |
| **1883** | DOM011 | Neopetrosia sp. | DOM011_bin.50.fa | p__Proteobacteria;g__;s__ |
| **1884** | DOM011,DOM012A,DOM026,DOM10,DOM33,DOM40,DOM43,Pf10,Pf4,Pf7 | Neopetrosia sp.,Petrosia hartmani,Aciculites cribrophora,Aplysina sp.,Petrocia ficiformis,Verongula sp.,Smenospongia sp. | DOM33_bin.28.fa,DOM10_bin.94.fa,DOM40_bin.33.fa,DOM43_bin.28.fa,Pf10_bin.11.fa | p__Acidobacteriota;g__;s__ |
| **1885** | DOM011 | Neopetrosia sp. | - | - |
| **1886** | DOM011,DOM14B,DOM33,gb7 | Caminus sp.,Aplysina sp.,Neopetrosia sp.,Geodia barretti NOR | DOM044_bin.25.fa,DOM14B_bin.48.fa,- | p__Poribacteria;g__;s__,- |
| **2549** | DOM011,DOM044,gb1,gb1_f,gb5_2,gb5_6_f,gb6 | Geodia barretti NOR,Neopetrosia sp.,Seawater ATL | - | - |
| **2322** | DOM012A,DOM044,DOM10,DOM43,Pf7 | Neopetrosia sp.,Aciculites cribrophora,Aplysina sp.,Petrocia ficiformis,Smenospongia sp. | DOM012A_bin.14.fa,DOM044_bin.23.fa,-,DOM43_bin.21.fa,Pf7_bin.1.fa | p__Chloroflexota;g__;s__,- |
| **2138** | DOM012A,DOM013,DOM026,DOM045,DOM14B,DOM33,DOM40,DOM43 | Petrosia hartmani,Aciculites cribrophora,Caminus sp.,Svenzea zeai,Aplysina sp.,Verongula sp. | DOM40_bin.36.fa,-,DOM43_bin.77.fa | p__Acidobacteriota;g__;s__,- |

| | | | | |
|---|---|---|---|---|
| **1895** | DOM012A | Aplysina sp. | DOM012A_bin.42.fa | p__Proteobacteria;g__Defluviicoccus;s__ |
| **1897** | DOM012A | Aplysina sp. | - | - |
| **2508** | DOM012A,DOM013,DOM015,DOM026,DOM10,DOM14B,DOM40,Pf10,Pf12,Pf8,gb10,gb126,gb2_2,gb3_2,gb4_2,gb8_2,gb9 | Petrosia hartmani,Caminus sp.,Svenzea zeai,Aplysina sp.,Petrocia ficiformis,Geodia barretti CAN,Verongula sp.,Geodia barretti NOR,Smenospongia sp.,Geodia atlantica | -,DOM013_bin.51.fa,DOM015_bin.46.fa,DOM14B_bin.24.fa | -,p__Acidobacteriota;g__;s__ |
| **1900** | DOM012A,DOM33 | Aplysina sp. | DOM012A_bin.42.fa,- | p__Proteobacteria;g__Defluviicoccus;s__,- |
| **1902** | DOM012A | Aplysina sp. | - | - |
| **1903** | DOM012A,DOM016,DOM40 | Petrosia hartmani,Aplysina sp.,Oceanapia bartschi | -,DOM016_bin.6.fa | -,p__Poribacteria;g__WGA-3G;s__ |
| **2098** | DOM012A,DOM14B,DOM33 | Caminus sp.,Aplysina sp. | -,DOM14B_bin.45.fa,DOM026_bin.47.fa | -,p__Tectomicrobia;g__SXND01;s__ |
| **1909** | DOM013 | Svenzea zeai | DOM015_bin.42.fa | p__Proteobacteria;g__;s__ |
| **2553** | DOM013,DOM044,DOM049,DOM43,gb10,gb126,gb1,gb278,gb2_2,gb3_2,gb4_2,gb5_2,gb6,gb7,gb8_2,gb9 | Neopetrosia sp.,Aciculites cribrophora,Svenzea zeai,Geodia barretti CAN,Geodia barretti NOR,Xestospongia muta,Geodia atlantica | DOM007A_bin.28.fa,-,DOM049_bin.18.fa,gb8_2_bin.30.fa | p__Acidobacteriota;g__;s__,- |
| **1911** | DOM013,DOM015,DOM026,DOM044,DOM057,DOM14B,DOM40,DOM43 | Oceanapia bartschi,Neopetrosia sp.,Petrosia hartmani,Aciculites cribrophora,Caminus sp.,Svenzea zeai,Verongula sp. | DOM015_bin.30.fa,DOM33_bin.49.fa,DOM044_bin.6.fa,DOM007A_bin.13.fa,DOM14B_bin.41.fa,DOM40_bin.38.fa,DOM43_bin.81.fa | p__Latescibacterota;g__;s__ |
| **2036** | DOM013,DOM015,DOM026,DOM045,DOM057,DOM10,DOM14B,DOM33,DOM43 | Oceanapia bartschi,Aciculites cribrophora,Caminus sp.,Svenzea zeai,Aplysina sp.,Verongula sp.,Smenospongia sp. | -,DOM10_bin.9.fa,DOM057_bin.8.fa,DOM057_bin.57.fa,DOM14B_bin.25.fa | -,p__Acidobacteriota;g__;s__ |

| | | | | |
|---|---|---|---|---|
| **1918** | DOM013,DOM015,DOM049 | Svenzea zeai,Xestospongia muta | DOM013_bin.24.fa,DOM049_bin.5.fa | p__Latescibacterota;g__GCA-2724215;s__ |
| **1930** | DOM013,DOM015 | Svenzea zeai | - | - |
| **1923** | DOM015 | Svenzea zeai | DOM015_bin.59.fa | p__Chloroflexota;g__;s__ |
| **2325** | DOM015,DOM14B,DOM43,Pf7,gb10,gb2_2,gb6,gb7 | Aciculites cribrophora,Caminus sp.,Svenzea zeai,Petrocia ficiformis,Geodia barretti NOR | DOM015_bin.49.fa,DOM14B_bin.29.fa,DOM43_bin.95.fa,Pf7_bin.59.fa,gb1_bin.4.fa,- | p__Latescibacterota;g__GCA-2724215;s__,- |
| **1937** | DOM015 | Svenzea zeai | DOM015_bin.18.fa | p__Proteobacteria;g__;s__ |
| **2158** | DOM015,DOM33,DOM40 | Petrosia hartmani,Svenzea zeai,Aplysina sp. | DOM015_bin.19.fa | p__Deinococcota;g__MPNL01;s__MPNL01 sp002239005 |
| **2453** | DOM015,gb2_2,gb5_2 | Svenzea zeai,Geodia barretti NOR | DOM007B_bin.31.fa,-,gb5_2_bin.1.fa | p__Poribacteria;g__;s__,- |
| **1941** | DOM015 | Svenzea zeai | - | - |
| **2255** | DOM026,DOM10,DOM14B,Pf10,Pf11,Pf12,Pf6,Pf7,Pf8 | Verongula sp.,Smenospongia sp.,Petrocia ficiformis,Caminus sp. | DOM026_bin.32.fa,DOM10_bin.47.fa,DOM10_bin.14.fa,-,Pf8_bin.6.fa | p__Poribacteria;g__PCPOR2b;s__,- |
| **2123** | DOM026,DOM33 | Verongula sp.,Aplysina sp. | DOM33_bin.28.fa | p__Acidobacteriota;g__;s__ |
| **1975** | DOM026 | Verongula sp. | - | - |
| **2286** | DOM026,DOM044,DOM045,DOM33,DOM43,Pf12,Pf4,Pf5,Pf9 | Neopetrosia sp.,Aciculites cribrophora,Aplysina sp.,Petrocia ficiformis,Verongula sp. | -,DOM044_bin.47.fa,DOM33_bin.71.fa,DOM43_bin.27.fa,Pf4_bin.17.fa,Pf5_bin.39.fa | -,p__Acidobacteriota;g__Bin61;s__ |
| **1977** | DOM026 | Verongula sp. | - | - |
| **1979** | DOM026 | Verongula sp. | - | - |
| **1981** | DOM026 | Verongula sp. | DOM10_bin.73.fa | p__Acidobacteriota;g__UBA8438;s__ |
| **1985** | DOM044 | Neopetrosia sp. | DOM044_bin.33.fa | p__Acidobacteriota;g__;s__ |

| | | | | |
|---|---|---|---|---|
| **2077** | DOM044,DOM10,DOM33 | Aplysina sp.,Neopetrosia sp.,Smenospongia sp. | DOM044_bin.27.fa | p__Chloroflexota;g__Bin90;s__ |
| **1987** | DOM044 | Neopetrosia sp. | DOM044_bin.13.fa | p__Proteobacteria;g__;s__ |
| **1989** | DOM044 | Neopetrosia sp. | DOM044_bin.13.fa | p__Proteobacteria;g__;s__ |
| **1992** | DOM044 | Neopetrosia sp. | DOM044_bin.42.fa | p__Proteobacteria;g__Endozoicomonas;s__ |
| **1994** | DOM044 | Neopetrosia sp. | DOM044_bin.41.fa | p__Tectomicrobia;g__;s__ |
| **1997** | DOM044 | Neopetrosia sp. | - | - |
| **2554** | DOM044,DOM40,gb10,gb2_2,gb4_2,gb6,gb8_2 | Petrosia hartmani,Geodia barretti NOR,Neopetrosia sp. | DOM044_bin.25.fa,DOM40_bin.32.fa,-,gb4_2_bin.25.fa | p__Poribacteria;g__;s__,- |
| **2011** | DOM045 | Verongula sp. | DOM10_bin.30.fa | p__Latescibacterota;g__GCA-2724215;s__ |
| **2012** | DOM045 | Verongula sp. | - | - |
| **2013** | DOM045,gb10,gb2_2,gb3_2,gb6,gb9 | Verongula sp.,Geodia barretti NOR,Geodia atlantica | -,gb5_2_bin.1.fa | -,p__Poribacteria;g__;s__ |
| **2014** | DOM045 | Verongula sp. | DOM045_bin.14.fa | p__Latescibacterota;g__;s__ |
| **2591** | DOM045,DOM40,DOM43,gb10,gb3_f,gb4_2,gb8_2 | Petrosia hartmani,Seawater ATL,Aciculites cribrophora,Verongula sp.,Geodia barretti NOR | -,DOM40_bin.52.fa | -,p__Chloroflexota;g__;s__ |
| **2386** | DOM045,DOM14B,gb10 | Verongula sp.,Geodia barretti NOR,Caminus sp. | - | - |
| **2017** | DOM045 | Verongula sp. | - | - |
| **2359** | DOM049,Pf10,Pf11,Pf5,Pf7,Pf9 | Petrocia ficiformis,Xestospongia muta | DOM049_bin.10.fa,Pf10_bin.9.fa,Pf9_bin.40.fa | p__Proteobacteria;g__;s__ |
| **2019** | DOM049 | Xestospongia muta | DOM049_bin.10.fa | p__Proteobacteria;g__;s__ |
| **2020** | DOM049 | Xestospongia muta | DOM049_bin.14.fa | p__Proteobacteria;g__Bin55;s__ |

| | | | | |
|---|---|---|---|---|
| **2023** | DOM049 | Xestospongia muta | DOM049_bin.11.fa | p__Acidobacteriota;g__Bin61;s__ |
| **2025** | DOM049 | Xestospongia muta | - | - |
| **2050** | DOM057 | Oceanapia bartschi | DOM057_bin.66.fa | p__Actinobacteriota;g__;s__ |
| **2526** | DOM057,DOM10,DOM14B,gb10,gb1,gb278,gb2_2,gb3_2,gb4_2,gb5_2,gb6,gb7,gb8_2,gb9 | Oceanapia bartschi,Caminus sp.,Geodia barretti CAN,Geodia barretti NOR,Smenospongia sp.,Geodia atlantica | DOM057_bin.62.fa,DOM10_bin.95.fa,DOM14B_bin.70.fa,gb8_2_bin.1.fa | p__Proteobacteria;g__Defluviicoccus;s__ |
| **2054** | DOM10 | Smenospongia sp. | DOM10_bin.63.fa | p__Acidobacteriota;g__Bin61;s__ |
| **2056** | DOM10 | Smenospongia sp. | DOM10_bin.95.fa | p__Proteobacteria;g__Defluviicoccus;s__ |
| **2513** | DOM10,gb4_2 | Geodia barretti NOR,Smenospongia sp. | DOM10_bin.38.fa,gb5_2_bin.24.fa | p__Chloroflexota;g__;s__ |
| **2060** | DOM10 | Smenospongia sp. | DOM10_bin.17.fa | p__Poribacteria;g__MSPOR6;s__ |
| **2061** | DOM10 | Smenospongia sp. | DOM10_bin.94.fa | p__Acidobacteriota;g__;s__ |
| **2063** | DOM10 | Smenospongia sp. | DOM10_bin.33.fa | p__Actinobacteriota;g__;s__ |
| **2067** | DOM10 | Smenospongia sp. | DOM011_bin.57.fa | p__Proteobacteria;g__;s__ |
| **2070** | DOM10 | Smenospongia sp. | DOM011_bin.57.fa | p__Proteobacteria;g__;s__ |
| **2076** | DOM10,Pf4,Pf8,gb278 | Smenospongia sp.,Petrocia ficiformis,Geodia barretti CAN | DOM012A_bin.20.fa,-,DOM044_bin.16.fa,gb278_bin.45.fa | p__Proteobacteria;g__;s__,- |
| **2080** | DOM10 | Smenospongia sp. | - | - |
| **2081** | DOM10 | Smenospongia sp. | DOM10_bin.59.fa | p__Chloroflexota;g__;s__ |
| **2084** | DOM14B | Caminus sp. | - | - |
| **2086** | DOM14B | Caminus sp. | DOM14B_bin.5.fa | p__Acidobacteriota;g__;s__ |

| | | | | |
|---|---|---|---|---|
| **2557** | DOM14B,gb1,gb3_2,gb4_2,gb6,gb8_2,gb9 | Caminus sp.,Geodia barretti NOR,Geodia atlantica | DOM14B_bin.48.fa,gb4_2_bin.25.fa,- | p__Poribacteria;g__;s__,- |
| **2089** | DOM14B | Caminus sp. | DOM14B_bin.17.fa | p__Proteobacteria;g__UBA2168;s__ |
| **2090** | DOM14B | Caminus sp. | DOM14B_bin.19.fa | p__Acidobacteriota;g__;s__ |
| **2091** | DOM14B | Caminus sp. | DOM14B_bin.70.fa | p__Proteobacteria;g__Defluviicoccus;s__ |
| **2092** | DOM14B | Caminus sp. | - | - |
| **2093** | DOM14B | Caminus sp. | - | - |
| **2333** | DOM14B,Pf10,Pf11,Pf12,Pf4,Pf5,Pf6,Pf7,Pf8,Pf9 | Caminus sp.,Petrocia ficiformis | DOM14B_bin.22.fa,Pf5_bin.31.fa | p__Poribacteria;g__MSPOR6;s__ |
| **2097** | DOM14B | Caminus sp. | DOM14B_bin.76.fa | p__Verrucomicrobiota;g__;s__ |
| **2271** | DOM14B,Pf10,Pf4,Pf5,Pf6,Pf8 | Caminus sp.,Petrocia ficiformis | -,Pf5_bin.19.fa | -,p__Acidobacteriota;g__;s__ |
| **2105** | DOM14B | Caminus sp. | - | - |
| **2108** | DOM14B | Caminus sp. | DOM14B_bin.49.fa | p__Proteobacteria;g__;s__ |
| **2109** | DOM14B | Caminus sp. | - | - |
| **2110** | DOM14B | Caminus sp. | - | - |
| **2111** | DOM14B | Caminus sp. | - | - |
| **2112** | DOM14B | Caminus sp. | - | - |
| **2119** | DOM33 | Aplysina sp. | DOM33_bin.17.fa | p__Actinobacteriota;g__Bin76;s__ |
| **2127** | DOM33 | Aplysina sp. | DOM33_bin.72.fa | p__Poribacteria;g__MSPOR6;s__ |
| **2137** | DOM33 | Aplysina sp. | - | - |
| **2139** | DOM40 | Petrosia hartmani | DOM40_bin.21.fa | p__Latescibacterota;g__;s__ |
| **2141** | DOM40 | Petrosia hartmani | DOM40_bin.45.fa | p__Chloroflexota;g__;s__ |

| | | | | |
|---|---|---|---|---|
| **2143** | DOM40 | Petrosia hartmani | DOM40_bin.64.fa | p__Proteobacteria;g__;s__ |
| **2144** | DOM40 | Petrosia hartmani | DOM40_bin.50.fa | p__Proteobacteria;g__;s__ |
| **2147** | DOM40 | Petrosia hartmani | DOM40_bin.50.fa | p__Proteobacteria;g__;s__ |
| **2153** | DOM40 | Petrosia hartmani | DOM40_bin.77.fa | p__Proteobacteria;g__;s__ |
| **2154** | DOM40 | Petrosia hartmani | DOM40_bin.70.fa | p__Proteobacteria;g__;s__ |
| **2161** | DOM40 | Petrosia hartmani | DOM40_bin.64.fa | p__Proteobacteria;g__;s__ |
| **2163** | DOM40 | Petrosia hartmani | DOM40_bin.84.fa | p__Latescibacterota;g__;s__ |
| **2165** | DOM43 | Aciculites cribrophora | DOM43_bin.67.fa | p__Proteobacteria;g__;s__ |
| **2169** | DOM43 | Aciculites cribrophora | DOM43_bin.19.fa | p__Acidobacteriota;g__;s__ |
| **2171** | DOM43 | Aciculites cribrophora | DOM43_bin.62.fa | p__Latescibacterota;g__;s__ |
| **2172** | DOM43 | Aciculites cribrophora | DOM43_bin.3.fa | p__Chloroflexota;g__;s__ |
| **2173** | DOM43,Pf11,Pf5 | Petrocia ficiformis,Aciculites cribrophora | DOM43_bin.27.fa,Pf5_bin.39.fa | p__Acidobacteriota;g__Bin61;s__ |
| **2174** | DOM43 | Aciculites cribrophora | - | - |
| **2175** | DOM43 | Aciculites cribrophora | - | - |
| **2176** | DOM43 | Aciculites cribrophora | DOM43_bin.93.fa | p__Proteobacteria;g__;s__ |
| **2177** | DOM43 | Aciculites cribrophora | DOM43_bin.47.fa | p__Latescibacterota;g__UBA8231;s__ |
| **2178** | DOM43 | Aciculites cribrophora | DOM43_bin.83.fa | p__Latescibacterota;g__;s__ |

| | | | | |
|---|---|---|---|---|
| **2180** | DOM43 | Aciculites cribrophora | DOM43_bin.32.fa | p__Tectomicrobia;g__;s__ |
| **2183** | DOM43 | Aciculites cribrophora | DOM43_bin.12.fa | p__Verrucomicrobiota;g__;s__ |
| **2184** | DOM43 | Aciculites cribrophora | - | - |
| **2185** | DOM43 | Aciculites cribrophora | DOM43_bin.54.fa | p__Actinobacteriota;g__;s__ |
| **2186** | DOM43 | Aciculites cribrophora | DOM43_bin.39.fa | p__Latescibacterota;g__;s__ |
| **2188** | DOM43 | Aciculites cribrophora | - | - |
| **2189** | DOM43 | Aciculites cribrophora | DOM43_bin.22.fa | p__;g__;s__ |
| **2190** | DOM43 | Aciculites cribrophora | - | - |
| **2191** | DOM43 | Aciculites cribrophora | DOM43_bin.62.fa | p__Latescibacterota;g__;s__ |
| **2192** | DOM43 | Aciculites cribrophora | - | - |
| **2193** | DOM43 | Aciculites cribrophora | - | - |
| **2194** | DOM43 | Aciculites cribrophora | - | - |
| **2195** | DOM43 | Aciculites cribrophora | - | - |
| **2198** | DOM43 | Aciculites cribrophora | - | - |
| **2202** | DOM43 | Aciculites cribrophora | DOM43_bin.94.fa | p__Proteobacteria;g__;s__ |
| **2203** | DOM43 | Aciculites cribrophora | DOM43_bin.101.fa | p__Acidobacteriota;g__;s__ |
| **2204** | DOM43 | Aciculites cribrophora | - | - |
| **2205** | DOM43 | Aciculites cribrophora | DOM43_bin.70.fa | p__Verrucomicrobiota;g__Moanabacter;s__ |
| **2206** | DOM43 | Aciculites cribrophora | - | - |
| **2232** | Pf10,Pf11,Pf12,Pf6,Pf7,Pf8,Pf9 | Petrocia ficiformis | Pf9_bin.7.fa | p__Proteobacteria;g__Bin55;s__ |
| **2239** | Pf10,Pf11,Pf5,Pf6,Pf7,Pf9 | Petrocia ficiformis | Pf11_bin.50.fa | p__Proteobacteria;g__Bin65;s__ |

| | | | | |
|---|---|---|---|---|
| **2258** | Pf10,Pf12 | Petrocia ficiformis | -,Pf4_bin.35.fa | -,p__Proteobacteria;g__Porisulfidus;s__ |
| **2270** | Pf10,Pf11,Pf4,Pf5,Pf6,Pf7,Pf8 | Petrocia ficiformis | Pf5_bin.30.fa | p__Verrucomicrobiota;g__UBA2970;s__ |
| **2299** | Pf10,Pf11,Pf12,Pf6,Pf7,Pf8,Pf9 | Petrocia ficiformis | Pf9_bin.7.fa | p__Proteobacteria;g__Bin55;s__ |
| **2218** | Pf10 | Petrocia ficiformis | - | - |
| **2332** | Pf10,Pf7,gb278 | Petrocia ficiformis,Geodia barretti CAN | Pf10_bin.11.fa,gb5_2_bin.68.fa | p__Acidobacteriota;g__;s__ |
| **2225** | Pf10 | Petrocia ficiformis | - | - |
| **2226** | Pf10,Pf6 | Petrocia ficiformis | - | - |
| **2227** | Pf10 | Petrocia ficiformis | - | - |
| **2273** | Pf10,Pf11,Pf12,Pf4 | Petrocia ficiformis | -,Pf4_bin.36.fa | -,p__Cyanobacteria;g__Aphanocapsa;s__Aphanocapsa feldmannii |
| **2230** | Pf10 | Petrocia ficiformis | - | - |
| **2238** | Pf11,Pf7,Pf8 | Petrocia ficiformis | Pf7_bin.23.fa | p__Proteobacteria;g__;s__ |
| **2366** | Pf11,Pf4,Pf5,Pf6,Pf7,Pf8,Pf9 | Petrocia ficiformis | -,Pf7_bin.64.fa | -,p__Actinobacteriota;g__;s__ |
| **2362** | Pf11,Pf7,Pf8,Pf9 | Petrocia ficiformis | Pf7_bin.5.fa | p__Latescibacterota;g__;s__ |
| **2248** | Pf11 | Petrocia ficiformis | - | - |
| **2259** | Pf12 | Petrocia ficiformis | Pf11_bin.25.fa | p__Chloroflexota;g__Bin22;s__ |
| **2261** | Pf12 | Petrocia ficiformis | Pf10_bin.11.fa | p__Acidobacteriota;g__;s__ |
| **2265** | Pf4 | Petrocia ficiformis | - | - |
| **2266** | Pf4 | Petrocia ficiformis | Pf4_bin.61.fa | p__Proteobacteria;g__Bin36;s__ |

| | | | | |
|---|---|---|---|---|
| **2268** | Pf4 | Petrocia ficiformis | Pf4_bin.61.fa | p__Proteobacteria;g__Bin36;s__ |
| **2278** | Pf4 | Petrocia ficiformis | Pf7_bin.54.fa | p__Chloroflexota;g__;s__ |
| **2282** | Pf4,Pf6,Pf8 | Petrocia ficiformis | -,Pf4_bin.33.fa | -,p__Acidobacteriota;g__UBA8438;s__ |
| **2283** | Pf4 | Petrocia ficiformis | Pf4_bin.4.fa | p__Latescibacterota;g__GCA-2724215;s__ |
| **2345** | Pf5,Pf6,Pf7,Pf8 | Petrocia ficiformis | -,Pf7_bin.19.fa | -,p__Acidobacteriota;g__;s__ |
| **2296** | Pf5,gb3_2,gb8_2,gb9 | Geodia barretti NOR,Petrocia ficiformis,Geodia atlantica | -,gb1_bin.4.fa | -,p__Latescibacterota;g__GCA-2724215;s__ |
| **2297** | Pf5 | Petrocia ficiformis | Pf7_bin.23.fa | p__Proteobacteria;g__;s__ |
| **2328** | Pf7,Pf8 | Petrocia ficiformis | - | - |
| **2330** | Pf7 | Petrocia ficiformis | Pf9_bin.53.fa | p__Gemmatimonadota;g__Bin94;s__ |
| **2334** | Pf7 | Petrocia ficiformis | Pf11_bin.25.fa | p__Chloroflexota;g__Bin22;s__ |
| **2356** | Pf8 | Petrocia ficiformis | - | - |
| **2367** | Pf9 | Petrocia ficiformis | Pf9_bin.54.fa | p__Latescibacterota;g__UBA8231;s__ |
| **2522** | gb10,gb1,gb2_2,gb3_2,gb4_2,gb5_2,gb6,gb7,gb8_2,gb9 | Geodia barretti NOR,Geodia atlantica | gb5_2_bin.9.fa,- | p__Acidobacteriota;g__UBA8438;s__,- |
| **2468** | gb10,gb126,gb1,gb278,gb2_2,gb305,gb7,gb8_2 | Geodia barretti NOR,Geodia barretti CAN | -,gb126_bin.26.fa | -,p__Acidobacteriota;g__;s__ |
| **2609** | gb10,gb2_2,gb6,gb8_2,gb9 | Geodia barretti NOR | - | - |

| | | | | |
|---|---|---|---|---|
| **2380** | gb10,gb7 | Geodia barretti NOR | - | - |
| **2527** | gb10,gb2_2,gb5_2,gb7 | Geodia barretti NOR | -,gb4_2_bin.25.fa | -,p__Poribacteria;g__;s__ |
| **2510** | gb10,gb126,gb1,gb278,gb305,gb4_2 | Geodia barretti NOR,Geodia barretti CAN | -,gb126_bin.18.fa | -,p__Acidobacteriota;g__;s__ |
| **2484** | gb10,gb3_2,gb4_2,gb8_2 | Geodia barretti NOR,Geodia atlantica | - | - |
| **2389** | gb10 | Geodia barretti NOR | - | - |
| **2620** | gb10_f,gb1_f,gb3_f,gb5_6_f,gb9_f | Seawater ATL | gb3_f_bin.30.fa | p__Proteobacteria;g__REDSEA-S09-B13;s__REDSEA-S09-B13 sp002456995 |
| **2391** | gb10_f,gb1_f,gb2_f,gb5_6_f | Seawater ATL | gb3_f_bin.12.fa | p__Proteobacteria;g__UBA9145;s__ |
| **2621** | gb10_f,gb1_f,gb2_f,gb3_f,gb5_6_f,gb9_f | Seawater ATL | gb10_f_bin.35.fa,- | p__Proteobacteria;g__;s__,- |
| **2495** | gb10_f,gb1_f,gb2_f,gb3_f,gb5_6_f,gb9_f | Seawater ATL | gb9_f_bin.12.fa,- | p__Proteobacteria;g__Thioglobus;s__Thioglobus singularis,- |
| **2426** | gb10_f,gb1_f,gb5_6_f,gb9_f | Seawater ATL | gb10_f_bin.30.fa | p__Proteobacteria;g__HTCC2207;s__HTCC2207 sp002685195 |
| **2463** | gb10_f,gb2_f,gb5_6_f | Seawater ATL | -,gb5_6_f_bin.55.fa | -,p__Verrucomicrobiota;g__EC70;s__ |
| **2396** | gb10_f | Seawater ATL | - | - |
| **2399** | gb10_f,gb2_f,gb3_f,gb5_6_f | Seawater ATL | - | - |
| **2398** | gb10_f | Seawater ATL | gb10_f_bin.64.fa | p__Latescibacterota;g__;s__ |
| **2400** | gb10_f | Seawater ATL | gb10_f_bin.41.fa | p__Planctomycetota;g__GCA-2686945;s__ |

| | | | | |
|---|---|---|---|---|
| **2444** | gb126,gb2_2,gb305 | Geodia barretti NOR,Geodia barretti CAN | gb5_2_bin.68.fa | p__Acidobacteriota;g__;s__ |
| **2457** | gb126,gb2_2,gb6,gb7,gb8_2,gb9 | Geodia barretti NOR,Geodia barretti CAN | - | - |
| **2616** | gb1,gb5_2,gb9 | Geodia barretti NOR | - | - |
| **2490** | gb1,gb2_2,gb3_2,gb4_2,gb5_2,gb6,gb7 | Geodia barretti NOR,Geodia atlantica | - | - |
| **2623** | gb1_f,gb2_f,gb3_f,gb5_6_f,gb9_f | Seawater ATL | gb9_f_bin.17.fa | p__Proteobacteria;g__UBA11791;s__ |
| **2427** | gb1_f | Seawater ATL | gb1_f_bin.23.fa | p__Actinobacteriota;g__;s__ |
| **2428** | gb1_f,gb2_f,gb3_f | Seawater ATL | gb1_f_bin.29.fa | p__Proteobacteria;g__GCA-2726415;s__ |
| **2431** | gb278,gb305,gb5_2,gb6 | Geodia barretti NOR,Geodia barretti CAN | gb278_bin.34.fa | p__Proteobacteria;g__;s__ |
| **2435** | gb278 | Geodia barretti CAN | gb305_bin.51.fa | p__Acidobacteriota;g__;s__ |
| **2437** | gb278 | Geodia barretti CAN | - | - |
| **2518** | gb2_2,gb305,gb4_2,gb6,gb8_2,gb9 | Geodia barretti NOR,Geodia barretti CAN | -,gb278_bin.45.fa | -,p__Proteobacteria;g__;s__ |
| **2519** | gb2_2,gb4_2,gb6,gb8_2 | Geodia barretti NOR | gb6_bin.52.fa | p__Acidobacteriota;g__Bin61;s__ |
| **2466** | gb2_f | Seawater ATL | - | - |
| **2467** | gb2_f | Seawater ATL | - | - |
| **2471** | gb305 | Geodia barretti CAN | gb305_bin.15.fa | p__Proteobacteria;g__;s__ |
| **2479** | gb3_2 | Geodia atlantica | gb4_2_bin.48.fa | p__Poribacteria;g__MSPOR6;s__ |
| **2486** | gb3_2 | Geodia atlantica | - | - |
| **2489** | gb3_2 | Geodia atlantica | - | - |
| **2491** | gb3_2 | Geodia atlantica | - | - |
| **2493** | gb3_f | Seawater ATL | - | - |

| | | | | |
|---|---|---|---|---|
| **2500** | gb3_f | Seawater ATL | - | - |
| **2514** | gb4_2 | Geodia barretti NOR | - | - |
| **2516** | gb4_2 | Geodia barretti NOR | - | - |
| **2520** | gb4_2 | Geodia barretti NOR | - | - |
| **2525** | gb5_2 | Geodia barretti NOR | gb2_2_bin.49.fa | p__Proteobacteria;g__;s__ |
| **2538** | gb5_6_f | Seawater ATL | - | - |
| **2547** | gb5_6_f | Seawater ATL | - | - |
| **2548** | gb5_6_f | Seawater ATL | - | - |
| **2550** | gb5_6_f | Seawater ATL | - | - |
| **2551** | gb5_6_f | Seawater ATL | - | - |
| **2555** | gb6 | Geodia barretti NOR | - | - |
| **2577** | gb7 | Geodia barretti NOR | - | - |
| **2619** | gb9_f | Seawater ATL | gb9_f_bin.8.fa | p__Proteobacteria;g__Methylobacterium;s__Methylobacterium sp900112625 |
| **2625** | gb9_f | Seawater ATL | gb5_6_f_bin.38.fa | p__Chloroflexota;g__UBA11795;s__ |
| **2627** | gb9_f | Seawater ATL | - | - |
| **2630** | sw_7,sw_8,sw_9 | Seawater MED | sw_7_bin.7.fa | p__Proteobacteria;g__Henriciella;s__Henriciella sp002172915 |
| **2629** | sw_7,sw_8,sw_9 | Seawater MED | sw_9_bin.6.fa | p__Proteobacteria;g__;s__ |
| **2631** | sw_8 | Seawater MED | - | - |
| **2633** | sw_8 | Seawater MED | sw_9_bin.8.fa | p__Proteobacteria;g__UBA8309;s__UBA8309 sp002457765 |

**Table S3.2**. Description of connected components featuring included gene cluster families (GCFs)and representative GCF.

| CC | GCFs | Reps |
|---|---|---|
| A | 2249, 2410, 2286, 2173, 2519 | 2410, 2286 |
| E | 2402, 1911 | 2402 |
| K | 1896, 2043, 2281, 1903 | 2043 |
| H | 2559, 2044, 2282 | 2559 |
| R | 1912, 2027, 2255 | 1912 |
| C | 1892, 2553 | 1892 |
| B | 2002, 2039, 2506, 2549, 2508, 2036, 2345, 2468, 2510, 2457 | 2002, 2039 |
| D | 1983, 2009, 2362 | 1983 |
| F | 1823, 1918, 2325, 2296 | 2325 |
| J | 1854, 1884 | 1884 |
| I | 1886, 2554 | 2554 |
| P | 2045, 2138 | 2138 |
| G | 2270 | 2270 |
| O | 2526 | 2526 |
| M | 2557, 2527 | 2557 |
| T | 2484, 2616 | 2484 |
| L | 2010 | 2010 |
| N | 2522 | 2522 |
| Q | 2609 | 2609 |
| S | 2366 | 2366 |

**Table S4.** Detailed BGC gene annotations

Table S4.1. Description of ORF borders for BGCs displayed in Figure 1**.**

| Contig | GCF | CC | Start ORF | Stop ORF |
|---|---|---|---|---|
| Pf12_c00118 | 2286 | A | 1 | 34 |
| gb1_c01994 | 2410 | A | 1 | 12 |
| gb5_2_c00041 | 2402 | E | 10 | 25 |
| Pf12_c00284 | 2043 | K | 17 | 24 |
| Pf11_c00818 | 2559 | H | 12 | 22 |
| DOM045_c00026 | 1912 | R | 10 | 24 |
| DOM007B_c00243 | 1892 | C | 1 | 21 |
| DOM045_c00004 | 2002 | B | 15 | 25 |
| DOM016_c00717 | 2039 | B | 19 | 31 |
| DOM044_c00402 | 1983 | D | 5 | 24 |
| Pf7_c00820 | 2325 | F | 12 | 27 |
| DOM43_c00257 | 1884 | J | 13 | 27 |
| gb4_2_c00117 | 2554 | I | 3 | 20 |
| DOM026_c00344 | 2138 | P | 7 | 22 |
| Pf8_c00244 | 2270 | G | 12 | 24 |
| DOM14B_c00569 | 2526 | O | 1 | 26 |
| gb4_2_c02126 | 2557 | M | 1 | 10 |
| gb3_2_c07789 | 2484 | T | 1 | 7 |
| DOM057_c01231 | 2010 | L | 11 | 26 |
| gb4_2_c00127 | 2522 | N | 2 | 17 |
| gb9_c09452 | 2609 | Q | 1 | 6 |
| Pf7_c00139 | 2366 | S | 13 | 19 |

Table S4.2.antiSMASH annotations of contig Pf12_c00118, detected in GCF 2286 and CC A

| Gene | SMCOG | PFAM & GO | Notes |
|---|---|---|---|
| **1** | | PF01979.22 (Amidohydrolase family)<br>PF01979.22: GO:0016787: hydrolase activity | |
| **2** | SMCOG1065:ABC-2 type transporter | PF01061.26 (ABC-2 type transporter)<br>PF01061.26: GO:0016020: membrane | transporter |
| **3** | SMCOG1000:ABC transporter ATP-binding protein | PF00005.29 (ABC transporter)<br>PF00005.29: GO:0005524: ATP binding | transporter |
| **4** | SMCOG1293:aldehyde oxidase and xanthine dehydrogenase | PF01315.24 (Aldehyde oxidase and xanthine dehydrogenase, a/b hammerhead domain)<br>PF02738.20 (Molybdopterin-binding domain of aldehyde dehydrogenase)<br>PF02738.20: GO:0016491: oxidoreductase activity<br>PF02738.20: GO:0055114: oxidation-reduction process | biosynthetic-additional |
| **5** | | PF01850.23 (PIN domain) | |
| **6** | | PF01402.23 (Ribbon-helix-helix protein, copG family)<br>PF01402.23: GO:0006355: regulation of transcription, DNA-templated | |
| **7** | | PF01145.27 (SPFH domain / Band 7 family)<br>PF16200.7 (C-terminal region of band_7) | |
| **8** | | PF01957.20 (NfeD-like C-terminal, partner-binding) | |
| **9** | | PF05063.16 (MethylTransferase-A70) | |
| **10** | SMCOG1168:O-succinylhomoserine sulfhydrylase | PF01053.22 (Cys/Met metabolism PLP-dependent enzyme)<br>PF01053.22: GO:0030170: pyridoxal phosphate binding<br>PF01053.22: GO:0019346: transsulfuration | biosynthetic-additional |
| **11** | | PF01242.21 (6-pyruvoyl tetrahydropterin synthase) | |

| | | | |
|---|---|---|---|
| **12** | | - | |
| **13** | SMCOG1034:cytochrome P450 | PF00067.24 (Cytochrome P450)<br>PF00067.24: GO:0005506: iron ion binding<br>PF00067.24: GO:0016705: oxidoreductase activity, acting on paired donors, with incorporation or reduction of molecular oxygen<br>PF00067.24: GO:0020037: heme binding<br>PF00067.24: GO:0055114: oxidation-reduction process | biosynthetic-additional<br>MiBIG:BGC0000678.1, sav_2999, Streptomyces avermitilis, Terpene, pentalenolactone |
| **14** | | PF04055.23 (Radical SAM superfamily)<br>PF04055.23: GO:0003824: catalytic activity<br>PF04055.23: GO:0051536: iron-sulfur cluster binding | biosynthetic-additional |
| **15** | SMCOG1119:halogenase | PF04820.16 (Tryptophan halogenase) | Biosynthetic<br>MiBIG: BGC0001465.1, bmp2, Marinomonas mediterranea MMB-1, Other, bromopyrroles/bromophenols |
| **16** | SMCOG1002:AMP-dependent synthetase and ligase | PF00501.30 (AMP-binding enzyme)<br>PF13193.8 (AMP-binding enzyme C-terminal domain) | Biosynthetic<br>MiBIG: BGC0000649.1, fadD, Streptomyces griseus, Terpene, carotenoid |
| **17** | | PF06983.15 (3-demethylubiquinone-9 3-methyltransferase) | |
| **18** | | PF09365.12 (Conserved hypothetical protein (DUF2461))<br>TIGR02453 (TIGR02453: TIGR02453 family protein) | |
| **19** | | - | |
| **20** | | PF02544.18 (3-oxo-5-alpha-steroid 4-dehydrogenase)<br>PF02544.18: GO:0016627: oxidoreductase activity, acting on the CH-CH group of donors<br>PF02544.18: GO:0006629: lipid metabolic process | |

| | | | |
|---|---|---|---|
| **21** | SMCOG1010:NAD-dependent epimerase/dehydratase | PF01370.23 (NAD dependent epimerase/dehydratase family)<br>PF01370.23: GO:0003824: catalytic activity | biosynthetic-additional |
| **22** | SMCOG1240:riboflavin biosynthesis protein RibD | PF00383.25 (Cytidine and deoxycytidylate deaminase zinc-binding region) | biosynthetic-additional |
| **23** | | PF00291.27 (Pyridoxal-phosphate dependent enzyme)<br>(PLP-dependent enzymes superfamily represented by the beta subunit of tryptophan synthase)<br>TIGR01274 (ACC_deam: 1-aminocyclopropane-1-carboxylate deaminase) | |
| **24** | SMCOG1036:alpha/beta hydrolase fold protein | PF00561.22 (alpha/beta hydrolase fold) | biosynthetic-additional |
| **25** | SMCOG1001:short-chain dehydrogenase/reductase SDR | PF00106.27 (short chain dehydrogenase)<br>PF00106.27: GO:0055114: oxidation-reduction process | biosynthetic-additional<br>MiBIG: BGC0001895.1, 11105, Streptomyces sp., Polyketide huanglongmycin |
| **26** | | PF01588.22 (Putative tRNA binding domain)<br>(This domain may perform a common function in tRNA aminoacylation)<br>PF01588.22: GO:0000049: tRNA binding | |
| **27** | | PF07676.14 (WD40-like Beta Propeller Repeat)<br>PF14684.8 (Tricorn protease C1 domain)<br>PF03572.20 (Peptidase family S41)<br>PF03572.20: GO:0008236: serine-type peptidase activity<br>PF03572.20: GO:0006508: proteolysis | biosynthetic-additional |
| **28** | | PF00916.22 (Sulfate permease family)<br>PF01740.23 (STAS domain)<br>PF00916.22: GO:0015116: sulfate transmembrane transporter activity | |

| | | |
|---|---|---|
| | | PF00916.22: GO:0008272: sulfate transport<br>PF00916.22: GO:0016021: integral component of membrane |
| **29** | | PF01244.23 (Membrane dipeptidase (Peptidase family M19))<br>PF01244.23: GO:0070573: metallodipeptidase activity<br>PF01244.23: GO:0006508: proteolysis |
| **30** | | PF02574.18 (Homocysteine S-methyltransferase) |
| **31** | | - |
| **32** | | - |

Table S4.3. antiSMASH annotations of contig gb1_c01994, detected in GCF 02410 and CC A

| Gene | SMCOG | PFAM & GO | Notes |
|---|---|---|---|
| **1** | SMCOG1002:AMP-dependent synthetase and ligase | PF00501.30 (AMP-binding enzyme)<br>PF13193.8 (AMP-binding enzyme C-terminal domain) | Biosynthetic<br>MiBIG: BGC0000649.1, fadD, Streptomyces griseus, Terpene, carotenoid |
| **2** | SMCOG1119:halogenase | PF04820.16 (Tryptophan halogenase) | Biosynthetic<br>MiBIG: BGC0001831.1, MXAN_6635, Myxococcus xanthus, Polyketide, alkylpyrones |
| **3** | | PF04055.23 (Radical SAM superfamily)<br>PF04055.23: GO:0003824: catalytic activity<br>PF04055.23: GO:0051536: iron-sulfur cluster binding | biosynthetic-additional |
| **4** | SMCOG1034:cytochrome P450 | PF00067.24 (Cytochrome P450)<br>PF00067.24: GO:0005506: iron ion binding<br>PF00067.24: GO:0016705: oxidoreductase activity, acting on paired donors, with incorporation or reduction of molecular oxygen<br>PF00067.24: GO:0020037: heme binding | biosynthetic-additional<br>p450 (11..429): active site cysteine inconclusive<br>MiBIG:BGC0001811.1, tri11, Fusarium asiaticum, Terpene, trichodiene-11-one |

| | | | |
|---|---|---|---|
| | | PF00067.24: GO:0055114: oxidation-reduction process | |
| **5** | SMCOG1034:cytochrome P450 | PF00067.24 (Cytochrome P450)<br>PF00067.24: GO:0005506: iron ion binding<br>PF00067.24: GO:0016705: oxidoreductase activity, acting on paired donors, with incorporation or reduction of molecular oxygen<br>PF00067.24: GO:0020037: heme binding<br>PF00067.24: GO:0055114: oxidation-reduction process | biosynthetic-additional<br>p450 (6..425): active site cysteine inconclusive<br>MiBIG: BGC0000675.1, SSCG_3690, Streptomyces clavuligerus ATCC 27064, Terpene, (+)-T-muurolol |
| **6** | SMCOG1034:cytochrome P450 | PF00067.24 (Cytochrome P450)<br>PF00067.24: GO:0005506: iron ion binding<br>PF00067.24: GO:0016705: oxidoreductase activity, acting on paired donors, with incorporation or reduction of molecular oxygen<br>PF00067.24: GO:0020037: heme binding<br>PF00067.24: GO:0055114: oxidation-reduction process | biosynthetic-additional<br>p450 (2..402): active site cysteine inconclusive<br>MiBIG: BGC0001910.1, cldC, Streptomyces cyslabdanicus, Terpene, cyslabdan |
| **7** | | - | |
| **8** | | PF01242.21 (6-pyruvoyl tetrahydropterin synthase) | |
| **9** | SMCOG1168:O-succinylhomoserine sulfhydrylase | PF01053.22 (Cys/Met metabolism PLP-dependent enzyme)<br>PF01053.22: GO:0030170: pyridoxal phosphate binding<br>PF01053.22: GO:0019346: transsulfuration | biosynthetic-additional |
| **10** | | PF05063.16 (MT-A70) | |
| **11** | | PF01957.20 (NfeD-like C-terminal, partner-binding) | |
| **12** | | PF01145.27 (SPFH domain / Band 7 family)<br>PF16200.7 (C-terminal region of band_7) | |

Table S4.4. antiSMASH annotations of contig gb5_2_c00041, detected in GCF 2402 and CC E

| Gene | SMCOG | PFAM & GO | Notes |
|---|---|---|---|

| | | | |
|---|---|---|---|
| **1** | SMCOG1210:glycerol kinase | PF00370.23 (FGGY family of carbohydrate kinases, N-terminal domain)<br>PF02782.18 (FGGY family of carbohydrate kinases, C-terminal domain)<br>PF00370.23: GO:0016773: phosphotransferase activity, alcohol group as acceptor<br>PF00370.23: GO:0005975: carbohydrate metabolic process<br>PF02782.18: GO:0016773: phosphotransferase activity, alcohol group as acceptor<br>PF02782.18: GO:0005975: carbohydrate metabolic process | **biosynthetic-additional** |
| **2** | | - | |
| **3** | | PF01259.20 (SAICAR synthetase/ligase) | |
| **4** | | - | |
| **5** | | PF05721.15 (Phytanoyl-CoA dioxygenase (PhyH)) | |
| **6** | | PF13360.8 (PQQ-like domain) | |
| **7** | | PF13360.8 (PQQ-like domain) | |
| **8** | | PF00501 (AMP-binding) | **biosynthetic-additional** |
| **9** | | PF13173.8 (AAA domain, ATPases)<br>PF13635.8 (Domain of unknown function (DUF4143)) | |
| **10** | SMCOG1017:aldehyde dehydrogenase | PF00171.24 (Aldehyde dehydrogenase family)<br>PF00171.24: GO:0016491: oxidoreductase activity<br>PF00171.24: GO:0055114: oxidation-reduction process | **biosynthetic-additional** |
| **11** | | PF00465.21 (Iron-containing alcohol dehydrogenase)<br>PF00465.21: GO:0016491: oxidoreductase activity<br>PF00465.21: GO:0046872: metal ion binding<br>PF00465.21: GO:0055114: oxidation-reduction process | |
| **12** | | PF13360.8 (PQQ-like domain) | |

| | | | |
|---|---|---|---|
| **13** | | PF04055.23 (Radical SAM superfamily)<br>PF04055.23: GO:0003824: catalytic activity<br>PF04055.23: GO:0051536: iron-sulfur cluster binding | **biosynthetic-additional** |
| **14** | SMCOG1051:TonB-dependent siderophore receptor | PF07715.17 (TonB-dependent Receptor Plug Domain)<br>PF00593.26 (TonB dependent receptor)<br>TIGR01783 (TonB-siderophor: TonB-dependent siderophore receptor) | **transport** |
| **15** | | PF14535.8 (AMP-binding enzyme C-terminal domain) | **Biosynthetic**<br>**MiBIG: BGC0000889.1, fevW, Streptomyces sp. WK-5344, Other, bagremycins** |
| **16** | SMCOG1119:halogenase | PF01494.21 (FAD binding domain)<br>PF01494.21: GO:0071949: FAD binding | **Biosynthetic**<br>**MiBIG: BGC0001366.1, orf12, Streptomyces sp. FJS31-2, Polyketide, zunyimycin** |
| **17** | SMCOG1045:glycosyl transferase group 1 | PF13439.8 (Glycosyltransferase Family 4)<br>PF00534.22 (Glycosyl transferases group 1)<br>PF00534.22: GO:0016757: transferase activity, transferring glycosyl groups | **biosynthetic-additional** |
| **18** | | | **-** |
| **19** | | PF03061.24 (Thioesterase superfamily) | |
| **20** | SMCOG1000:ABC transporter ATP-binding protein | PF00005.29 (ABC transporter)<br>PF00005.29: GO:0005524: ATP binding | **Transport**<br>**MiBIG: BGC0001526.1, brtI, Synechocystis salina LEGE 06099, Other, bartolosides** |
| **21** | | PF12704.9 (MacB-like periplasmic core domain)<br>PF02687.23 (FtsX-like permease family)<br>PF02687.23: GO:0016020: membrane | |
| **22** | | PF13360.8 (PQQ-like domain) | |

| | | |
|---|---|---|
| **23** | | PF13360.8 (PQQ-like domain) |
| **24** | | PF16868.7 (NMT1-like family, TAXI family of the substrate-binding proteins) |
| **25** | | PF06808.14 (Tripartite ATP-independent periplasmic transporter, DctM component)<br>TIGR02123 (TRAP_fused: TRAP transporter, 4TM/12TM fusion protein) |
| **26** | | PF16332.7 (Domain of unknown function (DUF4962))<br>PF07940.15 (Heparinase II/III-like protein)<br>PF07940.15: GO:0016829: lyase activity |
| **27** | | - |
| **28** | | PF00884.25 (Sulfatase)<br>PF00884.25: GO:0008484: sulfuric ester hydrolase activity |
| **29** | | PF06134.13 (L-rhamnose isomerase (RhaA))<br>TIGR01748 (rhaA: L-rhamnose isomerase) |
| **30** | | PF05721.15 (Phytanoyl-CoA dioxygenase (PhyH)) |
| **31** | | PF05721.15 (Phytanoyl-CoA dioxygenase (PhyH)) |
| **32** | | - |

Table S4.5. antiSMASH annotations of contig Pf12_c00284, detected in GCF 2043 and CC K

| Gene | SMCOG | PFAM & GO | Notes |
|---|---|---|---|
| **1** | SMCOG1079:oxidoreductase | PF01408.24 (Oxidoreductase family, NAD-binding Rossmann fold)<br>PF02894.19 (Oxidoreductase family, C-terminal alpha/beta domain)<br>PF01408.24: GO:0016491: oxidoreductase activity | biosynthetic-additional |
| **2** | SMCOG1079:oxidoreductase | PF01408.24 (Oxidoreductase family, NAD-binding Rossmann fold)<br>PF02894.19 (Oxidoreductase family, C-terminal alpha/beta domain) | biosynthetic-additional |

| | | | |
|---|---|---|---|
| | | PF01408.24: GO:0016491: oxidoreductase activity | |
| **3** | | - | |
| **4** | | PF01220.21 (Dehydroquinase class II)<br>TIGR01088 (aroQ: 3-dehydroquinate dehydratase, type II)<br>PF01220.21: GO:0003855: 3-dehydroquinate dehydratase activity | |
| **5** | SMCOG1056: DegT/DnrJ/EryC1/StrS aminotransferase | PF01041.19 (DegT/DnrJ/EryC1/StrS aminotransferase family) | biosynthetic-additional<br>MiBIG: BGC0000719.1, alloG, Streptoalloteichus hindustanus, Saccharide, tobramycin |
| **6** | | - | |
| **7** | | PF09924.11 (Phosphatidylglycerol lysyltransferase, C-terminal) (transfer of a lysyl group from L-lysyl-tRNA(Lys) to membrane-bound phosphatidylglycerol (PG)) | |
| **8** | | PF04263.18 (Thiamin pyrophosphokinase, catalytic domain)<br>PF04263.18: GO:0004788: thiamine diphosphokinase activity<br>PF04263.18: GO:0005524: ATP binding<br>PF04263.18: GO:0009229: thiamine diphosphate biosynthetic process | |
| **9** | SMCOG1081:cysteine synthase | PF00291.27 (Pyridoxal-phosphate dependent enzyme) | biosynthetic-additional |
| **10** | | - | |
| **11** | | PF00571.30 (CBS domain, bind ligands with an adenosyl group, AMP, ATP and S-AdoMe) | (regulatory) |
| **12** | | PF00571.30 (CBS domain, bind ligands with an adenosyl group, AMP, ATP and S-AdoMe) | (regulatory) |
| **13** | | PF01558.20 (Pyruvate ferredoxin/flavodoxin oxidoreductase) | |

| | | | |
|---|---|---|---|
| | | PF01855.21 (Pyruvate flavodoxin/ferredoxin oxidoreductase, thiamine diP-bdg)<br>PF17147.6 (Pyruvate:ferredoxin oxidoreductase core domain II)<br>TIGR03710 (OAFO_sf: 2-oxoacid:acceptor oxidoreductase, alpha subunit)<br>PF01558.20: GO:0016903: oxidoreductase activity, acting on the aldehyde or oxo group of donors<br>PF01558.20: GO:0055114: oxidation-reduction process<br>PF01855.21: GO:0016491: oxidoreductase activity<br>PF01855.21: GO:0055114: oxidation-reduction process | |
| **14** | | PF02775.23 (Thiamine pyrophosphate enzyme, C-terminal TPP binding domain)<br>TIGR02177 (PorB_KorB: 2-oxoacid:acceptor oxidoreductase, beta subunit, pyruvate/2-ketoisovalerate family)<br>PF02775.23: GO:0003824: catalytic activity<br>PF02775.23: GO:0030976: thiamine pyrophosphate binding | MiBIG: BGC0000131.1, dox5, Streptomyces sp., Polyketide, pyrrolomycin |
| **15** | | PF04951.15 (D-aminopeptidase) | |
| **16** | | PF13385.8 (Concanavalin A-like lectin/glucanases superfamily) | |
| **17** | | PF00465.21 (Iron-containing alcohol dehydrogenase)<br>PF00465.21: GO:0016491: oxidoreductase activity<br>PF00465.21: GO:0046872: metal ion binding<br>PF00465.21: GO:0055114: oxidation-reduction process | MiBIG: BGC0000897.1, dhpG, Streptomyces luridus, Other, dehydrophos |
| **18** | | PF13360.8 (PQQ-like domain)<br>PF13570.8 (PQQ-like domain)<br>PF13570.8 (PQQ-like domain) | |
| **19** | | PF01947.18 (p-hydroxybenzoic acid synthase (chorismate lyase), DUF98) | biosynthetic-additional<br>MiBIG: BGC0000891.1, pl2ta16_1240/bmp6, Pseudoalteromonas |

| | | | |
|---|---|---|---|
| | | | luteoviolacea, Other (Aminocoumarin), pentabromopseudilin |
| **20** | | PF04055.23 (Radical SAM superfamily)<br>PF06969.18 (HemN C-terminal domain)<br>PF04055.23: GO:0003824: catalytic activity<br>PF04055.23: GO:0051536: iron-sulfur cluster binding | biosynthetic-additional |
| **21** | | PF14535.8 (AMP-binding enzyme C-terminal domain) | Biosynthetic<br>MiBIG: BGC0000889.1, fevW, Streptomyces sp., Other, bagremycins |
| **22** | SMCOG1119:halogenase | PF04820.16 (Tryptophan halogenase) | Biosynthetic<br>MiBIG: BGC0001831.1,, MXAN_6635, Myxococcus xanthus, Polyketide, alkylpyrone |
| **23** | | PF16868.7 (NMT1-like family, TAXI family of the substrate-binding proteins)<br>TIGR02122 (TRAP_TAXI: TRAP transporter solute receptor, TAXI family) | |
| **24** | | PF05193.23 (Peptidase M16 inactive domain)<br>PF08367.13 (Peptidase M16C associated)<br>PF08367.13: GO:0006508: proteolysis | biosynthetic-additional |
| **25** | | PF00712.21 (DNA polymerase III beta subunit, N-terminal domain)<br>PF02767.18 (DNA polymerase III beta subunit, central domain)<br>PF02768.17 (DNA polymerase III beta subunit, C-terminal domain)<br>TIGR00663 (dnan: DNA polymerase III, beta subunit)<br>PF00712.21: GO:0003677: DNA binding<br>PF00712.21: GO:0003887: DNA-directed DNA polymerase activity<br>PF00712.21: GO:0008408: 3'-5' exonuclease activity<br>PF00712.21: GO:0006260: DNA replication<br>PF00712.21: GO:0009360: DNA polymerase III complex | |

| | | | |
|---|---|---|---|
| | | PF02767.18: GO:0003887: DNA-directed DNA polymerase activity<br>PF02767.18: GO:0008408: 3'-5' exonuclease activity<br>PF02767.18: GO:0006260: DNA replication<br>PF02767.18: GO:0009360: DNA polymerase III complex<br>PF02768.17: GO:0003677: DNA binding<br>PF02768.17: GO:0003887: DNA-directed DNA polymerase activity<br>PF02768.17: GO:0008408: 3'-5' exonuclease activity<br>PF02768.17: GO:0006260: DNA replication<br>PF02768.17: GO:0009360: DNA polymerase III complex | |
| **26** | SMCOG1272:TPR repeat-containing protein | PF13432.8 (Tetratricopeptide repeat)<br>PF00515.30 (Tetratricopeptide repeat)<br>PF13432.8 (Tetratricopeptide repeat)<br>PF00515.30: GO:0005515: protein binding | other |
| **27** | | PF11412.10 (Disulphide bond corrector protein DsbC - transmembrane electron transporter)<br>PF02683.17 (Cytochrome C biogenesis protein transmembrane region)<br>PF03190.17 (Protein of unknown function, DUF255)<br>PF02683.17: GO:0017004: cytochrome complex assembly<br>PF02683.17: GO:0055114: oxidation-reduction process<br>PF02683.17: GO:0016020: membrane | |
| **28** | | PF08665.14 (PglZ domain, putative phosphatase) | |
| **29** | SMCOG1261:hypothetical protein | PF01565.25 (FAD binding domain)<br>PF02913.21 (FAD linked oxidases, C-terminal domain)<br>PF01565.25: GO:0016491: oxidoreductase activity<br>PF01565.25: GO:0050660: flavin adenine dinucleotide binding<br>PF01565.25: GO:0055114: oxidation-reduction process<br>PF02913.21: GO:0003824: catalytic activity<br>PF02913.21: GO:0050660: flavin adenine dinucleotide binding | other |

| | | | |
|---|---|---|---|
| **30** | | - | |
| **31** | | - | |
| **32** | SMCOG1261:hypothetical protein | PF01565.25 (FAD binding domain)<br>PF02913.21 (FAD linked oxidases, C-terminal domain)<br>PF01565.25: GO:0016491: oxidoreductase activity<br>PF01565.25: GO:0050660: flavin adenine dinucleotide binding<br>PF01565.25: GO:0055114: oxidation-reduction process<br>PF02913.21: GO:0003824: catalytic activity<br>PF02913.21: GO:0050660: flavin adenine dinucleotide binding | other |
| **33** | | PF13385.8 (Concanavalin A-like lectin/glucanases superfamily) | |
| **34** | | - | |
| **35** | SMCOG1049:AcrB/AcrD/AcrF family protein | PF00873.21 (AcrB/AcrD/AcrF family)<br>PF00873.21: GO:0022857: transmembrane transporter activity<br>PF00873.21: GO:0055085: transmembrane transport<br>PF00873.21: GO:0016020: membrane | transport |
| **36** | | PF00873.21 (AcrB/AcrD/AcrF family)<br>PF00873.21: GO:0022857: transmembrane transporter activity<br>PF00873.21: GO:0055085: transmembrane transport<br>PF00873.21: GO:0016020: membrane | |

Table S4.6. antiSMASH annotations of contig Pf11_c00818, detected in GCF 2559 and CC H

| Gene | SMCOG | PFAM & GO | Notes |
|---|---|---|---|
| **1** | | PF03993.14 (Domain of Unknown Function (DUF349)) | |
| **2** | | PF00160.23 (Cyclophilin type peptidyl-prolyl cis-trans isomerase/CLD)<br>PF00160.23: GO:0003755: peptidyl-prolyl cis-trans isomerase activity<br>PF00160.23: GO:0000413: protein peptidyl-prolyl isomerization | |

| | | | |
|---|---|---|---|
| **3** | | PF00160.23 (Cyclophilin type peptidyl-prolyl cis-trans isomerase/CLD)<br>PF00160.23: GO:0003755: peptidyl-prolyl cis-trans isomerase activity<br>PF00160.23: GO:0000413: protein peptidyl-prolyl isomerization | |
| **4** | SMCOG1002:AMP-dependent synthetase and ligase | PF00550.27 (Phosphopantetheine attachment site)<br>PF00501.30 (AMP-binding enzyme)<br>PF01553.23 (Acyltransferase)<br>PF01553.23: GO:0016746: transferase activity, transferring acyl groups | Biosynthetic<br>MiBIG:BGC0001831.1, MXAN_6636, Myxococcus xanthus DK 1622, Polyketide, alkylpyrone |
| **5** | SMCOG1040:alcohol dehydrogenase | PF08240.14 (Alcohol dehydrogenase GroES-like domain)<br>PF00107.28 (Zinc-binding dehydrogenase)<br>PF00107.28: GO:0055114: oxidation-reduction process<br>PF08240.14: GO:0055114: oxidation-reduction process | biosynthetic-additional |
| **6** | | PF00583.27 (Acetyltransferase (GNAT) family)<br>PF00583.27: GO:0008080: N-acetyltransferase activity | |
| **7** | SMCOG1168:O-succinylhomoserine sulfhydrylase | PF01053.22 (Cys/Met metabolism PLP-dependent enzyme)<br>PF01053.22: GO:0030170: pyridoxal phosphate binding<br>PF01053.22: GO:0019346: transsulfuration | biosynthetic-additional<br>MiBIG: BGC0000783.1, metB, Xanthomonas oryzae, Saccharide, O-antigen |
| **8** | SMCOG1086:MATE efflux family protein | PF01554.20 (MatE)<br>TIGR00797 (matE: MATE efflux family protein)<br>PF01554.20: GO:0015297: antiporter activity<br>PF01554.20: GO:0042910: xenobiotic transmembrane transporter activity<br>PF01554.20: GO:0055085: transmembrane transport<br>PF01554.20: GO:0016020: membrane | transport |
| **9** | | PF00293.30 (NUDIX domain)<br>PF00293.30: GO:0016787: hydrolase activity | |
| **10** | | PF00521.22 (DNA gyrase/topoisomerase IV, subunit A) | |

| | | | |
|---|---|---|---|
| | | PF03989.15 (DNA gyrase C-terminal domain, beta-propeller)<br>PF00521.22: GO:0003677: DNA binding<br>PF00521.22: GO:0003918: DNA topoisomerase type II (double strand cut, ATP-hydrolyzing) activity<br>PF00521.22: GO:0005524: ATP binding<br>PF00521.22: GO:0006265: DNA topological change<br>PF03989.15: GO:0003677: DNA binding<br>PF03989.15: GO:0003916: DNA topoisomerase activity<br>PF03989.15: GO:0005524: ATP binding<br>PF03989.15: GO:0006265: DNA topological change | |
| **11** | | PF02518.28 (Histidine kinase-, DNA gyrase B-, and HSP90-like ATPase)<br>PF00204.27 (DNA gyrase B)<br>PF01751.24 (Toprim domain)<br>PF00986.23 (DNA gyrase B subunit, carboxyl terminus)<br>PF00204.27: GO:0003677: DNA binding<br>PF00204.27: GO:0003918: DNA topoisomerase type II (double strand cut, ATP-hydrolyzing) activity<br>PF00204.27: GO:0005524: ATP binding<br>PF00204.27: GO:0006265: DNA topological change<br>PF00986.23: GO:0003677: DNA binding<br>PF00986.23: GO:0003918: DNA topoisomerase type II (double strand cut, ATP-hydrolyzing) activity<br>PF00986.23: GO:0005524: ATP binding<br>PF00986.23: GO:0006265: DNA topological change | |
| **12** | | PF13570.8 (PQQ-like domain)<br>PF13360.8 (PQQ-like domain) | |
| **13** | | PF04055.23 (Radical SAM superfamily)<br>PF04055.23: GO:0003824: catalytic activity<br>PF04055.23: GO:0051536: iron-sulfur cluster binding | biosynthetic-additional |
| **14** | SMCOG1119:halogenase | PF04820.16 (Tryptophan halogenase) | Biosynthetic |

| | | | |
|---|---|---|---|
| | | | MiBiG:BGC0000891,<br>pl2ta16_1236/bmp2,<br>Pseudoalteromonas luteoviolacea, Other (Aminocoumarin), pentabromopseudilin |
| **15** | | PF03061.24 (Thioesterase superfamily) | |
| **16** | | PF00501.30 (AMP-binding enzyme)<br>PF14535.8 (AMP-binding enzyme C-terminal domain) | Biosynthetic<br>MiBIG:BGC0000889.1, fevW,<br>Streptomyces sp. Other, bagremycins |
| **17** | | PF00903.27 (Glyoxalase/Bleomycin resistance protein/Dioxygenase superfamily) | |
| **18** | | PF01593.26 (Flavin containing amine oxidoreductase)<br>PF01593.26: GO:0016491: oxidoreductase activity<br>PF01593.26: GO:0055114: oxidation-reduction process | (biosynthetic) |
| **19** | | PF00034.23 (Cytochrome c)<br>PF00034.23: GO:0009055: electron transfer activity<br>PF00034.23: GO:0020037: heme binding | |
| **20** | | PF16868.7 (NMT1-like family)<br>TIGR02122 (TRAP_TAXI: TRAP transporter solute receptor, TAXI family) | |
| **21** | | PF06808.14 (Tripartite ATP-independent periplasmic transporter, DctM component)<br>TIGR02123 (TRAP_fused: TRAP transporter, 4TM/12TM fusion protein) | |
| **22** | | PF00860.22 (Permease family)<br>PF00860.22: GO:0022857: transmembrane transporter activity<br>PF00860.22: GO:0055085: transmembrane transport<br>PF00860.22: GO:0016020: membrane | |

| | | | |
|---|---|---|---|
| **23** | | PF07045.13 (Domain of unknown function (DUF1330)) | |
| **24** | | - | |
| **25** | | PF13173.8 (AAA domain)<br>PF13635.8 (Domain of unknown function (DUF4143)) | |
| **26** | | PF01597.21 (Glycine cleavage H-protein)<br>(H-protein shuttles the methylamine group of glycine) | |
| **27** | | - | |
| **28** | SMCOG1178:cobalamin synthesis protein/P47K family protein | PF02492.21 (CobW/HypB/UreG, nucleotide-binding domain) | biosynthetic-additional |

Table S4.7. antiSMASH annotations of contig DOM045_c00026, detected in GCF 1912 and CC R

| Gene | SMCOG | PFAM & GO | Notes |
|---|---|---|---|
| **1** | | - | |
| **2** | SMCOG1272:TPR repeat-containing protein | PF13414.8 (TPR repeat) | Other |
| **3** | SMCOG1255:cold-shock DNA-binding domain protein | PF00313.24 ('Cold-shock' DNA-binding domain)<br>PF00313.24: GO:0003676: nucleic acid binding | regulatory |
| **4** | | - | |
| **5** | SMCOG1000:ABC transporter ATP-binding protein | PF00005.29 (ABC transporter)<br>PF00005.29: GO:0005524: ATP binding | transport |
| **6** | SMCOG1096:polar amino acid ABC transporter, inner membrane | PF00528.24 (Binding-protein-dependent transport system inner membrane component)<br>TIGR01726 (HEQRo_perm_3TM: amino ABC transporter, permease protein, 3-TM region, His/Glu/Gln/Arg/opine family) | transport |

| | | | |
|---|---|---|---|
| | | PF00528.24: GO:0055085: transmembrane transport<br>PF00528.24: GO:0016020: membrane | |
| **7** | SMCOG1096:polar amino acid ABC transporter, inner membrane | PF00528.24 (Binding-protein-dependent transport system inner membrane component)<br>TIGR01726 (HEQRo_perm_3TM: amino ABC transporter, permease protein, 3-TM region, His/Glu/Gln/Arg/opine family)<br>PF00528.24: GO:0055085: transmembrane transport<br>PF00528.24: GO:0016020: membrane | transport |
| **8** | SMCOG1091:glutamine-binding lipoprotein glnH | PF00497.22 (Bacterial extracellular solute-binding proteins, family 3) | biosynthetic-additional |
| **9** | | PF05833.13 (NFACT N-terminal and middle domains)<br>PF05670.15 (NFACT protein RNA binding domain) | |
| **10** | SMCOG1214:binding-protein-dependent transport systems | PF00528.24 (Binding-protein-dependent transport system inner membrane component)<br>PF00528.24 (Binding-protein-dependent transport system inner membrane component)<br>PF00528.24: GO:0055085: transmembrane transport<br>PF00528.24: GO:0016020: membrane | transport |
| **11** | | - | |
| **12** | | - | |
| **13** | | - | |
| **14** | | PF00941.23 (FAD binding domain in molybdopterin dehydrogenase)<br>PF03450.19 (CO dehydrogenase flavoprotein C-terminal domain)<br>PF00941.23: GO:0016491: oxidoreductase activity<br>PF00941.23: GO:0055114: oxidation-reduction process | |
| **15** | | PF00111.29 (2Fe-2S iron-sulfur cluster binding domain)<br>PF01799.22 ([2Fe-2S] binding domain) | |

| | | | |
|---|---|---|---|
| | | PF00111.29: GO:0009055: electron transfer activity<br>PF00111.29: GO:0051536: iron-sulfur cluster binding<br>PF01799.22: GO:0016491: oxidoreductase activity<br>PF01799.22: GO:0046872: metal ion binding<br>PF01799.22: GO:0055114: oxidation-reduction process | |
| **16** | SMCOG1293:aldehyde oxidase and xanthine dehydrogenase | PF01315.24 (Aldehyde oxidase and xanthine dehydrogenase, a/b hammerhead domain)<br>PF02738.20 (Molybdopterin-binding domain of aldehyde dehydrogenase)<br>PF02738.20: GO:0016491: oxidoreductase activity<br>PF02738.20: GO:0055114: oxidation-reduction process | biosynthetic-additional |
| **17** | | - | |
| **18** | SMCOG1002:AMP-dependent synthetase and ligase | PF16177.7 (Acetyl-coenzyme A synthetase N-terminus)<br>PF00501.30 (AMP-binding enzyme)<br>PF13193.8 (AMP-binding enzyme C-terminal domain) | biosynthetic |
| **19** | | PF01571.23 (Aminomethyltransferase folate-binding domain) | |
| **20** | | PF13510.8 (2Fe-2S iron-sulfur cluster binding domain)<br>PF07992.16 (Pyridine nucleotide-disulphide oxidoreductase)<br>PF17806.3 (Sarcosine oxidase A3 domain)<br>PF01571.23 (Aminomethyltransferase folate-binding domain)<br>PF08669.13 (Glycine cleavage T-protein C-terminal barrel domain)<br>PF07992.16: GO:0016491: oxidoreductase activity<br>PF07992.16: GO:0055114: oxidation-reduction process | |
| **21** | | PF04267.14 (Sarcosine oxidase, delta subunit family)<br>PF04267.14: GO:0008115: sarcosine oxidase activity<br>PF04267.14: GO:0046653: tetrahydrofolate metabolic process | |
| **22** | SMCOG1103:FAD dependent oxidoreductase | PF01266.26 (FAD dependent oxidoreductase)<br>PF01266.26: GO:0016491: oxidoreductase activity | (biosynthetic) |

| | | | |
|---|---|---|---|
| | | PF01266.26: GO:0055114: oxidation-reduction process | |
| **23** | | PF05721.15 (Phytanoyl-CoA dioxygenase (PhyH)) | |
| **24** | | PF12228.10 (Protein of unknown function (DUF3604)) | |
| **25** | SMCOG1268:mandelate racemase/muconate lactonizing enzyme | PF02746.18 (Mandelate racemase / muconate lactonizing enzyme, N-terminal domain)<br>PF13378.8 (Enolase C-terminal domain-like) | biosynthetic-additional |
| **26** | SMCOG1268:mandelate racemase/muconate lactonizing enzyme | PF13378.8 (Enolase C-terminal domain-like) | biosynthetic-additional |
| **27** | | PF01261.26 (Xylose isomerase-like TIM barrel) | |
| **28** | | PF03447.18 (Homoserine dehydrogenase, NAD binding domain)<br>PF00742.21 (Homoserine dehydrogenase)<br>PF01842.27 (ACT domain)<br>PF00742.21: GO:0006520: cellular amino acid metabolic process<br>PF00742.21: GO:0055114: oxidation-reduction process<br>PF03447.18: GO:0016491: oxidoreductase activity<br>PF03447.18: GO:0050661: NADP binding<br>PF03447.18: GO:0055114: oxidation-reduction process | |
| **29** | | PF00291.27 (Pyridoxal-phosphate dependent enzyme)<br>TIGR00260 (thrC: threonine synthase): [8:326](score: 320.5, e-value: 3.1e-96) | |
| **30** | | PF00696.30 (Amino acid kinase family)<br>PF01842.27 (ACT domain)<br>PF13840.8 (ACT domain)<br>TIGR00657 (asp_kinases: aspartate kinase)<br>TIGR00656 (asp_kin_monofn: aspartate kinase, monofunctional class) | |

| | | | |
|---|---|---|---|
| **31** | | PF02620.19 (Large ribosomal RNA subunit accumulation protein YceD) | |
| **32** | | PF01783.25 (Ribosomal L32p protein family)<br>TIGR01031 (rpmF_bact: ribosomal protein bL32)<br>PF01783.25: GO:0003735: structural constituent of ribosome<br>PF01783.25: GO:0006412: translation<br>PF01783.25: GO:0015934: large ribosomal subunit | |
| **33** | SMCOG1084:3-oxoacyl-(acyl carrier protein) synthase III | PF08545.12 (3-Oxoacyl-[acyl-carrier-protein (ACP)] synthase III)<br>PF08541.12 (3-Oxoacyl-[acyl-carrier-protein (ACP)] synthase III C terminal)<br>TIGR00747 (fabH: 3-oxoacyl-[acyl-carrier-protein] synthase III)<br>PF08545.12: GO:0004315: 3-oxoacyl-[acyl-carrier-protein] synthase activity<br>PF08545.12: GO:0006633: fatty acid biosynthetic process | biosynthetic-additional |
| **34** | SMCOG1021:malonyl CoA-acyl carrier protein transacylase | PF00698.23 (Acyl transferase domain)<br>TIGR00128 (fabD: malonyl CoA-acyl carrier protein transacylase) | biosynthetic-additional<br>PKS_AT (18..303)<br>(Methyl)Malonyl-CoA specificity inconclusive<br>found active site serine: True, scaffold matched GHGE: True |
| **35** | SMCOG1001:short-chain dehydrogenase/reductase SDR | PF13561.8 (Enoyl-(Acyl carrier protein) reductase)<br>TIGR01830 (3oxo_ACP_reduc: 3-oxoacyl-[acyl-carrier-protein] reductase) | biosynthetic-additional<br>PKS_KR (6..179)<br>KR domain putatively catalyzing D-configuration product formation<br>catalytic triad S,Y,N inconclusive |
| **36** | | PF00550.27 (Phosphopantetheine attachment site)<br>TIGR00517 (acyl_carrier: acyl carrier protein) | biosynthetic-additional |
| **37** | SMCOG1022:Beta-ketoacyl synthase | PF00109.28 (Beta-ketoacyl synthase, N-terminal domain)<br>PF02801.24 (Beta-ketoacyl synthase, C-terminal domain) | biosynthetic-additional<br>PKS_KS (6..411) |

| | | | |
|---|---|---|---|
| | | TIGR03150 (fabF: beta-ketoacyl-acyl-carrier-protein synthase II) | found active site cysteine: True,<br>scaffold matched GSSS: False<br>found active site histidines: True |
| **38** | | - | |
| **39** | SMCOG1013:aminotransferase class-III | PF00202.23 (Aminotransferase class-III)<br>PF00202.23: GO:0008483: transaminase activity<br>PF00202.23: GO:0030170: pyridoxal phosphate binding | biosynthetic-additional |
| **40** | | PF00903.27 (Glyoxalase/Bleomycin resistance protein/Dioxygenase superfamily) | |
| **41** | | PF12704.9 (MacB-like periplasmic core domain) | |
| **42** | | | |

Table S4.8. antiSMASH annotations of contig DOM007B_c00243, detected in GCF 1892 and CC C

| Gene | SMCOG | PFAM & GO | Notes |
|---|---|---|---|
| **1** | SMCOG1082:TonB-dependent siderophore receptor family | PF13620.8 (Carboxypeptidase regulatory-like domain)<br>PF00593.26 (TonB dependent receptor) | transport |
| **2** | SMCOG1045:glycosyl transferase group 1 | PF13439.8 (Glycosyltransferase Family 4)<br>PF00534.22 (Glycosyl transferases group 1)<br>PF00005.29 (ABC transporter)<br>PF00005.29: GO:0005524: ATP binding<br>PF00534.22: GO:0016757: transferase activity, transferring glycosyl groups | biosynthetic-additional<br>MiBIG: BGC0000487.1,cclG, Carnobacterium maltaromaticum, RiPP, carnocyclin |
| **3** | | - | |
| **4** | | PF04055.23 (Radical SAM superfamily)<br>PF04055.23: GO:0003824: catalytic activity<br>PF04055.23: GO:0051536: iron-sulfur cluster binding | biosynthetic-additional |

| | | | |
|---|---|---|---|
| **5** | SMCOG1119:halogenase | PF04820.16 (Tryptophan halogenase) | biosynthetic |
| **6** | | PF13279.8 (Thioesterase-like superfamily) | |
| **7** | | - | |
| **8** | MCOG1002:AMP-dependent synthetase and ligase | PF00501.30 (AMP-binding enzyme)<br>PF14535.8 (AMP-binding enzyme C-terminal domain) | biosynthetic |
| **9** | | PF13360.8 (PQQ-like domain) | |
| **10** | | PF12705.9 (PD-(D/E)XK nuclease superfamily) | |
| **11** | | PF00580.23 (UvrD/REP helicase N-terminal domain)<br>PF13361.8 (UvrD-like helicase C-terminal domain)<br>PF12705.9 (PD-(D/E)XK nuclease superfamily)<br>PF00580.23: GO:0005524: ATP binding<br>PF13361.8: GO:0005524: ATP binding<br>PF13361.8: GO:0016787: hydrolase activity | |
| **12** | | - | |
| **13** | | - | |
| **14** | | PF04055.23 (Radical SAM superfamily)<br>PF13186.8 (Iron-sulfur cluster-binding domain)<br>PF02810.17 (SEC-C motif)<br>TIGR03942 (sulfatase_rSAM: anaerobic sulfatase maturase)<br>TIGR04085 (rSAM_more_4Fe4S: radical SAM additional 4Fe4S-binding SPASM domain)<br>PF04055.23: GO:0003824: catalytic activity<br>PF04055.23: GO:0051536: iron-sulfur cluster binding | biosynthetic-additional |
| **15** | | PF03190.17 (Protein of unknown function, DUF255)<br>PF07221.13 (N-acylglucosamine 2-epimerase (GlcNAc 2-epimerase)) | |
| **16** | | PF00465.21 (Iron-containing alcohol dehydrogenase) | |

| | | | |
|---|---|---|---|
| | | PF00465.21: GO:0016491: oxidoreductase activity<br>PF00465.21: GO:0046872: metal ion binding<br>PF00465.21: GO:0055114: oxidation-reduction process | |
| **17** | | - | |
| **18** | SMCOG1000:ABC transporter ATP-binding protein | PF00005.29 (ABC transporter)<br>PF08402.12 (TOBE domain)<br>TIGR01187 (potA: polyamine ABC transporter, ATP-binding protein)<br>PF00005.29: GO:0005524: ATP binding<br>PF08402.12: GO:0005524: ATP binding<br>PF08402.12: GO:0022857: transmembrane transporter activity<br>PF08402.12: GO:0055085: transmembrane transport<br>PF08402.12: GO:0043190: ATP-binding cassette (ABC) transporter complex | Transport<br>MiBIG: BGC0001526.1, brtI, Synechocystis salina, Other, bartolosides |
| **19** | | PF13416.8 (Bacterial extracellular solute-binding protein) | |
| **20** | SMCOG1067:putative ABC transporter permease protein | PF00528.24 (Binding-protein-dependent transport system inner membrane component)<br>PF00528.24: GO:0055085: transmembrane transport<br>PF00528.24: GO:0016020: membrane | transport |
| **21** | SMCOG1069:putative ABC transporter permease protein | PF00528.24 (Binding-protein-dependent transport system inner membrane component)<br>PF00528.24: GO:0055085: transmembrane transport<br>PF00528.24: GO:0016020: membrane | transport |

Table S4.9. antiSMASH annotations of contig DOM045_c00004, detected in GCF 2002 and CC B

| Gene | SMCOG | PFAM & GO | Notes |
|---|---|---|---|
| **1** | | PF01751.24 (Toprim domain)<br>PF00986.23 (DNA gyrase B subunit, carboxyl terminus)<br>PF00986.23: GO:0003677: DNA binding | |

| | | | |
|---|---|---|---|
| | | PF00986.23: GO:0003918: DNA topoisomerase type II (double strand cut, ATP-hydrolyzing) activity<br>PF00986.23: GO:0005524: ATP binding<br>PF00986.23: GO:0006265: DNA topological change | |
| **2** | | - | |
| **3** | | PF03099.21 (Biotin/lipoate A/B protein ligase family)<br>PF03099.21: GO:0006464: cellular protein modification process | |
| **4** | SMCOG1139:aminotransferase class V | PF00266.21 (Aminotransferase class-V) | biosynthetic-additional |
| **5** | | PF01597.21 (Glycine cleavage H-protein) | |
| **6** | SMCOG1178:cobalamin synthesis protein/P47K family protein | PF02492.21 (CobW/HypB/UreG, nucleotide-binding domain): | biosynthetic-additional |
| **7** | | - | |
| **8** | | - | |
| **9** | | PF04055.23 (Radical SAM superfamily)<br>PF04055.23: GO:0003824: catalytic activity<br>PF04055.23: GO:0051536: iron-sulfur cluster binding | biosynthetic-additional |
| **10** | SMCOG1182:Polyprenyl synthetase | PF01976.19 (Protein of unknown function DUF116)<br>PF00348.19 (Polyprenyl synthetase)<br>PF00348.19: GO:0008299: isoprenoid biosynthetic process | biosynthetic-additional |
| **11** | | PF00432.23 (Prenyltransferase and squalene oxidase repeat)<br>PF00432.23: GO:0003824: catalytic activity | |
| **12** | | PF13249.8 (Squalene-hopene cyclase N-terminal domain) | Biosynthetic (terpene_cyclase) |
| **13** | | PF03235.16 (Protein of unknown function DUF262) | |

| | | | |
|---|---|---|---|
| **14** | | PF18735.3 (RiboL-PSP-HEPN) | |
| **15** | | PF00465.21 (Iron-containing alcohol dehydrogenase)<br>PF00465.21: GO:0016491: oxidoreductase activity<br>PF00465.21: GO:0046872: metal ion binding<br>PF00465.21: GO:0055114: oxidation-reduction process | |
| **16** | | PF13570.8 (PQQ-like domain)<br>PF01011.23 (PQQ enzyme repeat) | |
| **17** | | PF05685.14 (Putative restriction endonuclease) | |
| **18** | | PF04055.23 (Radical SAM superfamily)<br>PF04055.23: GO:0003824: catalytic activity<br>PF04055.23: GO:0051536: iron-sulfur cluster binding | biosynthetic-additional |
| **19** | | - | |
| **20** | SMCOG1119:halogenase | PF04820.16 (Tryptophan halogenase) | Biosynthetic<br>MiBIG: BGC0000890.1, bmp2, Pseudoalteromonas phenolica, Other (Aminocoumarin), pentabromopseudilin |
| **21** | | PF03061.24 (Thioesterase superfamily) | |
| **22** | SMCOG1002:AMP-dependent synthetase and ligase | PF00501.30 (AMP-binding enzyme)<br>PF14535.8 (AMP-binding enzyme C-terminal domain) | biosynthetic |
| **23** | | PF00903.27 (Glyoxalase/Bleomycin resistance protein/Dioxygenase superfamily) | |
| **24** | | PF16868.7 (NMT1-like family)<br>TIGR02122 (TRAP_TAXI: TRAP transporter solute receptor, TAXI family) | |
| **25** | | PF06808.14 (Tripartite ATP-independent periplasmic transporter, DctM component) | |

| | | | |
|---|---|---|---|
| | | TIGR02123 (TRAP_fused: TRAP transporter, 4TM/12TM fusion protein) | |
| **26** | | PF04174.15 (A circularly permuted ATPgrasp) | |
| **27** | | PF04168.14 (A predicted alpha-helical domain with a conserved ER motif.) | |
| **28** | | PF00909.23 (Ammonium Transporter Family)<br>PF00909.23: GO:0008519: ammonium transmembrane transporter activity<br>PF00909.23: GO:0015696: ammonium transport<br>PF00909.23: GO:0016020: membrane | |
| **29** | | PF03641.16 (Possible lysine decarboxylase)<br>TIGR00730 (TIGR00730: TIGR00730 family protein) | |
| **30** | | PF01740.23 (STAS (Sulphate Transporter and AntiSigma factor antagonist) domain,<br>TIGR00377 (ant_ant_sig: anti-anti-sigma factor) | In ecoli shown to interact with acyl carrier protein (ACP)) |
| **31** | | PF13581.8 (Histidine kinase-like ATPase domain) | |
| **32** | | PF01740.23 (STAS domain)<br>TIGR00377 (ant_ant_sig: anti-anti-sigma factor) | |
| **33** | | PF13581.8 (Histidine kinase-like ATPase domain) | |
| **34** | SMCOG1054:hypothetical protein | PF00672.27 (HAMP domain)<br>PF07228.14 (Stage II sporulation protein E (SpoIIE))<br>PF00672.27: GO:0007165: signal transduction<br>PF00672.27: GO:0016021: integral component of membrane<br>PF07228.14: GO:0016791: phosphatase activity | Other |
| **35** | | - | |
| **36** | SMCOG1122:ATP-dependent RNA helicase | PF00270.31 (DEAD/DEAH box helicase)<br>PF00271.33 (Helicase conserved C-terminal domain) | |

| | | | |
|---|---|---|---|
| | | PF16124.7 (RecQ zinc-binding)<br>PF09382.12 (RQC domain)<br>PF00570.25 (HRDC domain)<br>TIGR01389 (recQ: ATP-dependent DNA helicase RecQ)<br>PF00270.31: GO:0003676: nucleic acid binding<br>PF00270.31: GO:0005524: ATP binding<br>PF00570.25: GO:0003676: nucleic acid binding<br>PF09382.12: GO:0043138: 3'-5' DNA helicase activity<br>PF09382.12: GO:0006260: DNA replication<br>PF09382.12: GO:0006281: DNA repair | |
| **37** | | PF13360.8 (PQQ-like domain) | |
| **38** | | PF02190.18 (ATP-dependent protease La (LON) substrate-binding domain) | |
| **39** | SMCOG1036:alpha/beta hydrolase fold protein | PF00561.22 (alpha/beta hydrolase fold) | biosynthetic-additional |
| **40** | | - | |
| **41** | | PF16313.7 (Met-zincin, This domain carries the highly characteristic met-zincin mmotif HExxHxxGxxH, the extended zinc-binding domain of metallopeptidases.) | |
| **42** | | | |

Table S4.10. antiSMASH annotations of contig DOM016_c00717, detected in GCF 2039 and CC B

| Gene | SMCOG | PFAM & GO | Notes |
|---|---|---|---|
| **1** | | - | |
| **2** | | PF04306.15 (Protein of unknown function (DUF456)) | |
| **3** | | PF10041.11 (Uncharacterized conserved protein (DUF2277)) | |

| | | | |
|---|---|---|---|
| **4** | SMCOG1268:mandelate racemase/muconate lactonizing enzyme | PF02746.18 (Mandelate racemase / muconate lactonizing enzyme, N-terminal domain)<br>PF13378.8 (Enolase C-terminal domain-like) | biosynthetic-additional |
| **5** | | PF13709.8 (Domain of unknown function (DUF4159)) | |
| **6** | | - | |
| **7** | | PF12146.10 (Serine aminopeptidase, S33) | biosynthetic-additional (PF00561) |
| **8** | | PF07978.15 (NIPSNAP, hypothetical proteins) | |
| **9** | | PF00413.26 (Matrixin)<br>PF00413.26: GO:0004222: metalloendopeptidase activity<br>PF00413.26: GO:0008270: zinc ion binding<br>PF00413.26: GO:0006508: proteolysis<br>PF00413.26: GO:0031012: extracellular matrix | |
| **10** | | - | |
| **11** | | PF02190.18 (ATP-dependent protease La (LON) substrate-binding domain) | |
| **12** | | - | |
| **13** | | - | |
| **14** | SMCOG1054:hypothetical protein | PF00672.27 (HAMP domain)<br>PF07228.14 (Stage II sporulation protein E (SpoIIE))<br>PF00672.27: GO:0007165: signal transduction<br>PF00672.27: GO:0016021: integral component of membrane<br>PF07228.14: GO:0016791: phosphatase activity | MiBIG:BGC0000518.1, SSQG_07265, Streptomyces viridochromogenes, RiPP, informatipeptin |
| **15** | | PF13581.8 (Histidine kinase-like ATPase domain) | |
| **16** | | PF01740.23 (STAS domain)<br>TIGR00377 (ant_ant_sig: anti-anti-sigma factor) | |

| | | | |
|---|---|---|---|
| **17** | | PF00909.23 (Ammonium Transporter Family)<br>PF00909.23: GO:0008519: ammonium transmembrane transporter activity<br>PF00909.23: GO:0015696: ammonium transport<br>PF00909.23: GO:0016020: membrane | |
| **18** | SMCOG1082:TonB-dependent siderophore receptor family | PF00593.26 (TonB dependent receptor) | transport |
| **19** | | PF06808.14 (Tripartite ATP-independent periplasmic transporter, DctM component)<br>TIGR02123 (TRAP_fused: TRAP transporter, 4TM/12TM fusion protein) | |
| **20** | | PF16868.7 (NMT1-like family)<br>TIGR02122 (TRAP_TAXI: TRAP transporter solute receptor, TAXI family) | |
| **21** | | PF00903.27 (Glyoxalase/Bleomycin resistance protein/Dioxygenase superfamily) | |
| **22** | SMCOG1002:AMP-dependent synthetase and ligase | PF00501.30 (AMP-binding enzyme)<br>PF14535.8 (AMP-binding enzyme C-terminal domain) | Biosynthetic<br>MiBIG: BGC0000889.1, fevW, Streptomyces sp., Other, bagremycins |
| **23** | | PF03061.24 (Thioesterase superfamily) | |
| **24** | SMCOG1119:halogenase | PF01494.21 (FAD binding domain)<br>PF01494.21: GO:0071949: FAD binding | biosynthetic<br>MiBIG: BGC0001831.1, MXAN_6635, Myxococcus xanthus, Polyketide, alkylpyrones |
| **25** | | - | |
| **26** | | PF04055.23 (Radical SAM superfamily)<br>PF04055.23: GO:0003824: catalytic activity<br>PF04055.23: GO:0051536: iron-sulfur cluster binding | biosynthetic-additional |

| | | | |
|---|---|---|---|
| **27** | | PF05685.14 (Putative restriction endonuclease) | MiBIG: BGC0000479.1, AED99428.1, Planktothrix agardhii, RiPP, prenylagaramides |
| **28** | | PF13570.8 (PQQ-like domain) | |
| **29** | | PF00465.21 (Iron-containing alcohol dehydrogenase)<br>PF00465.21: GO:0016491: oxidoreductase activity<br>PF00465.21: GO:0046872: metal ion binding<br>PF00465.21: GO:0055114: oxidation-reduction process | |
| **30** | | PF13564.8 (DoxX-like family, uncharacterised transmembrane proteins known as DoxX) | |
| **31** | | - | |

Table S4.11. antiSMASH annotations of contig DOM044_c00402, detected in GCF 1983 and CC D

| Gene | SMCOG | PFAM & GO | Notes |
|---|---|---|---|
| **1** | | PF02366.20 (Dolichyl-phosphate-mannose-protein mannosyltransferase)<br>PF14559.8 (Tetratricopeptide repeat)<br>PF02366.20: GO:0000030: mannosyltransferase activity<br>PF02366.20: GO:0006493: protein O-linked glycosylation<br>PF02366.20: GO:0016020: membrane | |
| **2** | SMCOG1123:polyprenol-monophosphomannose synthase ppm1 | PF00535.28 (Glycosyl transferase family 2) | biosynthetic-additional |
| **3** | | PF09861.11 (Lactate racemase N-terminal domain)<br>PF09861.11: GO:0050043: lactate racemase activity | |
| **4** | | PF19501.1 (YetA-like protein)<br>PF07944.14 (Beta-L-arabinofuranosidase, GH127) | |

| | | | |
|---|---|---|---|
| **5** | SMCOG1079:oxidoreductase | PF01408.24 (Oxidoreductase family, NAD-binding Rossmann fold)<br>PF02894.19 (Oxidoreductase family, C-terminal alpha/beta domain)<br>PF01408.24: GO:0016491: oxidoreductase activity | biosynthetic-additional<br>MiBIG: BGC0000720.1, tbmC (putative dehydrogenase), Streptoalloteichus tenebrarius, Saccharide, tobramycin |
| **6** | SMCOG1079:oxidoreductase | PF01408.24 (Oxidoreductase family, NAD-binding Rossmann fold)<br>PF02894.19 (Oxidoreductase family, C-terminal alpha/beta domain)<br>PF01408.24: GO:0016491: oxidoreductase activity | biosynthetic-additional |
| **7** | | AMP-binding | biosynthetic-additional<br>MiBIG: BGC0000662.1, sgr_4243(AMP), Streptomyces griseus, Terpene, grixazone |
| **8** | | PF00171.24 (Aldehyde dehydrogenase family)<br>PF00171.24: GO:0016491: oxidoreductase activity<br>PF00171.24: GO:0055114: oxidation-reduction process | |
| **9** | | PF00465.21 (Iron-containing alcohol dehydrogenase)<br>PF00465.21: GO:0016491: oxidoreductase activity<br>PF00465.21: GO:0046872: metal ion binding<br>PF00465.21: GO:0055114: oxidation-reduction process | MiBIG: BGC0000807.1, snas_5663, Stackebrandtia nassauensis, Saccharide, phosphonoglycans |
| **10** | | PF13570.8 (PQQ-like domain) | |
| **11** | | PF04055.23 (Radical SAM superfamily)<br>PF04055.23: GO:0003824: catalytic activity<br>PF04055.23: GO:0051536: iron-sulfur cluster binding | biosynthetic-additional |
| **12** | | PF04255.16 (Protein of unknown function (DUF433)) | |
| **13** | | PF13360.8 (PQQ-like domain) | |
| **14** | SMCOG1051:TonB-dependent siderophore receptor | PF07715.17 (TonB-dependent Receptor Plug Domain)<br>PF00593.26 (TonB dependent receptor) | transport |

| | | | |
|---|---|---|---|
| | | TIGR01783 (TonB-siderophor: TonB-dependent siderophore receptor) | |
| **15** | | PF14535.8 (AMP-binding enzyme C-terminal domain) | biosynthetic |
| **16** | SMCOG1119:halogenase | PF01494.21 (FAD binding domain)<br>PF01494.21: GO:0071949: FAD binding | Biosynthetic<br>MiBIG:BGC0000837.1, vf_0841, Aliivibrio fischeri , Other, APE Vf |
| **17** | SMCOG1045:glycosyl transferase group 1 | PF13579.8 (Glycosyl transferase 4-like domain)<br>PF00534.22 (Glycosyl transferases group 1)<br>PF00534.22: GO:0016757: transferase activity, transferring glycosyl groups | biosynthetic-additional |
| **18** | | - | |
| **19** | | PF03061.24 (Thioesterase superfamily) | |
| **20** | SMCOG1000:ABC transporter ATP-binding protein | PF00005.29 (ABC transporter)<br>PF00005.29: GO:0005524: ATP binding | Transport<br>MiBIG: BGC0001526.1, brtI, Synechocystis salina, Other, bartolosides |
| **21** | | PF02687.23 (FtsX-like permease family)<br>PF02687.23: GO:0016020: membrane | |
| **22** | | PF13360.8 (PQQ-like domain) | |
| **23** | | PF16868.7 (NMT1-like family)<br>TIGR02122 (TRAP_TAXI: TRAP transporter solute receptor, TAXI family) | |
| **24** | | PF06808.14 (Tripartite ATP-independent periplasmic transporter, DctM component)<br>TIGR02123 (TRAP_fused: TRAP transporter, 4TM/12TM fusion protein) | |
| **25** | | - | |
| **26** | | PF01402.23 (Ribbon-helix-helix protein, copG family) | |

| | | | |
|---|---|---|---|
| | | PF03683.15 (Uncharacterised protein family (UPF0175))<br>PF01402.23: GO:0006355: regulation of transcription, DNA-templated | |
| **27** | SMCOG1059:acetyl-CoA carboxylase, carboxyl transferase | PF01039.24 (Carboxyl transferase domain) | biosynthetic-additional |
| **28** | SMCOG1189:L-carnitine dehydratase/bile acid-inducible | PF02515.19 (CoA-transferase family III)<br>PF02515.19: GO:0008410: CoA-transferase activity | biosynthetic-additional |
| **29** | | PF10282.11 (Lactonase, 7-bladed beta-propeller) | |
| **30** | | PF05721.15 (Phytanoyl-CoA dioxygenase (PhyH)) | |
| **31** | | PF01261.26 (Xylose isomerase-like TIM barrel) | |
| **32** | | PF05721.15 (Phytanoyl-CoA dioxygenase (PhyH)) | |

Table S4.12. antiSMASH annotations of contig Pf7_c00820, detected in GCF 2325 and CC F

| Gene | SMCOG | PFAM & GO | Notes |
|---|---|---|---|
| **1** | | - | |
| **2** | | PF13338.8 (Transcriptional regulator, AbiEi antitoxin) | |
| **3** | SMCOG1115:HAD-superfamily hydrolase, subfamily IA, variant | PF13419.8 (Haloacid dehalogenase-like hydrolase)<br>TIGR01549 (HAD-SF-IA-v1: HAD hydrolase, family IA, variant 1) | biosynthetic-additional |
| **4** | | - | |
| **5** | | PF01850.23 (PIN domain, ribonuclease toxic components of toxin-antitoxin (TA) systems) | |
| **6** | | - | |

| | | | |
|---|---|---|---|
| **7** | SMCOG1082:TonB-dependent siderophore receptor family | PF07715.17 (TonB-dependent Receptor Plug Domain)<br>PF00593.26 (TonB dependent receptor) | transport |
| **8** | SMCOG1017:aldehyde dehydrogenase | PF00171.24 (Aldehyde dehydrogenase family)<br>PF00171.24: GO:0016491: oxidoreductase activity<br>PF00171.24: GO:0055114: oxidation-reduction process | biosynthetic-additional |
| **9** | | PF00465.21 (Iron-containing alcohol dehydrogenase)<br>PF00465.21: GO:0016491: oxidoreductase activity<br>PF00465.21: GO:0046872: metal ion binding<br>PF00465.21: GO:0055114: oxidation-reduction process | MiBIG: BGC0000807.1, snas_5663, Stackebrandtia nassauensis, Saccharide, phosphonoglycans |
| **10** | | PF05721.15 (Phytanoyl-CoA dioxygenase (PhyH)) | |
| **11** | SMCOG1288:ABC transporter related protein | PF00664.25 (ABC transporter transmembrane region)<br>PF00005.29 (ABC transporter)<br>PF00005.29: GO:0005524: ATP binding<br>PF00664.25: GO:0005524: ATP binding<br>PF00664.25: GO:0042626: ATPase-coupled transmembrane transporter activity<br>PF00664.25: GO:0055085: transmembrane transport<br>PF00664.25: GO:0016021: integral component of membrane | Transport<br>MiBIG:BGC0000497.1, bhtT, Streptococcus ratti, RiPP, BHT-A |
| **12** | | - | |
| **13** | | PF01850.23 (PIN domain, ribonuclease toxic components of toxin-antitoxin (TA) systems) | |
| **14** | | PF01011.23 (PQQ enzyme repeat)<br>PF13360.8 (PQQ-like domain) | |
| **15** | | PF04055.23 (Radical SAM superfamily)<br>PF06969.18 (HemN C-terminal domain)<br>PF04055.23: GO:0003824: catalytic activity<br>PF04055.23: GO:0051536: iron-sulfur cluster binding | biosynthetic-additional |

| | | | |
|---|---|---|---|
| **16** | | PF13360.8 (PQQ-like domain) | |
| **17** | | PF13360.8 (PQQ-like domain) | |
| **18** | | PF01011.23 (PQQ enzyme repeat)<br>PF13360.8 (PQQ-like domain) | |
| **19** | SMCOG1051:TonB-dependent siderophore receptor | PF07715.17 (TonB-dependent Receptor Plug Domain)<br>PF00593.26 (TonB dependent receptor)<br>TIGR01783 (TonB-siderophor: TonB-dependent siderophore receptor) | transport |
| **20** | | PF14535.8 (AMP-binding enzyme C-terminal domain) | Biosynthetic<br>MiBIG:BGC0000889.1, fevW, Streptomyces sp., Other, bagremycins |
| **21** | SMCOG1119:halogenase | PF04820.16 (Tryptophan halogenase) | Biosynthetic<br>MiBIG:BGC0001366.1, orf12, Streptomyces sp, Polyketide, zunyimycin A |
| **22** | | PF11755.10 (Protein of unknown function (DUF3311)) | |
| **23** | | PF00474.19 (Sodium:solute symporter family)<br>PF00474.19: GO:0022857: transmembrane transporter activity<br>PF00474.19: GO:0055085: transmembrane transport<br>PF00474.19: GO:0016020: membrane | |
| **24** | SMCOG1045:glycosyl transferase group 1 | PF13439.8 (Glycosyltransferase Family 4)<br>PF00534.22 (Glycosyl transferases group 1)<br>PF00534.22: GO:0016757: transferase activity, transferring glycosyl groups | biosynthetic-additional |
| **25** | | PF13173.8 (AAA domain)<br>PF13635.8 (Domain of unknown function (DUF4143)) | |

| | | | |
|---|---|---|---|
| **26** | | - | |
| **27** | SMCOG1014:LysR family transcriptional regulator | PF00126.29 (Bacterial regulatory helix-turn-helix protein, lysR family)<br>PF03466.22 (LysR substrate binding domain)<br>PF00126.29: GO:0003700: DNA-binding transcription factor activity<br>PF00126.29: GO:0006355: regulation of transcription, DNA-templated | regulatory |
| **28** | SMCOG1179:hydrolase | PF08282.14 (haloacid dehalogenase-like hydrolase) | biosynthetic-additional |

Table S4.13. antiSMASH annotations of contig DOM43_c00257, detected in GCF 1884 and CC J

| Gene | SMCOG | PFAM & GO | Notes |
|---|---|---|---|
| **1** | | - | |
| **2** | | - | |
| **3** | SMCOG1274:gamma-glutamyltranspeptidase | PF01019.23 (Gamma-glutamyltranspeptidase)<br>TIGR00066 (g_glut_trans: gamma-glutamyltransferase) | biosynthetic-additional |
| **4** | | PF00465.21 (Iron-containing alcohol dehydrogenase)<br>PF00465.21: GO:0016491: oxidoreductase activity<br>PF00465.21: GO:0046872: metal ion binding<br>PF00465.21: GO:0055114: oxidation-reduction process | |
| **5** | | PF13570.8 (PQQ-like domain) | |
| **6** | | PF00465.21 (Iron-containing alcohol dehydrogenase)<br>PF00465.21: GO:0016491: oxidoreductase activity<br>PF00465.21: GO:0046872: metal ion binding<br>PF00465.21: GO:0055114: oxidation-reduction process | |
| **7** | | - | |
| **8** | | PF01850.23 (PIN domain) | |
| **9** | | PF00474.19 (Sodium:solute symporter family) | |

| | | | |
|---|---|---|---|
| | | PF00474.19: GO:0022857: transmembrane transporter activity<br>PF00474.19: GO:0055085: transmembrane transport<br>PF00474.19: GO:0016020: membrane | |
| **10** | | - | |
| **11** | | PF12796.9 (Ankyrin repeats (3 copies)) | |
| **12** | | - | |
| **13** | | - | |
| **14** | | PF04221.14 (RelB antitoxin)<br>TIGR02384 (RelB_DinJ: addiction module antitoxin, RelB/DinJ family) | |
| **15** | | PF15738.7 (Bacterial toxin of type II toxin-antitoxin system, YafQ)<br>TIGR02385 (RelE_StbE: addiction module toxin, RelE/StbE family) | |
| **16** | | PF16868.7 (NMT1-like family)<br>TIGR02122 (TRAP_TAXI: TRAP transporter solute receptor, TAXI family) | |
| **17** | | PF06808.14 (Tripartite ATP-independent periplasmic transporter, DctM component)<br>TIGR02123 (TRAP_fused: TRAP transporter, 4TM/12TM fusion protein) | |
| **18** | | PF04055.23 (Radical SAM superfamily)<br>PF04055.23: GO:0003824: catalytic activity<br>PF04055.23: GO:0051536: iron-sulfur cluster binding | biosynthetic-additional |
| **19** | SMCOG1119:halogenase | PF04820.16 (Tryptophan halogenase) | Biosynthetic<br>MiBIG:BGC0001333.1,abeX2, uncultured AB1650, alkaloid BE-54017 |
| **20** | | PF03061.24 (Thioesterase superfamily) | |

| | | | |
|---|---|---|---|
| **21** | MCOG1002:AMP-dependent synthetase and ligase | PF00501.30 (AMP-binding enzyme)<br>PF14535.8 (AMP-binding enzyme C-terminal domain) | biosynthetic |
| **22** | | PF13360.8 (PQQ-like domain) | |
| **23** | | - | |
| **24** | SMCOG1000:ABC transporter ATP-binding protein | PF00005.29 (ABC transporter)<br>PF00005.29: GO:0005524: ATP binding | transport |
| **25** | | - | |
| **26** | | PF06182.13 (ABC-2 family transporter protein) | |
| **27** | | PF06182.13 (ABC-2 family transporter protein) | |
| **28** | | PF03485.18 (Arginyl tRNA synthetase N terminal domain)<br>PF00750.21 (tRNA synthetases class I (R))<br>PF05746.17 (DALR anticodon binding domain)<br>PF03485.18: GO:0000166: nucleotide binding<br>PF03485.18: GO:0004814: arginine-tRNA ligase activity<br>PF03485.18: GO:0005524: ATP binding<br>PF03485.18: GO:0006420: arginyl-tRNA aminoacylation<br>PF03485.18: GO:0005737: cytoplasm<br>PF05746.17: GO:0004814: arginine-tRNA ligase activity<br>PF05746.17: GO:0005524: ATP binding<br>PF05746.17: GO:0006420: arginyl-tRNA aminoacylation | |
| **29** | | PF13669.8 (Glyoxalase/Bleomycin resistance protein/Dioxygenase superfamily)<br>TIGR03081 (metmalonyl_epim: methylmalonyl-CoA epimerase) | |
| **30** | | PF01048.22 (Phosphorylase superfamily)<br>TIGR01694 (MTAP: methylthioadenosine phosphorylase)<br>PF01048.22: GO:0003824: catalytic activity<br>PF01048.22: GO:0009116: nucleoside metabolic process | |

| | | | |
|---|---|---|---|
| **31** | SMCOG1276:PfkB domain protein | PF00294.26 (pfkB family carbohydrate kinase) | biosynthetic-additional |
| **32** | | PF01678.21 (Diaminopimelate epimerase)<br>TIGR00652 (DapF: diaminopimelate epimerase)<br>PF01678.21: GO:0008837: diaminopimelate epimerase activity<br>PF01678.21: GO:0009089: lysine biosynthetic process via diaminopimelate | MiBIG:BGC0001733.1, 6800, Pleurocapsa sp., RiPP, PcpA |
| **33** | | PF01894.19 (Uncharacterised protein family UPF0047)<br>TIGR00149 (TIGR00149_YjbQ: secondary thiamine-phosphate synthase enzyme) | |
| **34** | SMCOG1106:major facilitator transporter | PF07690.18 (Major Facilitator Superfamily)<br>PF07690.18: GO:0022857: transmembrane transporter activity<br>PF07690.18: GO:0055085: transmembrane transport | transport |
| **35** | | PF02517.18 (Type II CAAX prenyl endopeptidase Rce1-like)<br>PF02517.18: GO:0004222: metalloendopeptidase activity<br>PF02517.18: GO:0071586: CAAX-box protein processing<br>PF02517.18: GO:0016020: membrane | biosynthetic-additional (Abi) |
| **36** | SMCOG1220:bifunctional ornithine | PF01960.20 (ArgJ family)<br>TIGR00120 (ArgJ: glutamate N-acetyltransferase/amino-acid acetyltransferase) | biosynthetic-additional |
| **37** | SMCOG1063:argininosuccinate lyase/adenylosuccinate lyase | PF00206.22 (Lyase)<br>PF14698.8 (Argininosuccinate lyase C-terminal)<br>TIGR00838 (argH: argininosuccinate lyase) | biosynthetic-additional |
| **38** | SMCOG1291:argininosuccinate synthase | PF00764.21 (Arginosuccinate synthase)<br>TIGR00032 (argG: argininosuccinate synthase)<br>PF00764.21: GO:0004055: argininosuccinate synthase activity<br>PF00764.21: GO:0005524: ATP binding<br>PF00764.21: GO:0006526: arginine biosynthetic process | biosynthetic-additional |

| | | | |
|---|---|---|---|
| **39** | SMCOG1114:ornithine carbamoyltransferase | PF02729.23 (Aspartate/ornithine carbamoyltransferase, carbamoyl-P binding domain)<br>PF00185.26 (Aspartate/ornithine carbamoyltransferase, Asp/Orn binding domain)<br>TIGR00658 (orni_carb_tr: ornithine carbamoyltransferase)<br>PF00185.26: GO:0016597: amino acid binding<br>PF00185.26: GO:0016743: carboxyl- or carbamoyltransferase activity<br>PF00185.26: GO:0006520: cellular amino acid metabolic process<br>PF02729.23: GO:0016743: carboxyl- or carbamoyltransferase activity<br>PF02729.23: GO:0006520: cellular amino acid metabolic process | biosynthetic-additional |
| **40** | SMCOG1013:aminotransferase class-III | PF00202.23 (Aminotransferase class-III)<br>TIGR00707 (argD: transaminase, acetylornithine/succinylornithine family)<br>PF00202.23: GO:0008483: transaminase activity<br>PF00202.23: GO:0030170: pyridoxal phosphate binding | biosynthetic-additional<br>MiBIG:BGC0000855.1, ectB ( L-2,4-diaminobutyric acid acetyl transferase), Methylomicrobium kenyense, Other, ectoine |

Table S4.14. antiSMASH annotations of contig gb4_2_c00117, detected in GCF 2554 and CC I

| Gene | SMCOG | PFAM & GO | Notes |
|---|---|---|---|
| **1** | SMCOG1173:WD-40 repeat-containing protein | PF00400.34 (WD domain, G-beta repeat)<br>PF00400.34: GO:0005515: protein binding | |
| **2** | | - | |
| **3** | | PF02633.16 (Creatinine amidohydrolase) | |
| **4** | | PF00465.21 (Iron-containing alcohol dehydrogenase)<br>PF00465.21: GO:0016491: oxidoreductase activity<br>PF00465.21: GO:0046872: metal ion binding<br>PF00465.21: GO:0055114: oxidation-reduction process | |
| **5** | | PF01011.23 (PQQ enzyme repeat) | |

| | | | |
|---|---|---|---|
| **6** | | PF04055.23 (Radical SAM superfamily)<br>PF06969.18 (HemN C-terminal domain)<br>PF04055.23: GO:0003824: catalytic activity<br>PF04055.23: GO:0051536: iron-sulfur cluster binding | biosynthetic-additional |
| **7** | | PF13360.8 (PQQ-like domain) | |
| **8** | | PF01344.27 (Kelch motif)<br>TIGR04183 (Por_Secre_tail: Por secretion system C-terminal sorting domain)<br>PF01344.27: GO:0005515: protein binding | |
| **9** | | PF14535.8 (AMP-binding enzyme C-terminal domain) | Biosynthetic<br>MiBIG: BGC0000889.1, fevW, Streptomyces sp., other, bagremycins |
| **10** | SMCOG1119:halogenase | PF04820.16 (Tryptophan halogenase) | biosynthetic |
| **11** | | PF16868.7 (NMT1-like family)<br>TIGR02122 (TRAP_TAXI: TRAP transporter solute receptor, TAXI family) | |
| **12** | | PF06808.14 (Tripartite ATP-independent periplasmic transporter, DctM component)<br>TIGR02123 (TRAP_fused: TRAP transporter, 4TM/12TM fusion protein) | |
| **13** | | PF11755.10 (Protein of unknown function (DUF3311)) | |
| **14** | | PF00474.19 (Sodium:solute symporter family)<br>PF00474.19: GO:0022857: transmembrane transporter activity<br>PF00474.19: GO:0055085: transmembrane transport<br>PF00474.19: GO:0016020: membrane | |
| **15** | SMCOG1045:glycosyl transferase group 1 | PF13439.8 (Glycosyltransferase Family 4)<br>PF00534.22 (Glycosyl transferases group 1) | biosynthetic-additional |

| | | | |
|---|---|---|---|
| | | PF00534.22: GO:0016757: transferase activity, transferring glycosyl groups | |
| **16** | | - | |
| **17** | | PF03061.24 (Thioesterase superfamily) | |
| **18** | SMCOG1000:ABC transporter ATP-binding protein | PF00005.29 (ABC transporter)<br>PF00005.29: GO:0005524: ATP binding | Transport<br>MiBIG: BGC0001526.1,brtI, Synechocystis salina, other, bartolosides |
| **19** | | PF02687.23 (FtsX-like permease family)<br>PF02687.23: GO:0016020: membrane | |
| **20** | | PF00892.22 (EamA-like transporter family)<br>PF00892.22: GO:0016020: membrane<br>PF00892.22: GO:0016021: integral component of membrane | |
| **21** | SMCOG1122:ATP-dependent RNA helicase | PF00270.31 (DEAD/DEAH box helicase)<br>PF00271.33 (Helicase conserved C-terminal domain)<br>PF09369.12 (Domain of unknown function (DUF1998))<br>PF00270.31: GO:0003676: nucleic acid binding<br>PF00270.31: GO:0005524: ATP binding | |
| **22** | SMCOG1079:oxidoreductase | PF01408.24 (Oxidoreductase family, NAD-binding Rossmann fold)<br>PF02894.19 (Oxidoreductase family, C-terminal alpha/beta domain)<br>PF01408.24: GO:0016491: oxidoreductase activity | biosynthetic-additional |
| **23** | | PF01555.20 (DNA methylase)<br>PF01555.20: GO:0003677: DNA binding<br>PF01555.20: GO:0008170: N-methyltransferase activity<br>PF01555.20: GO:0006306: DNA methylation | |
| **24** | | PF01555.20 (DNA methylase)<br>PF01555.20: GO:0003677: DNA binding | |

| | | |
|---|---|---|
| | | PF01555.20: GO:0008170: N-methyltransferase activity<br>PF01555.20: GO:0006306: DNA methylation |
| **25** | | - |

Table S4.15. antiSMASH annotations of contig DOM026_c00344, detected in GCF 2138 and CC P

| Gene | SMCOG | PFAM & GO | Notes |
|---|---|---|---|
| **1** | | PF02687.23 (FtsX-like permease family)<br>PF12704.9 (MacB-like periplasmic core domain)<br>PF02687.23: GO:0016020: membrane | |
| **2** | SMCOG1000:ABC transporter ATP-binding protein | PF00005.29 (ABC transporter)<br>PF00005.29: GO:0005524: ATP binding | Transport<br>MiBIG: BGC0000563.1, venT, Streptomyces venezuelae , RiPP, venezuelin |
| **3** | SMCOG1112:sigma-54 dependent trancsriptional regulator | PF00072.26 (Response regulator receiver domain)<br>PF00158.28 (Sigma-54 interaction domain)<br>PF02954.21 (Bacterial regulatory protein, Fis family)<br>PF00072.26: GO:0000160: phosphorelay signal transduction system<br>PF00158.28: GO:0005524: ATP binding<br>PF00158.28: GO:0008134: transcription factor binding<br>PF00158.28: GO:0006355: regulation of transcription, DNA-templated<br>PF02954.21: GO:0043565: sequence-specific DNA binding | Regulatory<br>MiBIG:BGC0001589.1, RS61070, Burkholderia cenocepacia, Saccharide, exopolysaccharides |
| **4** | SMCOG1003:sensor histidine kinase | PF02518.28 (Histidine kinase-, DNA gyrase B-, and HSP90-like ATPase) | regulatory |
| **5** | | PF13202.8 (EF hand)<br>PF13202.8: GO:0005509: calcium ion binding | |
| **6** | | - | |
| **7** | | PF02604.21 (Antitoxin Phd_YefM, type II toxin-antitoxin system) | |

| | | | |
|---|---|---|---|
| | | TIGR01552 (phd_fam: prevent-host-death family protein) | |
| **8** | | PF01850.23 (PIN domain, ribonuclease toxic components of toxin-antitoxin (TA) systems) | |
| **9** | SMCOG1268:mandelate racemase/muconate lactonizing enzyme | PF02746.18 (Mandelate racemase / muconate lactonizing enzyme, N-terminal domain)<br>PF13378.8 (Enolase C-terminal domain-like) | biosynthetic-additional |
| **10** | | PF12804.9 (MobA-like NTP transferase domain) | |
| **11** | | PF04055.23 (Radical SAM superfamily)<br>PF04055.23: GO:0003824: catalytic activity<br>PF04055.23: GO:0051536: iron-sulfur cluster binding | biosynthetic-additional |
| **12** | | PF00215.26 (Orotidine 5'-phosphate decarboxylase / HUMPS family)<br>PF00215.26: GO:0004590: orotidine-5'-phosphate decarboxylase activity<br>PF00215.26: GO:0006207: 'de novo' pyrimidine nucleobase biosynthetic process | |
| **13** | | PF01380.24 (SIS (Sugar Isomerase) domain)<br>PF01380.24: GO:0097367: carbohydrate derivative binding<br>PF01380.24: GO:1901135: carbohydrate derivative metabolic process | |
| **14** | | PF01850.23 (PIN domain) | |
| **15** | | PF04014.20 (Antidote-toxin recognition MazE, bacterial antitoxin)<br>PF04014.20: GO:0003677: DNA binding | |
| **16** | SMCOG1030:serine/threonine protein kinase | PF00069.27 (Protein kinase domain)<br>PF08308.13 (PEGA domain)<br>PF00069.27: GO:0004672: protein kinase activity<br>PF00069.27: GO:0005524: ATP binding<br>PF00069.27: GO:0006468: protein phosphorylation | regulatory |

| | | | |
|---|---|---|---|
| **17** | SMCOG1255:cold-shock DNA-binding domain protein | PF00313.24 ('Cold-shock' DNA-binding domain)<br>PF00313.24: GO:0003676: nucleic acid binding | regulatory |
| **18** | | PF13660.8 (Domain of unknown function (DUF4147))<br>PF05161.15 (MOFRL family) | |
| **19** | | PF05235.16 (CHAD domain, uncharacterized, metal binding?) | |
| **20** | | PF04055.23 (Radical SAM superfamily)<br>PF06969.18 (HemN C-terminal domain)<br>PF04055.23: GO:0003824: catalytic activity<br>PF04055.23: GO:0051536: iron-sulfur cluster binding | biosynthetic-additional |
| **21** | SMCOG1119:halogenase | PF04820.16 (Tryptophan halogenase) | biosynthetic<br>MiBIG: BGC0001897.1, 0379, Burkholderia ambifaria, Polyketide, cepacin A |
| **22** | SMCOG1002:AMP-dependent synthetase and ligase | PF00501.30 (AMP-binding enzyme)<br>PF14535.8 (AMP-binding enzyme C-terminal domain) | biosynthetic |

Table S4.16. antiSMASH annotations of contig Pf8_c00244, detected in GCF 2270 and CC G

| **Gene** | **SMCOG** | **PFAM & GO** | **Notes** |
|---|---|---|---|
| **1** | | PF12796.9 (Ankyrin repeats (3 copies)) | |
| **2** | SMCOG1139:aminotransferase class V | PF00266.21 (Aminotransferase class-V) | biosynthetic-additional |
| **3** | | PF13444.8 (Acetyltransferase (GNAT) domain) | |
| **4** | | PF14236.8 (Domain of unknown function (DUF4338))<br>PF14706.8 (Transposase DNA-binding) | |
| **5** | | PF00872.20 (Transposase, Mutator family) | |

| | | | |
|---|---|---|---|
| | | PF00872.20: GO:0003677: DNA binding<br>PF00872.20: GO:0004803: transposase activity<br>PF00872.20: GO:0006313: transposition, DNA-mediated | |
| **6** | | - | |
| **7** | | PF13540.8 (Regulator of chromosome condensation (RCC1) repeat) | |
| **8** | | - | |
| **9** | | PF13374.8 (Tetratricopeptide repeat) | |
| **10** | | - | |
| **11** | | - | |
| **12** | | PF04290.14 (Tripartite ATP-independent periplasmic transporters, DctQ component)<br>TIGR00786 (dctM: TRAP transporter, DctM subunit) | |
| **13** | | PF03480.15 (Bacterial extracellular solute-binding protein, family 7)<br>PF03480.15: GO:0055085: transmembrane transport | |
| **14** | | - | |
| **15** | | - | |
| **16** | | - | |
| **17** | | PF13570.8 (PQQ-like domain) | |
| **18** | | PF04055.23 (Radical SAM superfamily)<br>PF04055.23: GO:0003824: catalytic activity<br>PF04055.23: GO:0051536: iron-sulfur cluster binding | biosynthetic-additional |
| **19** | | - | |

| | | | |
|---|---|---|---|
| **20** | SMCOG1119:halogenase | PF04820.16 (Tryptophan halogenase) | Biosynthetic<br>MiBIG: BGC0000840.1, AAG38844.2, Xanthomonas oryzae, other, xanthomonadin I, (bmp2) |
| **21** | SMCOG1045:glycosyl transferase group 1 | PF13439.8 (Glycosyltransferase Family 4)<br>PF00534.22 (Glycosyl transferases group 1)<br>PF00534.22: GO:0016757: transferase activity, transferring glycosyl groups | biosynthetic-additional<br>MiBIG: BGC0000720.1, tbmD, Streptoalloteichus tenebrarius, Saccharide, tobramycin |
| **22** | | PF03061.24 (Thioesterase superfamily) | |
| **23** | | - | |
| **24** | SMCOG1002:AMP-dependent synthetase and ligase | PF00501.30 (AMP-binding enzyme)<br>PF14535.8 (AMP-binding enzyme C-terminal domain) | Biosynthetic<br>MiBIG:BGC0000466.1, YtkN, Streptomyces sp, NRP, yatakemycin |
| **25** | | - | |
| **26** | | - | |
| **27** | SMCOG1139:aminotransferase class V | PF00266.21 (Aminotransferase class-V) | biosynthetic-additional |
| **28** | | - | |
| **29** | | - | |
| **30** | | PF19866.1 (Family of unknown function (DUF6339)) | |
| **31** | | PF03235.16 (Protein of unknown function DUF262) | |
| **32** | | PF00872.20 (Transposase, Mutator family)<br>PF00872.20: GO:0003677: DNA binding | |

| | | |
|---|---|---|
| | | PF00872.20: GO:0004803: transposase activity<br>PF00872.20: GO:0006313: transposition, DNA-mediated |

Table S4.17. antiSMASH annotations of contig DOM14B_c00569, detected in GCF 2526 and CC O

| Gene | SMCOG | PFAM & GO | Notes |
|---|---|---|---|
| **2** | SMCOG1113:inner-membrane translocator | PF02653.18 (Branched-chain amino acid transport system / permease component)<br>PF02653.18: GO:0022857: transmembrane transporter activity<br>PF02653.18: GO:0055085: transmembrane transport<br>PF02653.18: GO:0016021: integral component of membrane | transport |
| **3** | SMCOG1000:ABC transporter ATP-binding protein | PF00005.29 (ABC transporter)<br>PF00005.29: GO:0005524: ATP binding | Transport<br>MiBIG: BGC0000643.1, ABC50111.1, Brevundimonas vesicularis, terpene, carotenoid |
| **4** | SMCOG1066:alpha/beta hydrolase domain-containing protein | PF07859.15 (alpha/beta hydrolase fold)<br>PF07859.15: GO:0016787: hydrolase activity | biosynthetic-additional<br>MiBIG: BGC0000820.1, psmB, Streptomyces griseofuscus, alkaloid, physostigmine |
| **5** | | PF01717.20 (Cobalamin-independent synthase, Catalytic domain)<br>PF01717.20: GO:0003871: 5-methyltetrahydropteroyltriglutamate-homocysteine S-methyltransferase activity<br>PF01717.20: GO:0008270: zinc ion binding<br>PF01717.20: GO:0009086: methionine biosynthetic process | |
| **6** | | PF01558.20 (Pyruvate ferredoxin/flavodoxin oxidoreductase)<br>PF01558.20: GO:0016903: oxidoreductase activity, acting on the aldehyde or oxo group of donors<br>PF01558.20: GO:0055114: oxidation-reduction process | |

| | | | |
|---|---|---|---|
| **7** | | PF02775.23 (Thiamine pyrophosphate enzyme, C-terminal TPP binding domain)<br>PF02775.23: GO:0003824: catalytic activity<br>PF02775.23: GO:0030976: thiamine pyrophosphate binding | |
| **8** | | PF01381.24 (Helix-turn-helix)<br>TIGR02607 (antidote_HigA: addiction module antidote protein, HigA family) | |
| **9** | | PF05015.15 (RelE-like toxin of type II toxin-antitoxin system HigB) | |
| **10** | | PF13380.8 (CoA binding domain)<br>PF13607.8 (Succinyl-CoA ligase like flavodoxin domain)<br>PF13549.8 (ATP-grasp domain) | |
| **11** | | PF06803.14 (Protein of unknown function (DUF1232)) | |
| **12** | | PF02538.16 (Hydantoinase B/oxoprolinase)<br>PF02538.16: GO:0003824: catalytic activity | |
| **13** | | PF03401.16 (Tripartite tricarboxylate transporter family receptor) | |
| **14** | | PF13673.9 (Acetyltransferase (GNAT) domain)<br>PF13673.9: GO:0008080: N-acetyltransferase activity | |
| **15** | | PF03358.17 (NADPH-dependent FMN reductase)<br>PF03358.17: GO:0016491: oxidoreductase activity | |
| **16** | SMCOG1293:aldehyde oxidase and xanthine dehydrogenase | PF01315.24 (Aldehyde oxidase and xanthine dehydrogenase, a/b hammerhead domain)<br>PF02738.20 (Molybdopterin-binding domain of aldehyde dehydrogenase)<br>PF02738.20: GO:0016491: oxidoreductase activity<br>PF02738.20: GO:0055114: oxidation-reduction process | biosynthetic-additional |
| **17** | | PF00941.23 (FAD binding domain in molybdopterin dehydrogenase)<br>PF03450.19 (CO dehydrogenase flavoprotein C-terminal domain) | |

| | | | |
|---|---|---|---|
| | | PF00941.23: GO:0016491: oxidoreductase activity<br>PF00941.23: GO:0055114: oxidation-reduction process | |
| **18** | | PF00111.29 (2Fe-2S iron-sulfur cluster binding domain)<br>PF01799.22 ([2Fe-2S] binding domain)<br>PF00111.29: GO:0009055: electron transfer activity<br>PF00111.29: GO:0051536: iron-sulfur cluster binding<br>PF01799.22: GO:0016491: oxidoreductase activity<br>PF01799.22: GO:0046872: metal ion binding<br>PF01799.22: GO:0055114: oxidation-reduction process | MiBIG: BGC0001973.1, Shi4225, Streptomyces hiroshimensis, other, fervenulin |
| **19** | SMCOG1002:AMP-dependent synthetase and ligase | PF00501.30 (AMP-binding enzyme)<br>PF13193.8 (AMP-binding enzyme C-terminal domain) | Biosynthetic<br>MiBIG: BGC0000649.1, fadD, Streptomyces griseus, Terpene, carotenoid |
| **20** | SMCOG1023:enoyl-CoA hydratase | PF00378.22 (Enoyl-CoA hydratase/isomerase)<br>PF00378.22: GO:0003824: catalytic activity | biosynthetic-additional |
| **21** | SMCOG1020:major facilitator transporter | PF05977.15 (Transmembrane secretion effector) | transport |
| **22** | SMCOG1050:monooxygenase FAD-binding | PF01494.21 (FAD binding domain)<br>PF01494.21: GO:0071949: FAD binding | biosynthetic |
| **23** | | PF02776.20 (Thiamine pyrophosphate enzyme, N-terminal TPP binding domain)<br>PF02776.20: GO:0030976: thiamine pyrophosphate binding | |
| **24** | | PF02775.23 (Thiamine pyrophosphate enzyme, C-terminal TPP binding domain)<br>PF02775.23: GO:0003824: catalytic activity<br>PF02775.23: GO:0030976: thiamine pyrophosphate binding | |
| **25** | | - | |

| | | |
|---|---|---|
| **26** | | PF01966.24 (HD domain, metal dependent phosphohydrolase) |

Table S4.18. antiSMASH annotations of contig gb4_2_c02126, detected in GCF 2557 and CC M

| Gene | SMCOG | PFAM & GO | Notes |
|---|---|---|---|
| **1** | | PF00939.21 (Sodium:sulfate symporter transmembrane region)<br>TIGR00785 (dass: transporter, divalent anion:Na+ symporter (DASS) family<br>PF00939.21: GO:0022857: transmembrane transporter activity<br>PF00939.21: GO:0055085: transmembrane transport<br>PF00939.21: GO:0016020: membrane | |
| **2** | SMCOG1175:pyridine nucleotide-disulfide oxidoreductase | PF07992.16 (Pyridine nucleotide-disulphide oxidoreductase)<br>PF02852.24 (Pyridine nucleotide-disulphide oxidoreductase, dimerisation domain)<br>TIGR01350 (lipoamide_DH: dihydrolipoyl dehydrogenase) | biosynthetic |
| **3** | SMCOG1002:AMP-dependent synthetase and ligase | PF00501.30 (AMP-binding enzyme) | biosynthetic |
| **4** | | - | |
| **5** | | - | |
| **6** | SMCOG1001:short-chain dehydrogenase/reductase SDR | PF13561.8 (Enoyl-(Acyl carrier protein) reductase)<br>2,3-dihydroxybenzoate-2,3-dehydrogenase | biosynthetic-additional<br>MiBIG: BGC0001538.1, 23870, Streptomyces sp., NRP, Polyketide, caboxamycin |
| **7** | | PF05721.15 (Phytanoyl-CoA dioxygenase (PhyH)) | |
| **8** | | PF00753.29 (Metallo-beta-lactamase superfamily)<br>PF07521.14 (Zn-dependent metallo-hydrolase RNA specificity domain, pre-mRNA 3'-end-processing endonucleases )<br>PF17770.3 (Ribonuclease J C-terminal domain)<br>TIGR00649 (MG423: beta-CASP ribonuclease, RNase J family) | |

| | | | |
|---|---|---|---|
| **9** | | Peptidase_S9 | biosynthetic-additional |
| **10** | SMCOG1184:major facilitator transporter | PF07690.18 (Major Facilitator Superfamily)<br>PF07690.18: GO:0022857: transmembrane transporter activity<br>PF07690.18: GO:0055085: transmembrane transport | transport |

Table S4.19. antiSMASH annotations of contig gb3_2_c07789, detected in GCF 2484 and CC T

| Gene | SMCOG | PFAM & GO | Notes |
|---|---|---|---|
| **1** | | PF08734.13 (GYD domain, unknown function) | |
| **2** | SMCOG1002:AMP-dependent synthetase and ligase | PF16177.7 (Acetyl-coenzyme A synthetase N-terminus)<br>PF00501.30 (AMP-binding enzyme)<br>PF13193.8 (AMP-binding enzyme C-terminal domain)<br>TIGR02188 (Ac_CoA_lig_AcsA: acetate--CoA ligase) | biosynthetic |
| **3** | SMCOG1161:oxidoreductase | PF05834.14 (Lycopene cyclase protein)<br>TIGR02032 (GG-red-SF: geranylgeranyl reductase family) | Biosynthetic<br>MiBIG: BGC0000282.1, SGR_470 (monooxygenase), Streptomyces griseus, polyketide, alkylresorcinol |
| **4** | SMCOG1146:sugar transferase | PF02397.18 (Bacterial sugar transferase)<br>UDP-glucose dehydrogenase | biosynthetic-additional<br>MiBIG: BGC0000868.1, tuaA, Bacillus subtilis, other, teichuronic acid |
| **5** | | PF02673.20 (Bacitracin resistance protein BacA)<br>PF02673.20: GO:0050380: undecaprenyl-diphosphatase activity<br>PF02673.20: GO:0016311: dephosphorylation<br>PF02673.20: GO:0016020: membrane | |
| **6** | SMCOG1226:hypothetical protein | PF03781.18 (Sulfatase-modifying factor enzyme 1, domain is also found in iron(II)-dependent oxidoreductase ) | |

| | | | |
|---|---|---|---|
| **7** | SMCOG1268:mandelate racemase/muconate lactonizing enzyme | PF02746.18 (Mandelate racemase / muconate lactonizing enzyme, N-terminal domain, also found in dehydratases )<br>PF13378.8 (Enolase C-terminal domain-like) | biosynthetic-additional |

Table S4.20. antiSMASH annotations of contig DOM057_c01231, detected in GCF 2010 and CC L

| Gene | SMCOG | PFAM & GO | Notes |
|---|---|---|---|
| **1** | | - | |
| **2** | SMCOG1045:glycosyl transferase group 1 | PF13439.8 (Glycosyltransferase Family 4)<br>PF00534.22 (Glycosyl transferases group 1)<br>PF00534.22: GO:0016757: transferase activity, transferring glycosyl groups | biosynthetic-additional |
| **3** | | PF04069.14 (Substrate binding domain of ABC-type glycine betaine transport system)<br>PF04069.14: GO:0022857: transmembrane transporter activity<br>PF04069.14: GO:0055085: transmembrane transport<br>PF04069.14: GO:0043190: ATP-binding cassette (ABC) transporter complex | |
| **4** | | PF17645.3 (Arylmalonate decarboxylase) | |
| **5** | | PF01029.20 (NusB family)<br>PF01189.19 (16S rRNA methyltransferase RsmB/F)<br>TIGR00563 (rsmB: 16S rRNA (cytosine(967)-C(5))-methyltransferase)<br>PF01029.20: GO:0003723: RNA binding<br>PF01029.20: GO:0006355: regulation of transcription, DNA-templated<br>PF01189.19: GO:0008168: methyltransferase activity | MiBIG: BGC0002034.1, 28395, Streptomyces sp., other, perquinolines, rRNA cytosine-C5-methyltransferase |
| **6** | SMCOG1228:methionyl-tRNA formyltransferase | PF00551.21 (Formyl transferase)<br>PF02911.20 (Formyl transferase, C-terminal domain)<br>TIGR00460 (fmt: methionyl-tRNA formyltransferase)<br>PF00551.21: GO:0016742: hydroxymethyl-, formyl- and related transferase activity<br>PF00551.21: GO:0009058: biosynthetic process | biosynthetic-additional |

| | | | |
|---|---|---|---|
| | | PF02911.20: GO:0016742: hydroxymethyl-, formyl- and related transferase activity<br>PF02911.20: GO:0009058: biosynthetic process | |
| **7** | SMCOG1275:peptide deformylase | PF01327.23 (Polypeptide deformylase)<br>TIGR00079 (pept_deformyl: peptide deformylase)<br>PF01327.23: GO:0042586: peptide deformylase activity | biosynthetic-additional |
| **8** | | PF17764.3 (3'DNA-binding domain (3'BD))<br>PF04851.17 (Type III restriction enzyme, res subunit)<br>PF18319.3 (PriA DNA helicase Cys-rich region (CRR) domain)<br>PF00271.33 (Helicase conserved C-terminal domain)<br>PF18074.3 (Primosomal protein N C-terminal domain)<br>TIGR00595 (priA: primosomal protein N’)<br>PF04851.17: GO:0003677: DNA binding<br>PF04851.17: GO:0005524: ATP binding<br>PF04851.17: GO:0016787: hydrolase activity<br>PF17764.3: GO:0003677: DNA binding | |
| **9** | | PF11104.10 (Type IV pilus assembly protein PilM;)<br>TIGR01175 (pilM: type IV pilus assembly protein PilM) | |
| **10** | | PF05137.15 (Fimbrial assembly protein (PilN))<br>PF04350.15 (Pilus assembly protein, PilO)<br>PF04351.15 (Pilus assembly protein, PilP)<br>PF04350.15: GO:0043107: type IV pilus-dependent motility<br>PF04350.15: GO:0043683: type IV pilus biogenesis | |
| **11** | | - | |
| **12** | SMCOG1001:short-chain dehydrogenase/reductase SDR | PF00106.27 (short chain dehydrogenase)<br>PF00106.27: GO:0055114: oxidation-reduction process | biosynthetic-additional |
| **13** | | PF07075.13 (Protein of unknown function (DUF1343)) | |

| | | | |
|---|---|---|---|
| **14** | SMCOG1166:transporter, EamA family | PF00892.22 (EamA-like transporter family)<br>PF00892.22: GO:0016020: membrane<br>PF00892.22: GO:0016021: integral component of membrane | transport |
| **15** | | PF12838.9 (4Fe-4S dicluster domain, in bacterial ferredoxins, various dehydrogenases, and various reductases) | |
| **16** | | PF00890.26 (FAD binding domain)<br>PF02910.22 (Fumarate reductase flavoprotein C-term)<br>PF02910.22: GO:0016491: oxidoreductase activity<br>PF02910.22: GO:0055114: oxidation-reduction process | biosynthetic |
| **17** | SMCOG1002:AMP-dependent synthetase and ligase | PF16177.7 (Acetyl-coenzyme A synthetase N-terminus)<br>PF00501.30 (AMP-binding enzyme)<br>PF13193.8 (AMP-binding enzyme C-terminal domain) | Biosynthetic<br>MiBIG: BGC0001538.1, 23850, Streptomyces sp., NRP, Polyketide, caboxamycin, Triostin synthetase I |
| **18** | | PF01977.18 (3-octaprenyl-4-hydroxybenzoate carboxy-lyase)<br>PF01977.18: GO:0016831: carboxy-lyase activity | MiBIG: BGC0000889.1, fevL, Streptomyces sp., other, bagremycin, decarboxylase |
| **19** | | PF00903.27 (Glyoxalase/Bleomycin resistance protein/Dioxygenase superfamily) | |
| **20** | | - | |
| **21** | | PF04909.16 (Amidohydrolase)<br>PF04909.16: GO:0016787: hydrolase activity | |
| **22** | | PF04909.16 (Amidohydrolase)<br>PF04909.16: GO:0016787: hydrolase activity | |
| **23** | | DXP_synthase_N (PF13292 - transketolase)<br>PF00676.22 (Dehydrogenase E1 component) | biosynthetic-additional |

| | | | |
|---|---|---|---|
| | | PF00676.22: GO:0016624: oxidoreductase activity, acting on the aldehyde or oxo group of donors, disulfide as acceptor | |
| **24** | SMCOG1110:1-deoxy-D-xylulose-5-phosphate synthase | PF02779.26 (Transketolase, pyrimidine binding domain)<br>PF02780.22 (Transketolase, C-terminal domain) | biosynthetic-additional |
| **25** | | PF02633.16 (Creatinine amidohydrolase) | |
| **26** | | - | |
| **27** | | - | |
| **28** | | PF12441.10 (CopG antitoxin of type II toxin-antitoxin system) | |
| **29** | | - | |
| **30** | SMCOG1040:alcohol dehydrogenase | PF08240.14 (Alcohol dehydrogenase GroES-like domain)<br>PF00107.28 (Zinc-binding dehydrogenase)<br>PF00107.28: GO:0055114: oxidation-reduction process<br>PF08240.14: GO:0055114: oxidation-reduction process | biosynthetic-additional |
| **31** | | PF07883.13 (Cupin domain) | |
| **32** | | PF17645.3 (Arylmalonate decarboxylase) | |
| **33** | | PF03992.18 (Antibiotic biosynthesis monooxygenase) | |
| **34** | SMCOG1189:L-carnitine dehydratase/bile acid-inducible | PF02515.19 (CoA-transferase family III)<br>PF02515.19: GO:0008410: CoA-transferase activity | biosynthetic-additional |
| **35** | | PF05145.14 (Transition state regulatory protein AbrB)<br>PF05145.14: GO:0010468: regulation of gene expression<br>PF05145.14: GO:0016021: integral component of membrane | |
| **36** | | PF00561 (alpha beta hydrolase fold) | biosynthetic-additional |

| | | | |
|---|---|---|---|
| | | Peptidase_S9<br>PF02129.20 (X-Pro dipeptidyl-peptidase (S15 family))<br>PF02129.20: GO:0016787: hydrolase activity | |
| **37** | | - | |
| **38** | SMCOG1293:aldehyde oxidase and xanthine dehydrogenase | PF01315.24 (Aldehyde oxidase and xanthine dehydrogenase, a/b hammerhead domain)<br>PF02738.20 (Molybdopterin-binding domain of aldehyde dehydrogenase)<br>PF02738.20: GO:0016491: oxidoreductase activity<br>PF02738.20: GO:0055114: oxidation-reduction process | biosynthetic-additional |
| **39** | | - | |

Table S4.21. antiSMASH annotations of contig gb4_2_c00127, detected in GCF 2522 and CC N

| Gene | SMCOG | PFAM & GO | Notes |
|---|---|---|---|
| **1** | | - | |
| **2** | | PF02604.21 (Antitoxin Phd_YefM, type II toxin-antitoxin system)<br>TIGR01552 (phd_fam: prevent-host-death family protein) | |
| **3** | | PF01850.23 (PIN domain) | |
| **4** | | PF01145.27 (SPFH domain / Band 7 family – membrane associated)<br>PF15975.7 (Flotillin) | |
| **5** | | - | |
| **6** | | - | |
| **7** | | PF06808.14 (Tripartite ATP-independent periplasmic transporter, DctM component)<br>TIGR02123 (TRAP_fused: TRAP transporter, 4TM/12TM fusion protein) | |
| **8** | | PF16868.7 (NMT1-like family) | |

| | | | |
|---|---|---|---|
| | | TIGR02122 (TRAP_TAXI: TRAP transporter solute receptor, TAXI family) | |
| **9** | | PF01661.23 (Macro domain, chromatin biology, DNA repair and transcription regulation) | MiBIG: BGC0000249.1, 42849.1, Streptomyces nogalater, PKS, nogalamycin |
| **10** | | PF00903.27 (Glyoxalase/Bleomycin resistance protein/Dioxygenase superfamily) | |
| **11** | | PF00501.30 (AMP-binding enzyme)<br>PF14535.8 (AMP-binding enzyme C-terminal domain) | Biosynthetic<br>MiBIG: BGC0000889.1, fevW, Streptomyces sp., other, bagremycin |
| **12** | | PF03061.24 (Thioesterase superfamily) | |
| **13** | SMCOG1119:halogenase | PF04820.16 (Tryptophan halogenase) | Biosynthetic<br>MiBIG: BGC0001366.1, orf12, Streptomyces sp, PKS, zunyimycin, Trp_halogenase |
| **14** | | - | |
| **15** | | PF04055.23 (Radical SAM superfamily)<br>PF04055.23: GO:0003824: catalytic activity<br>PF04055.23: GO:0051536: iron-sulfur cluster binding | biosynthetic-additional |
| **16** | | PF13570.8 (PQQ-like domain) | |
| **17** | | PF00465.21 (Iron-containing alcohol dehydrogenase)<br>PF00465.21: GO:0016491: oxidoreductase activity<br>PF00465.21: GO:0046872: metal ion binding<br>PF00465.21: GO:0055114: oxidation-reduction proces | |
| **18** | | PF02518.28 (Histidine kinase-, DNA gyrase B-, and HSP90-like ATPase)<br>PF00204.27 (DNA gyrase B) | MiBIG: |

| | | | |
|---|---|---|---|
| | | PF01751.24 (Toprim domain)<br>PF00986.23 (DNA gyrase B subunit, carboxyl terminus)<br>PF00204.27: GO:0003677: DNA binding<br>PF00204.27: GO:0003918: DNA topoisomerase type II (double strand cut, ATP-hydrolyzing) activity<br>PF00204.27: GO:0005524: ATP binding<br>PF00204.27: GO:0006265: DNA topological change<br>PF00986.23: GO:0003677: DNA binding<br>PF00986.23: GO:0003918: DNA topoisomerase type II (double strand cut, ATP-hydrolyzing) activity<br>PF00986.23: GO:0005524: ATP binding<br>PF00986.23: GO:0006265: DNA topological change | |
| **19** | | PF00521.22 (DNA gyrase/topoisomerase IV, subunit A<br>PF00521.22: GO:0003677: DNA binding<br>PF00521.22: GO:0003918: DNA topoisomerase type II (double strand cut, ATP-hydrolyzing) activity<br>PF00521.22: GO:0005524: ATP binding<br>PF00521.22: GO:0006265: DNA topological change | |
| **20** | | PF00293.30 (NUDIX domain, pyrophosphohydrolases)<br>PF00293.30: GO:0016787: hydrolase activity | |
| **21** | SMCOG1086:MATE efflux family protein | PF01554.20 (MatE)<br>TIGR00797 (matE: MATE efflux family protein)<br>PF01554.20: GO:0015297: antiporter activity<br>PF01554.20: GO:0042910: xenobiotic transmembrane transporter activity<br>PF01554.20: GO:0055085: transmembrane transport<br>PF01554.20: GO:0016020: membrane | transport |
| **22** | SMCOG1168:O-succinylhomoserine sulfhydrylase | PF01053.22 (Cys/Met metabolism PLP-dependent enzyme)<br>PF01053.22: GO:0030170: pyridoxal phosphate binding<br>PF01053.22: GO:0019346: transsulfuration | biosynthetic-additional |

| | | | |
|---|---|---|---|
| **23** | | PF00583.27 (Acetyltransferase (GNAT) family)<br>PF00583.27: GO:0008080: N-acetyltransferase activity | |
| **24** | SMCOG1040:alcohol dehydrogenase | PF08240.14 (Alcohol dehydrogenase GroES-like domain)<br>PF00107.28 (Zinc-binding dehydrogenase)<br>PF00107.28: GO:0055114: oxidation-reduction process<br>PF08240.14: GO:0055114: oxidation-reduction process | biosynthetic-additional |
| **25** | | - | |
| **26** | SMCOG1002:AMP-dependent synthetase and ligase | PF00550.27 (Phosphopantetheine attachment site)<br>PF00501.30 (AMP-binding enzyme)<br>PF01553.23 (Acyltransferase)<br>PF01553.23: GO:0016746: transferase activity, transferring acyl groups | biosynthetic |
| **27** | | PF00160.23 (Cyclophilin type peptidyl-prolyl cis-trans isomerase/CLD)<br>PF00160.23: GO:0003755: peptidyl-prolyl cis-trans isomerase activity<br>PF00160.23: GO:0000413: protein peptidyl-prolyl isomerization | |
| **28** | | PF00160.23 (Cyclophilin type peptidyl-prolyl cis-trans isomerase/CLD)<br>PF00160.23: GO:0003755: peptidyl-prolyl cis-trans isomerase activity<br>PF00160.23: GO:0000413: protein peptidyl-prolyl isomerization | |
| **29** | | - | |

Table S4.22. antiSMASH annotations of contig gb9_c09452, detected in GCF 2609 and CC Q

| Gene | SMCOG | PFAM & GO | Notes |
|---|---|---|---|
| **1** | | PF12697.9 (Alpha/beta hydrolase family) | biosynthetic-additional |
| **2** | SMCOG1002:AMP-dependent synthetase and ligase | PF00501.30 (AMP-binding enzyme)<br>PF13193.8 (AMP-binding enzyme C-terminal domain) | Biosynthetic<br>MiBIG: BGC0000649.1, fadD, Streptomyces griseus, terpene, carotenoid |

| | | | |
|---|---|---|---|
| **3** | SMCOG1119:halogenase | PF04820.16 (Tryptophan halogenase) | Biosynthetic<br>MiBIG: BGC0000840.1, 38844.2, Xanthomonas oryzae, other, xanthomonadin |
| **4** | | -<br>(blast – puttative ACP) | |
| **5** | SMCOG1001:short-chain dehydrogenase/reductase SDR | PF13561.8 (Enoyl-(Acyl carrier protein) reductase) | biosynthetic-additional<br>MiBIG: BGC0001987.1, 02930, Pseudomonas koreensis, PKS, koreenceines, 3-oxoacyl-ACP reductase |
| **6** | | PF00916.22 (Sulfate permease family)<br>PF00916.22: GO:0015116: sulfate transmembrane transporter activity<br>PF00916.22: GO:0008272: sulfate transport<br>PF00916.22: GO:0016021: integral component of membrane | |

Table S4.23. antiSMASH annotations of contig Pf7_c00139, detected in GCF 2366 and CC S

| **Gene** | **SMCOG** | **PFAM & GO** | **Notes** |
|---|---|---|---|
| **1** | | - | |
| **2** | SMCOG1207:inositol monophosphatase | PF00459.27 (Inositol monophosphatase family)<br>PF00459.27: GO:0046855: inositol phosphate dephosphorylation | biosynthetic-additional |
| **3** | | PF01222.19 (Ergosterol biosynthesis ERG4/ERG24 family)<br>PF01222.19: GO:0016628: oxidoreductase activity, acting on the CH-CH group of donors, NAD or NADP as acceptor<br>PF01222.19: GO:0016126: sterol biosynthetic process<br>PF01222.19: GO:0016020: membrane | |
| **4** | | PF01222.19 (Ergosterol biosynthesis ERG4/ERG24 family) | |

| | | | |
|---|---|---|---|
| | | PF01222.19: GO:0016628: oxidoreductase activity, acting on the CH-CH group of donors, NAD or NADP as acceptor<br>PF01222.19: GO:0016126: sterol biosynthetic process<br>PF01222.19: GO:0016020: membrane | |
| **5** | | - | |
| **6** | | - | |
| **7** | | PF03061.24 (Thioesterase superfamily) | |
| **8** | | PF08768.13 (THAP4-like, heme-binding beta-barrel domain), putatively related to fatty acid-binding proteins | |
| **9** | SMCOG1092:hypothetical protein | PF13450.8 (NAD(P)-binding Rossmann-like domain) | |
| **10** | | PF01212.23 (Beta-eliminating lyase)<br>PF01212.23: GO:0016829: lyase activity<br>PF01212.23: GO:0006520: cellular amino acid metabolic process<br>(PF01041: pyridoxal-phosphate-dependent aminotransferase enzymes with a variety of molecular functions) | biosynthetic-additional<br>DegT_DnrJ_EryC1 |
| **11** | | PF07969.13 (Amidohydrolase family) | |
| **12** | | - | |
| **13** | SMCOG1160:flavin reductase domain protein FMN-binding | PF01613.20 (Flavin reductase like domain)<br>PF01613.20: GO:0010181: FMN binding | biosynthetic-additional |
| **14** | | PF01176.21 (Translation initiation factor 1A / IF-1)<br>TIGR00008 (infA: translation initiation factor IF-1)<br>PF01176.21: GO:0003723: RNA binding<br>PF01176.21: GO:0003743: translation initiation factor activity<br>PF01176.21: GO:0006413: translational initiation | |

| | | | |
|---|---|---|---|
| **15** | SMCOG1023:enoyl-CoA hydratase | PF00378.22 (Enoyl-CoA hydratase/isomerase)<br>PF00378.22: GO:0003824: catalytic activity | biosynthetic-additional |
| **16** | SMCOG1006:acyl-CoA dehydrogenase | PF02771.18 (Acyl-CoA dehydrogenase, N-terminal domain)<br>PF02770.21 (Acyl-CoA dehydrogenase, middle domain)<br>PF00441.26 (Acyl-CoA dehydrogenase, C-terminal domain)<br>PF00441.26: GO:0016627: oxidoreductase activity, acting on the CH-CH group of donors<br>PF00441.26: GO:0055114: oxidation-reduction process<br>PF02770.21: GO:0016627: oxidoreductase activity, acting on the CH-CH group of donors<br>PF02770.21: GO:0055114: oxidation-reduction process<br>PF02771.18: GO:0016627: oxidoreductase activity, acting on the CH-CH group of donors<br>PF02771.18: GO:0050660: flavin adenine dinucleotide binding<br>PF02771.18: GO:0055114: oxidation-reduction process | biosynthetic-additional |
| **17** | | PF13416.8 (Bacterial extracellular solute-binding protein): | |
| **18** | SMCOG1069:putative ABC transporter permease protein | PF00528.24 (Binding-protein-dependent transport system inner membrane component)<br>PF00528.24: GO:0055085: transmembrane transport<br>PF00528.24: GO:0016020: membrane | transport |
| **19** | SMCOG1069:putative ABC transporter permease protein | PF00528.24 (Binding-protein-dependent transport system inner membrane component)<br>PF00528.24: GO:0055085: transmembrane transport<br>PF00528.24: GO:0016020: membrane | transport |
| **20** | SMCOG1000:ABC transporter ATP-binding protein | PF00005.29 (ABC transporter)<br>PF08402.12 (TOBE domain)<br>TIGR01187 (potA: polyamine ABC transporter, ATP-binding protein)<br>PF00005.29: GO:0005524: ATP binding<br>PF08402.12: GO:0005524: ATP binding | transport |

| | | | |
|---|---|---|---|
| | | PF08402.12: GO:0022857: transmembrane transporter activity<br>PF08402.12: GO:0055085: transmembrane transport<br>PF08402.12: GO:0043190: ATP-binding cassette (ABC) transporter complex | |
| **21** | | PF00491.23 (Arginase family)<br>TIGR01230 (agmatinase: agmatinase)<br>PF00491.23: GO:0046872: metal ion binding | |
| **22** | | - | |
| **23** | SMCOG1103:FAD dependent oxidoreductase | PF01266.26 (FAD dependent oxidoreductase)<br>PF01266.26: GO:0016491: oxidoreductase activity<br>PF01266.26: GO:0055114: oxidation-reduction process | biosynthetic |
| **24** | | PF00753.29 (Metallo-beta-lactamase superfamily) | |
| **25** | | PF11716.10 (Mycothiol maleylpyruvate isomerase metal-binding N-terminal domain)<br>TIGR03086 (TIGR03086: TIGR03086 family protein)<br>TIGR03083 (TIGR03083: uncharacterized Actinobacterial protein TIGR03083)<br>PF11716.10: GO:0046872: metal ion binding | |
| **26** | | - | |
| **27** | | PF03631.17 (Virulence factor BrkB) | |
| **28** | SMCOG1002:AMP-dependent synthetase and ligase | PF00501.30 (AMP-binding enzyme)<br>PF13193.8 (AMP-binding enzyme C-terminal domain) | Biosynthetic<br>MiBIG: BGC0001538.1, 23850, Streptomyces sp., NRP, Polyketide, caboxamycin |
| **29** | SMCOG1134:flavodoxin | PF04879.18 (Molybdopterin oxidoreductase Fe4S4 domain)<br>PF00384.24 (Molybdopterin oxidoreductase)<br>PF01568.23 (Molydopterin dinucleotide binding domain)<br>PF00384.24: GO:0016491: oxidoreductase activity<br>PF00384.24: GO:0055114: oxidation-reduction process | biosynthetic-additional<br>MiBIG: BGC0001441.1, belN, Streptomyces sp., other, belactosin |

| | | | |
|---|---|---|---|
| | | PF01568.23: GO:0016491: oxidoreductase activity<br>PF01568.23: GO:0043546: molybdopterin cofactor binding<br>PF01568.23: GO:0055114: oxidation-reduction process<br>PF04879.18: GO:0016491: oxidoreductase activity<br>PF04879.18: GO:0055114: oxidation-reduction process | |
| **30** | SMCOG1202:major facilitator transportee | PF07690.18 (Major Facilitator Superfamily)<br>PF07690.18: GO:0022857: transmembrane transporter activity<br>PF07690.18: GO:0055085: transmembrane transport | transport |
| **31** | | PF01699.26 (Sodium/calcium exchanger protein)<br>TIGR00367 (TIGR00367: K+-dependent Na+/Ca+ exchanger homolog)<br>PF01699.26: GO:0055085: transmembrane transport<br>PF01699.26: GO:0016021: integral component of membrane | |
| **32** | | PF00920.23 (Dehydratase family)<br>PF00920.23: GO:0003824: catalytic activity | |
| **33** |  | PF00676.22 (pyruvate Dehydrogenase E1 component, acetyl transferring)<br>PF00676.22: GO:0016624: oxidoreductase activity, acting on the aldehyde or oxo group of donors, disulfide as acceptor | MiBIG: BGC0001535.1, bkdA1, Streptomyces filamentosus, other, daptomycin analogs |
| **34** | SMCOG1110:1-deoxy-D-xylulose-5-phosphate synthase | PF02779.26 (Transketolase, pyrimidine binding domain<br>PF02780.22 (Transketolase, C-terminal domain) | biosynthetic-additional<br>MiBIG: BGC0001535.1, bkdB1, Streptomyces filamentosus, other, daptomycin analogs |
| **35** | | PF01513.23 (ATP-NAD kinase N-terminal domain)<br>PF01513.23: GO:0003951: NAD+ kinase activity<br>PF01513.23: GO:0006741: NADP biosynthetic process | |
| **36** |  | PF00364.24 (Biotin-requiring enzyme)<br>PF02817.19 (e3 binding domain)<br>PF00198.25 (2-oxoacid dehydrogenases acyltransferase (catalytic domain))<br>PF00198.25: GO:0016746: transferase activity, transferring acyl groups | MiBIG: BGC0001535.1, bkdC2, Streptomyces filamentosus, other, daptomycin analogs |

| | | | |
|---|---|---|---|
| | | PF02817.19: GO:0016746: transferase activity, transferring acyl groups | |
| **37** | SMCOG1006:acyl-CoA dehydrogenase | PF02771.18 (Acyl-CoA dehydrogenase, N-terminal domain)<br>PF02770.21 (Acyl-CoA dehydrogenase, middle domain)<br>PF00441.26 (Acyl-CoA dehydrogenase, C-terminal domain)<br>PF00441.26: GO:0016627: oxidoreductase activity, acting on the CH-CH group of donors<br>PF00441.26: GO:0055114: oxidation-reduction process<br>PF02770.21: GO:0016627: oxidoreductase activity, acting on the CH-CH group of donors<br>PF02770.21: GO:0055114: oxidation-reduction process<br>PF02771.18: GO:0016627: oxidoreductase activity, acting on the CH-CH group of donors<br>PF02771.18: GO:0050660: flavin adenine dinucleotide binding<br>PF02771.18: GO:0055114: oxidation-reduction process | biosynthetic-additional<br>MiBIG: BGC0000886.1, allD, Streptomyces tsukubensis, other, allylmalonyl-CoA |
| **38** | | PF00903.27 (Glyoxalase/Bleomycin resistance protein/Dioxygenase superfamily) | |
| **39** | SMCOG1092:hypothetical protein | PF13450.8 (NAD(P)-binding Rossmann-like domain) | MiBIG: BGC0001625.1, ifqQ, Streptomyces sp., other, isofuranonaphthoquinone, Baeyer-Villiger monooxygenase |
| **40** | | PF12710.9 (haloacid dehalogenase-like hydrolase)<br>TIGR02137 (HSK-PSP: phosphoserine phosphatase/homoserine phosphotransferase bifunctional protein) | |
| **41** | | PF08534.12 (Redoxin)<br>PF08534.12: GO:0016491: oxidoreductase activity | |
| **42** | SMCOG1108:ROK family protein | PF00480.22 (ROK (Repressor, ORF, Kinase) family) | |

| | | |
|---|---|---|
| **43** | | PF02113.17 (D-Ala-D-Ala carboxypeptidase 3 (S13) family)<br>TIGR00666 (PBP4: D-alanyl-D-alanine carboxypeptidase/D-alanyl-D-alanine-endopeptidase)<br>PF02113.17: GO:0004185: serine-type carboxypeptidase activity<br>PF02113.17: GO:0006508: proteolysis |
| **44** | | PF02190.18 (ATP-dependent protease La (LON) substrate-binding domain) |

**Table S5: Description of regulator elements as detected by antiSMASH 7 (beta) on CC representative contigs.**

| Contig | GCF | CC | Regulator | Location | Description | Score |
|---|---|---|---|---|---|---|
| **Pf12_c00118** | 2286 | A | AfsR | 5,401 | Pleiotropic regulatory for antibiotic production | 18.46 out of 23.05 |
| **Pf12_c00118** | 2286 | A | AbrC3 | 29,809 | Antibiotic production activator | 18.07 out of 18.16 |
| **gb5_2_c00041** | 2402 | E | AbrC3 | 11,303 | Antibiotic production activator | 17.35 out of 18.16 |
| **gb5_2_c00041** | 2402 | E | CatR | 15,415 | H2O2-responsive repressor | 20.57 out of 25.31 |
| **Pf12_c00284** | 2043 | K | ZuR | 32,803 | Zinc-responsive repressor | 20.85 out of 28.19 |
| **Pf11_c00818** | 2559 | H | NrdR | 12,295 | Represses ribonucleotide reductase encoding genes | 23.69 out of 27.18 |
| **Pf11_c00818** | 2559 | H | LexA | 18,416 | Repressor of DNA damage response | 23.92 out of 29.65 |
| **DOM044_c00402** | 1983 | D | AbrC3 | 32,418 | Antibiotic production activator | 18.07 out of 18.16 |
| **DOM044_c00402** | 1983 | D | ZuR | 32,531 | Zinc-responsive repressor | 21.63 out of 28.19 |
| **DOM14B_c00569** | 2526 | O | CatR | 14,503 | H2O2-responsive repressor | 21.85 out of 25.31 |
| **gb9_c09452** | 2609 | Q | BldD | 1,031 | Development and antibiotic global regulator | 20.01 out of 23.43 |

### S7: CC A A-domain AdenylPred

**Table S6.** CC A A-domain AdenylPred functional class and substrate specificity predictions.

| A-domain | Predicted functional class (FC) | FC prediction probability | Predicted substrate specificity (SS) | SS prediction probability |
|---|---|---|---|---|
| Pf6_c44645_NODE_44...region001_haloAMP_AMP-binding_1_304_1795 | Aryl-CoA ligase | 0.41 | cinnamate and succinylbenzoate derivatives | 0.24 |
| gb9_c07451_NODE_74...region001_haloAMP_AMP-binding_5_6089_7670 | Aryl-CoA ligase | 0.42 | cinnamate and succinylbenzoate derivatives | 0.27 |

| | | | | |
|---|---|---|---|---|
| DOM049_bin.11.strict_NODE_225_length_5496_cov_4.161100.region001_haloAMP_AMP-binding_3_2962_4495 | Aryl-CoA ligase | 0.42 | cinnamate and succinylbenzoate derivatives | 0.27 |
| gb10_c02489_NODE_24...region001_haloAMP_AMP-binding_1_0_1551 | Aryl-CoA ligase | 0.42 | cinnamate and succinylbenzoate derivatives | 0.27 |
| DOM044_c02058_NODE_20...region001_haloAMP_AMP-binding_9_6580_8362 | Aryl-CoA ligase | 0.42 | cinnamate and succinylbenzoate derivatives | 0.27 |
| gb5_2_c09272_NODE_92...region001_haloAMP_AMP-binding_5_6167_7748 | Aryl-CoA ligase | 0.42 | cinnamate and succinylbenzoate derivatives | 0.27 |
| DOM43_c00933_NODE_93...region001_haloAMP_AMP-binding_17_17919_19428 | Aryl-CoA ligase | 0.43 | cinnamate and succinylbenzoate derivatives | 0.27 |
| Pf11_c16311_NODE_16...region001_haloAMP_AMP-binding_1_0_1470 | Aryl-CoA ligase | 0.41 | cinnamate and succinylbenzoate derivatives | 0.24 |
| Pf4_c01746_NODE_17...region001_haloAMP_AMP-binding_1_344_1835 | Aryl-CoA ligase | 0.41 | cinnamate and succinylbenzoate derivatives | 0.24 |
| Pf7_c22554_NODE_22...region001_haloAMP_AMP-binding_1_87_1578 | Aryl-CoA ligase | 0.41 | cinnamate and succinylbenzoate derivatives | 0.24 |
| Aply22_bin.20.strict_NODE_213_length_4775_cov_6.472669.region001_haloAMP_AMP-binding_4_3197_4688 | Aryl-CoA ligase | 0.41 | cinnamate and succinylbenzoate derivatives | 0.24 |
| Pf8_c16919_NODE_16...region001_haloAMP_AMP-binding_3_2598_4089 | Aryl-CoA ligase | 0.41 | cinnamate and succinylbenzoate derivatives | 0.24 |
| Pf12_bin.52.strict_NODE_4_length_127629_cov_21.520321.region001_haloAMP_AMP-binding_17_16001_17492 | Aryl-CoA ligase | 0.41 | cinnamate and succinylbenzoate derivatives | 0.24 |
| Aply23_bin.9.strict_NODE_85_length_10472_cov_3.930498.region001_haloAMP_AMP-binding_4_2789_4280 | Aryl-CoA ligase | 0.41 | cinnamate and succinylbenzoate derivatives | 0.24 |
| Pf10_c10573_NODE_10...region001_haloAMP_AMP-binding_1_85_1576 | Aryl-CoA ligase | 0.41 | cinnamate and succinylbenzoate derivatives | 0.24 |
| DOM011_c01245_NODE_12...region001_haloAMP_AMP-binding_14_16222_17731 | Aryl-CoA ligase | 0.42 | cinnamate and succinylbenzoate derivatives | 0.27 |
| Pf11_c00891_NODE_89...region001_haloAMP_AMP-binding_25_20000_21491 | Aryl-CoA ligase | 0.46 | cinnamate and succinylbenzoate derivatives | 0.28 |
| Pf9_c05741_NODE_57...region001_haloAMP_AMP-binding_11_9051_9366 | Aryl-CoA ligase | 0.56 | cinnamate and succinylbenzoate derivatives | 0.35 |

| | | | | |
|---|---|---|---|---|
| Pf5_c01370_NODE_13...region001_haloAMP_AMP-binding_16_16011_17502 | Aryl-CoA ligase | 0.41 | cinnamate and succinylbenzoate derivatives | 0.24 |
| Pf9_c28236_NODE_28...region001_haloAMP_AMP-binding_2_557_2048 | Aryl-CoA ligase | 0.41 | cinnamate and succinylbenzoate derivatives | 0.24 |
| Aply21_c08313_NODE_83...region001_haloAMP_AMP-binding_4_2959_3403 | Aryl-CoA ligase | 0.55 | cinnamate and succinylbenzoate derivatives | 0.32 |
| DOM33_c00105_NODE_10...region001_haloAMP_AMP-binding_2_1113_2622 | Aryl-CoA ligase | 0.42 | cinnamate and succinylbenzoate derivatives | 0.27 |
| DOM14B_c25243_NODE_25...region001_haloAMP_AMP-binding_1_57_1575 | Aryl-CoA ligase | 0.41 | cinnamate and succinylbenzoate derivatives | 0.24 |
| DOM026_c12884_NODE_12...region001_haloAMP_AMP-binding_5_2363_3872 | Aryl-CoA ligase | 0.41 | cinnamate and succinylbenzoate derivatives | 0.24 |
| Pf5_bin.39.strict_NODE_3_length_52011_cov_5.124158.region001_haloAMP_AMP-binding_3_2028_3519 | Aryl-CoA ligase | 0.46 | cinnamate and succinylbenzoate derivatives | 0.28 |
| gb1_bin.48.strict_NODE_59_length_14622_cov_7.645239.region001_haloAMP_AMP-binding_13_13050_14622 | Aryl-CoA ligase | 0.42 | cinnamate and succinylbenzoate derivatives | 0.27 |
| DOM045_bin.2.strict_NODE_2_length_313427_cov_20.378002.region001_haloAMP_AMP-binding_11_10546_12055 | Aryl-CoA ligase | 0.42 | cinnamate and succinylbenzoate derivatives | 0.27 |
| Aply16_c09216_NODE_92...region001_haloAMP_AMP-binding_4_2977_3976 | Aryl-CoA ligase | 0.5 | cinnamate and succinylbenzoate derivatives | 0.26 |
| gb4_2_bin.24.strict_NODE_46_length_17554_cov_6.212451.region001_haloAMP_AMP-binding_12_12732_14685 | Aryl-CoA ligase | 0.42 | cinnamate and succinylbenzoate derivatives | 0.27 |
| gb2_2_bin.25.strict_NODE_63_length_13684_cov_7.323436.region001_haloAMP_AMP-binding_12_12703_13684 | Aryl-CoA ligase | 0.5 | cinnamate and succinylbenzoate derivatives | 0.3 |
| DOM43_bin.27.strict_NODE_22_length_35237_cov_8.092804.region001_haloAMP_AMP-binding_2_642_2151 | Aryl-CoA ligase | 0.43 | cinnamate and succinylbenzoate derivatives | 0.27 |
| gb6_bin.52.strict_NODE_30_length_20515_cov_7.543987.region001_haloAMP_AMP-binding_17_19651_20515 | Aryl-CoA ligase | 0.32 | cinnamate and succinylbenzoate derivatives | 0.13 |
| gb8_2_c49583_NODE_49...region001_haloAMP_AMP-binding_2_1257_2838 | Aryl-CoA ligase | 0.42 | cinnamate and succinylbenzoate derivatives | 0.27 |

**Table S7.** PARAS extracted and aligned Stachelhaus and extended active site residues of A-domains in CC A, Genieuos alignment statistics and WebLogos.

| A-domain | DomainStachelhaus | Active Site |
|---|---|---|
| Aply16_c09216_NODE_92...region001_haloAMP_AMP-binding_4_2977_3976 | AVMFGGHPL | SSTVHAVGLMIQFLGVGGATDHYGQSETGPLTIL |
| Aply21_c08313_NODE_83...region001_haloAMP_AMP-binding_4_2959_3403 | ---YLV-VR | -----------AYALVVDE----GRIGTEVRSAV |
| Aply22_bin.20.strict_NODE_213_length_4775_cov_6.472669.region001_haloAMP_AMP-binding_4_3197_4688 | AVMFGGHPL | SSTVHAVGLMIQFLGVGGATDHYGQSETGPLTIL |
| Aply23_bin.9.strict_NODE_85_length_10472_cov_3.930498.region001_haloAMP_AMP-binding_4_2789_4280 | AVMFGGHPL | SSTVHAVGLMIQFLGVGGATDHYGQSETGPLTIL |
| DOM011_c01245_NODE_12...region001_haloAMP_AMP-binding_14_16222_17731 | AVMFGGHPL | SSTVHAVGLMIQFLGVGGATDHYGQSETGPLTIL |
| DOM026_c12884_NODE_12...region001_haloAMP_AMP-binding_5_2363_3872 | AVMFGGHPL | SSTVHAVGLMIQFLGVGGATDHYGQSETGPLTIL |
| DOM044_c02058_NODE_20...region001_haloAMP_AMP-binding_9_6580_8362 | AVMFGGHPL | SSTVHAVGLMIQFLGVGGATDHYGQSETGPLTIL |
| DOM045_bin.2.strict_NODE_2_length_313427_cov_20.378002.region001_haloAMP_AMP-binding_11_10546_12055 | AVMFGGHPL | SSTVHAVGLMIQFLGVGGATDHYGQSETGPLTIL |
| DOM049_bin.11.strict_NODE_225_length_5496_cov_4.161100.region001_haloAMP_AMP-binding_3_2962_4495 | AVMFGGHPL | SSTVHAVGLMIQFLGVGGATDHYGQSETGPLTIL |
| DOM14B_c25243_NODE_25...region001_haloAMP_AMP-binding_1_57_1575 | AVMFGGHPL | SSTTHAVGLMIAFLGVGGATDHYGQSETGPLTIL |
| DOM33_c00105_NODE_10...region001_haloAMP_AMP-binding_2_1113_2622 | AVMFGGHPL | SSTVHAVGLMIQFLGVGGATDHYGQSETGPLTIL |
| DOM43_bin.27.strict_NODE_22_length_35237_cov_8.092804.region001_haloAMP_AMP-binding_2_642_2151 | AVMFGGHPL | STTTHAVGLMIAFLGVGGATDHYGQSETGPLTIL |
| DOM43_c00933_NODE_93...region001_haloAMP_AMP-binding_17_17919_19428 | AVMFGGHPL | STTTHAVGLMIAFLGVGGATDHYGQSETGPLTIL |
| Pf10_c10573_NODE_10...region001_haloAMP_AMP-binding_1_85_1576 | AVMFGGHPL | SSTVHAVGLMIQFLGVGGATDHYGQSETGPLTIL |
| Pf11_c00891_NODE_89...region001_haloAMP_AMP-binding_25_20000_21491 | AVMFGGHPL | STTTHAVGLMIAFLGVGGATDHYGQSETGPLTIL |
| Pf11_c16311_NODE_16...region001_haloAMP_AMP-binding_1_0_1470 | AVMFGGHPL | SSTVHAVGLMIQFLGVGGATDHYGQSETGPLTIL |
| Pf12_bin.52.strict_NODE_4_length_127629_cov_21.520321.region001_haloAMP_AMP-binding_17_16001_17492 | AVMFGGHPL | SSTVHAVGLMIQFLGVGGATDHYGQSETGPLTIL |
| Pf4_c01746_NODE_17...region001_haloAMP_AMP-binding_1_344_1835 | AVMFGGHPL | SSTVHAVGLMIQFLGVGGATDHYGQSETGPLTIL |
| Pf5_bin.39.strict_NODE_3_length_52011_cov_5.124158.region001_haloAMP_AMP-binding_3_2028_3519 | AVMFGGHPL | STTTHAVGLMIAFLGVGGATDHYGQSETGPLTIL |
| Pf5_c01370_NODE_13...region001_haloAMP_AMP-binding_16_16011_17502 | AVMFGGHPL | SSTVHAVGLMIQFLGVGGATDHYGQSETGPLTIL |

| | | |
|---|---|---|
| Pf6_c44645_NODE_44...region001_haloAMP_AMP-binding_1_304_1795 | AVMFGGHPL | SSTVHAVGLMIQFLGVGGATDHYGQSETGPLTIL |
| Pf7_c22554_NODE_22...region001_haloAMP_AMP-binding_1_87_1578 | AVMFGGHPL | SSTVHAVGLMIQFLGVGGATDHYGQSETGPLTIL |
| Pf8_c16919_NODE_16...region001_haloAMP_AMP-binding_3_2598_4089 | AVMFGGHPL | SSTVHAVGLMIQFLGVGGATDHYGQSETGPLTIL |
| Pf9_c05741_NODE_57...region001_haloAMP_AMP-binding_11_9051_9366 | ---L----- | -----------GLY-------------------- |
| Pf9_c28236_NODE_28...region001_haloAMP_AMP-binding_2_557_2048 | AVMFGGHPL | SSTVHAVGLMIQFLGVGGATDHYGQSETGPLTIL |
| gb10_c02489_NODE_24...region001_haloAMP_AMP-binding_1_0_1551 | AVMFGGHPL | SSTTHAVGLMIAFLGVGGATDHYGQSETGPLTIL |
| gb1_bin.48.strict_NODE_59_length_14622_cov_7.645239.region001_haloAMP_AMP-binding_13_13050_14622 | AVMFGGHPL | SSTTHAVGLMIAFLGVGGATDHYGQSETGPLTIL |
| gb2_2_bin.25.strict_NODE_63_length_13684_cov_7.323436.region001_haloAMP_AMP-binding_12_12703_13684 | AVMFGGH-- | SSTTHAVGLMIAFLGVGGATDHYGQSET------ |
| gb4_2_bin.24.strict_NODE_46_length_17554_cov_6.212451.region001_haloAMP_AMP-binding_12_12732_14685 | AVMFGGHPL | SSTTHAVGLMIAFLGVGGATDHYGQSETGPLTIL |
| gb5_2_c09272_NODE_92...region001_haloAMP_AMP-binding_5_6167_7748 | AVMFGGHPL | SSTTHAVGLMIAFLGVGGATDHYGQSETGPLTIL |
| gb6_bin.52.strict_NODE_30_length_20515_cov_7.543987.region001_haloAMP_AMP-binding_17_19651_20515 | AVMFGG--- | SSTTHAVGLMIAFLGVGGA--------------- |
| gb8_2_c49583_NODE_49...region001_haloAMP_AMP-binding_2_1257_2838 | AVMFGGHPL | SSTTHAVGLMIAFLGVGGATDHYGQSETGPLTIL |
| gb9_c07451_NODE_74...region001_haloAMP_AMP-binding_5_6089_7670 | AVMFGGHPL | SSTTHAVGLMIAFLGVGGATDHYGQSETGPLTIL |
| **PARAS active site statistics (ungapped only):** | | |
| Length (mean): 34 aa (34 codons) | | |
| Sequences: 29 | | |
| Identical Sites: 31 (91.2%) | | |
| Pairwise Identity: 96.4% | | |
| Pairwise Positive (BLSM62): 97.1% | | |
| 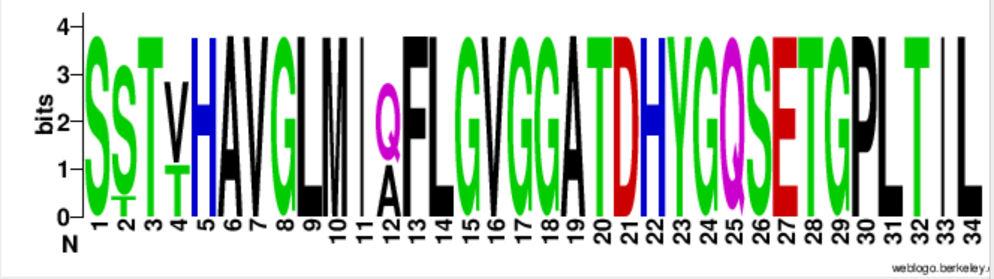 | | |
| **Stachelhaus residues statistics (ungapped only):** | | |

| |
|---|
| Length (mean): 9 aa (9 codons) |
| Sequences: 29 |
| Identical Sites: 9 (100.0%) |
| Pairwise Identity: 100.0% |
| Pairwise Positive (BLSM62): 100.0% |
| 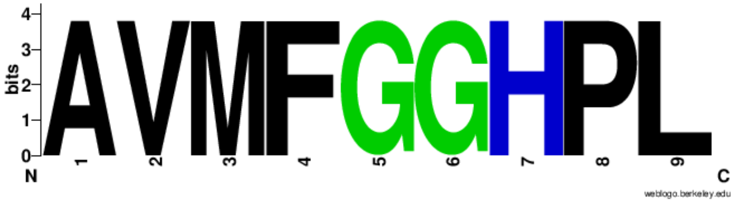 |

**Table S8.** Acidobacteria Bin61 MAG information.

| Genome | Completeness | Contamination | Strain heterogeneity | Length | N50 | Taxonomic classification |
|---|---|---|---|---|---|---|
| Aply22_bin.20.strict.fa | 87.29 | 0.96 | 33.33 | 3441866 | 17362 | d__Bacteria;p__Acidobacteriota;c__Bin61;o__Bin61;f__Bin61;g__Bin61;s__Bin61 sp002238705 |
| Aply23_bin.9.strict.fa | 71.12 | 1.29 | 0.0 | 3019528 | 9569 | d__Bacteria;p__Acidobacteriota;c__Bin61;o__Bin61;f__Bin61;g__Bin61;s__Bin61 sp002238705 |
| DOM007A_bin.35.strict.fa | 88.84 | 0.85 | 0.0 | 2705393 | 1384733 | d__Bacteria;p__Acidobacteriota;c__Bin61;o__Bin61;f__Bin61;g__Bin61;s__ |
| DOM007B_bin.2.strict.fa | 88.84 | 0.85 | 0.0 | 2766179 | 486662 | d__Bacteria;p__Acidobacteriota;c__Bin61;o__Bin61;f__Bin61;g__Bin61;s__ |
| DOM011_bin.33.orig.fa | 92.68 | 0.85 | 0.0 | 4410379 | 48964 | d__Bacteria;p__Acidobacteriota;c__Bin61;o__Bin61;f__Bin61;g__Bin61;s__ |
| DOM012A_bin.31.strict.fa | 83.56 | 1.78 | 0.0 | 2963632 | 16420 | d__Bacteria;p__Acidobacteriota;c__Bin61;o__Bin61;f__Bin61;g__Bin61;s__ |
| DOM013_bin.40.orig.fa | 80.17 | 1.78 | 33.33 | 2537336 | 8169 | d__Bacteria;p__Acidobacteriota;c__Bin61;o__Bin61;f__Bin61;g__Bin61;s__ |
| DOM015_bin.63.strict.fa | 85.09 | 2.14 | 0.0 | 2631338 | 12893 | d__Bacteria;p__Acidobacteriota;c__Bin61;o__Bin61;f__Bin61;g__Bin61;s__ |

| | | | | | | |
|---|---|---|---|---|---|---|
| DOM016_bin.38.permissive.fa | 87.98 | 0.9 | 0.0 | 2708755 | 492822 | d__Bacteria;p__Acidobacteriota;c__Bin61;o__Bin61;f__Bin61;g__Bin61;s__ |
| DOM026_bin.53.orig.fa | 82.05 | 0.85 | 0.0 | 2918552 | 14826 | d__Bacteria;p__Acidobacteriota;c__Bin61;o__Bin61;f__Bin61;g__Bin61;s__ |
| DOM044_bin.47.orig.fa | 92.74 | 2.56 | 25.0 | 4505327 | 47868 | d__Bacteria;p__Acidobacteriota;c__Bin61;o__Bin61;f__Bin61;g__Bin61;s__ |
| DOM045_bin.2.strict.fa | 91.88 | 2.19 | 0.0 | 4273376 | 120250 | d__Bacteria;p__Acidobacteriota;c__Bin61;o__Bin61;f__Bin61;g__Bin61;s__ |
| DOM049_bin.11.strict.fa | 83.01 | 0.85 | 0.0 | 3504759 | 14813 | d__Bacteria;p__Acidobacteriota;c__Bin61;o__Bin61;f__Bin61;g__Bin61;s__ |
| DOM057_bin.38.orig.fa | 85.42 | 0.85 | 0.0 | 2642392 | 322731 | d__Bacteria;p__Acidobacteriota;c__Bin61;o__Bin61;f__Bin61;g__Bin61;s__ |
| DOM10_bin.63.orig.fa | 86.75 | 0.0 | 0.0 | 3532383 | 141441 | d__Bacteria;p__Acidobacteriota;c__Bin61;o__Bin61;f__Bin61;g__Bin61;s__ |
| DOM14B_bin.52.orig.fa | 90.12 | 0.85 | 0.0 | 4101197 | 26286 | d__Bacteria;p__Acidobacteriota;c__Bin61;o__Bin61;f__Bin61;g__Bin61;s__ |
| DOM33_bin.71.orig.fa | 91.88 | 0.85 | 0.0 | 4124016 | 167581 | d__Bacteria;p__Acidobacteriota;c__Bin61;o__Bin61;f__Bin61;g__Bin61;s__ |
| DOM40_bin.43.strict.fa | 89.79 | 0.85 | 0.0 | 3487366 | 40484 | d__Bacteria;p__Acidobacteriota;c__Bin61;o__Bin61;f__Bin61;g__Bin61;s__ |
| DOM43_bin.14.orig.fa | 59.07 | 5.23 | 11.11 | 3539402 | 2915 | d__Bacteria;p__Acidobacteriota;c__Bin61;o__Bin61;f__SMYC01;g__;s__ |
| DOM43_bin.27.strict.fa | 87.87 | 3.42 | 20.0 | 3216245 | 27215 | d__Bacteria;p__Acidobacteriota;c__Bin61;o__Bin61;f__Bin61;g__Bin61;s__ |
| gb1_bin.48.strict.fa | 87.18 | 0.91 | 0.0 | 3121056 | 12235 | d__Bacteria;p__Acidobacteriota;c__Bin61;o__Bin61;f__Bin61;g__Bin61;s__ |
| gb10_bin.24.orig.fa | 72.81 | 0.0 | 0.0 | 2826534 | 5528 | d__Bacteria;p__Acidobacteriota;c__Bin61;o__Bin61;f__Bin61;g__Bin61;s__ |
| gb126_bin.28.strict.fa | 83.05 | 6.03 | 0.0 | 3987243 | 9045 | d__Bacteria;p__Acidobacteriota;c__Bin61;o__Bin61;f__SMYC01;g__SMYC01;s__ |
| gb126_bin.4.strict.fa | 54.74 | 0.0 | 0.0 | 2194644 | 4614 | d__Bacteria;p__Acidobacteriota;c__Bin61;o__Bin61;f__Bin61;g__Bin61;s__ |

| | | | | | | |
|---|---|---|---|---|---|---|
| gb126_bin.57.orig.fa | 50.68 | 4.13 | 50.0 | 3028711 | 2666 | d__Bacteria;p__Acidobacteriota;c__Bin61;o__Bin61;f__SMYC01;g__;s__ |
| gb2_2_bin.25.strict.fa | 83.12 | 0.85 | 0.0 | 3072413 | 12488 | d__Bacteria;p__Acidobacteriota;c__Bin61;o__Bin61;f__Bin61;g__Bin61;s__ |
| gb278_bin.12.strict.fa | 58.95 | 7.26 | 0.0 | 2733024 | 2974 | d__Bacteria;p__Acidobacteriota;c__Bin61;o__Bin61;f__SMYC01;g__;s__ |
| gb278_bin.29.strict.fa | 65.93 | 2.14 | 62.5 | 3443931 | 6145 | d__Bacteria;p__Acidobacteriota;c__Bin61;o__Bin61;f__SMYC01;g__SMYC01;s__ |
| gb278_bin.7.strict.fa | 73.36 | 0.11 | 0.0 | 2574323 | 5603 | d__Bacteria;p__Acidobacteriota;c__Bin61;o__Bin61;f__Bin61;g__Bin61;s__ |
| gb3_2_bin.12.strict.fa | 82.48 | 0.85 | 0.0 | 3020087 | 12511 | d__Bacteria;p__Acidobacteriota;c__Bin61;o__Bin61;f__Bin61;g__Bin61;s__ |
| gb305_bin.4.strict.fa | 81.31 | 6.93 | 0.0 | 5007556 | 9998 | d__Bacteria;p__Acidobacteriota;c__Bin61;o__Bin61;f__SMYC01;g__SMYC01;s__ |
| gb305_bin.59.strict.fa | 73.65 | 0.85 | 0.0 | 2407354 | 6046 | d__Bacteria;p__Acidobacteriota;c__Bin61;o__Bin61;f__Bin61;g__Bin61;s__ |
| gb4_2_bin.24.strict.fa | 83.45 | 1.71 | 0.0 | 3165550 | 13053 | d__Bacteria;p__Acidobacteriota;c__Bin61;o__Bin61;f__Bin61;g__Bin61;s__ |
| gb5_2_bin.40.orig.fa | 82.48 | 0.85 | 0.0 | 2884449 | 5273 | d__Bacteria;p__Acidobacteriota;c__Bin61;o__Bin61;f__Bin61;g__Bin61;s__ |
| gb6_bin.52.strict.fa | 88.68 | 0.91 | 0.0 | 3095017 | 12948 | d__Bacteria;p__Acidobacteriota;c__Bin61;o__Bin61;f__Bin61;g__Bin61;s__ |
| gb7_bin.26.strict.fa | 81.41 | 0.85 | 0.0 | 2910222 | 8040 | d__Bacteria;p__Acidobacteriota;c__Bin61;o__Bin61;f__Bin61;g__Bin61;s__ |
| gb8_2_bin.28.orig.fa | 72.98 | 0.0 | 0.0 | 2901044 | 5907 | d__Bacteria;p__Acidobacteriota;c__Bin61;o__Bin61;f__Bin61;g__Bin61;s__ |
| gb9_bin.7.orig.fa | 83.76 | 0.85 | 0.0 | 2845379 | 6180 | d__Bacteria;p__Acidobacteriota;c__Bin61;o__Bin61;f__Bin61;g__Bin61;s__ |
| Pf10_bin.25.orig.fa | 87.98 | 0.85 | 0.0 | 3227732 | 15336 | d__Bacteria;p__Acidobacteriota;c__Bin61;o__Bin61;f__Bin61;g__Bin61;s__ |
| Pf11_bin.29.orig.fa | 66.67 | 0.0 | 0.0 | 2216665 | 42881 | d__Bacteria;p__Acidobacteriota;c__Bin61;o__Bin61;f__Bin61;g__Bin61;s__ |

| | | | | | | |
|---|---|---|---|---|---|---|
| Pf11_bin.41.strict.fa | 52.63 | 1.75 | 100.0 | 1829438 | 9676 | d__Bacteria;p__Acidobacteriota;c__Bin61;o__Bin61;f__Bin61;g__Bin61;s__ |
| Pf12_bin.52.strict.fa | 91.4 | 0.9 | 0.0 | 3790910 | 65216 | d__Bacteria;p__Acidobacteriota;c__Bin61;o__Bin61;f__Bin61;g__Bin61;s__ |
| Pf4_bin.17.orig.fa | 91.83 | 0.85 | 0.0 | 3822207 | 42050 | d__Bacteria;p__Acidobacteriota;c__Bin61;o__Bin61;f__Bin61;g__Bin61;s__ |
| Pf5_bin.12.strict.fa | 91.4 | 0.85 | 0.0 | 3720584 | 55861 | d__Bacteria;p__Acidobacteriota;c__Bin61;o__Bin61;f__Bin61;g__Bin61;s__Bin61 sp002238705 |
| Pf5_bin.39.strict.fa | 90.64 | 2.02 | 20.0 | 2993178 | 21997 | d__Bacteria;p__Acidobacteriota;c__Bin61;o__Bin61;f__Bin61;g__Bin61;s__ |
| Pf6_bin.12.orig.fa | 87.13 | 0.85 | 0.0 | 3112957 | 12796 | d__Bacteria;p__Acidobacteriota;c__Bin61;o__Bin61;f__Bin61;g__Bin61;s__ |
| Pf8_bin.35.orig.fa | 88.84 | 0.85 | 0.0 | 3370558 | 14062 | d__Bacteria;p__Acidobacteriota;c__Bin61;o__Bin61;f__Bin61;g__Bin61;s__ |
| Pf9_bin.26.orig.fa | 82.0 | 1.71 | 20.0 | 2994752 | 21154 | d__Bacteria;p__Acidobacteriota;c__Bin61;o__Bin61;f__Bin61;g__Bin61;s__ |
| Pf9_bin.51.orig.fa | 74.07 | 1.85 | 20.0 | 2429284 | 5242 | d__Bacteria;p__Acidobacteriota;c__Bin61;o__Bin61;f__Bin61;g__Bin61;s__ |

**Table S9.** CC A and SwissProt single domain P450 CYP/KO annotation**.**

| P450 | KO | e-value | bitscore | CYP | e-value2 | bitscore3 |
|---|---|---|---|---|---|---|
| **gb9_c07451_NODE_74...region001_haloAMP_p450_1_184_1519** | K21164 | 1.1e-127 | 427.2 | CYP253A1 | 5.27e-91 | 282 |
| gb9_c07451_NODE_74...region001_haloAMP_p450_2_1772_3116 | K21164 | 1.9e-110 | 370.5 | CYP253B2 | 1.13e-86 | 271 |
| gb10_c02489_NODE_24...region001_haloAMP_p450_4_4524_5868 | K21164 | 6.6e-111 | 372.0 | CYP253B2 | 7.90e-86 | 270 |
| gb10_c02489_NODE_24...region001_haloAMP_p450_5_6121_7456 | K21164 | 1.1e-127 | 427.2 | CYP253A1 | 5.27e-91 | 282 |
| gb10_c02489_NODE_24...region001_haloAMP_p450_6_7699_8968 | K21164 | 3.9e-107 | 359.6 | CYP253A1 | 6.08e-76 | 243 |

| | | | | | | |
|---|---|---|---|---|---|---|
| DOM044_c02058_NODE_20...region001_haloAMP_p450_13_11922_13269 | K21164 | 9.9e-127 | 424.2 | CYP253B2 | 4.53e-95 | 293 |
| gb5_2_c09272_NODE_92...region001_haloAMP_p450_1_262_1597 | K21164 | 1.1e-127 | 427.2 | CYP253A1 | 5.27e-91 | 282 |
| gb5_2_c09272_NODE_92...region001_haloAMP_p450_2_1850_3194 | K21164 | 7.2e-112 | 375.2 | CYP253B2 | 1.42e-85 | 269 |
| DOM43_c00933_NODE_93...region001_haloAMP_p450_19_22387_23731 | K21164 | 1.1e-123 | 414.1 | CYP253B1 | 7.35e-90 | 280 |
| Pf4_c01746_NODE_17...region001_haloAMP_p450_4_4545_5895 | K21164 | 1.1e-126 | 424.0 | CYP197A1 | 1.03e-96 | 296 |
| Aply22_bin.20.strict_NODE_213_length_4775_cov_6.472669.region001_haloAMP_p450_1_2_482 | K21164 | 1.6e-56 | 192.8 | CYP253C1 | 9.62e-45 | 150 |
| Pf12_bin.52.strict_NODE_4_length_127629_cov_21.520321.region001_haloAMP_p450_14_11936_13286 | K21164 | 3.2e-126 | 422.5 | CYP197A1 | 2.10e-96 | 296 |
| Pf10_c10573_NODE_10...region001_haloAMP_p450_4_4291_5641 | K21164 | 9,00E-127 | 424.3 | CYP197A1 | 1.32e-96 | 296 |
| DOM011_c01245_NODE_12...region001_haloAMP_p450_17_20451_21801 | K21164 | 1.7e-126 | 423.4 | CYP197A1 | 6.16e-94 | 289 |
| Pf11_c00891_NODE_89...region001_haloAMP_p450_28_24188_25535 | K21164 | 5.2e-125 | 418.5 | CYP253B2 | 1.22e-92 | 287 |
| Pf9_c05741_NODE_57...region001_haloAMP_p450_8_5005_6352 | K21164 | 1.7e-123 | 413.5 | CYP253B2 | 7.12e-95 | 293 |
| Pf5_c01370_NODE_13...region001_haloAMP_p450_13_11947_13297 | K21164 | 9,00E-127 | 424.3 | CYP197A1 | 1.32e-96 | 296 |
| Aply21_c08313_NODE_83...region001_haloAMP_p450_1_1_244 | K15001 | 4.2e-23 | 82.1 | CYP262A1 | 6.05e-21 | 83.2 |
| **DOM33_c00105_NODE_10...region001_haloAMP_p450_5_5344_6694** | K21164 | 2.5e-126 | 422.8 | CYP253B2 | 1.69e-95 | 295 |
| DOM14B_c25243_NODE_25...region001_haloAMP_p450_4_4321_5671 | K21164 | 2,00E-125 | 419.9 | CYP253A2 | 1.46e-93 | 288 |
| gb1_bin.48.strict_NODE_59_length_14622_cov_7.645239.region001_haloAMP_p450_8_5644_6913 | K21164 | 5.8e-107 | 359.0 | CYP253A1 | 6.08e-76 | 243 |

| | | | | | | |
|---|---|---|---|---|---|---|
| gb1_bin.48.strict_NODE_59_length_14622_cov_7.645239.region001_haloAMP_p450_9_7157_8492 | K21164 | 1.1e-127 | 427.2 | CYP253A1 | 5.27e-91 | 282 |
| gb1_bin.48.strict_NODE_59_length_14622_cov_7.645239.region001_haloAMP_p450_10_8745_10089 | K21164 | 2.1e-111 | 373.7 | CYP253B2 | 2.28e-86 | 271 |
| DOM045_bin.2.strict_NODE_2_length_313427_cov_20.378002.region001_haloAMP_p450_7_5633_6983 | K21164 | 5.9e-126 | 421.6 | CYP253B2 | 5.78e-94 | 290 |
| Aply16_c09216_NODE_92...region001_haloAMP_p450_1_2_263 | K15001 | 6.6e-28 | 98.0 | CYP253C1 | 1.33e-22 | 88.2 |
| **gb4_2_bin.24.strict_NODE_46_length_17554_cov_6.212451.region001_haloAMP_p450_7_5314_6583** | K21164 | 3.9e-107 | 359.6 | CYP253A1 | 6.08e-76 | 243 |
| gb4_2_bin.24.strict_NODE_46_length_17554_cov_6.212451.region001_haloAMP_p450_8_6827_8162 | K21164 | 1.1e-127 | 427.2 | CYP253A1 | 5.27e-91 | 282 |
| gb4_2_bin.24.strict_NODE_46_length_17554_cov_6.212451.region001_haloAMP_p450_9_8415_9759 | K21164 | 2.5e-111 | 373.4 | CYP253B2 | 2.07e-86 | 271 |
| gb2_2_bin.25.strict_NODE_63_length_13684_cov_7.323436.region001_haloAMP_p450_7_5285_6554 | K21164 | 3.9e-107 | 359.6 | CYP253A1 | 6.08e-76 | 243 |
| gb2_2_bin.25.strict_NODE_63_length_13684_cov_7.323436.region001_haloAMP_p450_8_6798_8133 | K21164 | 1.1e-127 | 427.2 | CYP253A1 | 5.27e-91 | 282 |
| gb2_2_bin.25.strict_NODE_63_length_13684_cov_7.323436.region001_haloAMP_p450_9_8386_9730 | K21164 | 1.9e-110 | 370.5 | CYP253B2 | 1.13e-86 | 271 |
| DOM43_bin.27.strict_NODE_22_length_35237_cov_8.092804.region001_haloAMP_p450_5_4872_6216 | K21164 | 5.5e-123 | 411.8 | CYP253B1 | 2.26e-90 | 281 |
| gb6_bin.52.strict_NODE_30_length_20515_cov_7.543987.region001_haloAMP_p450_12_12245_13514 | K21164 | 3.9e-107 | 359.6 | CYP253A1 | 6.08e-76 | 243 |
| gb6_bin.52.strict_NODE_30_length_20515_cov_7.543987.region001_haloAMP_p450_13_13758_15093 | K21164 | 1.1e-127 | 427.2 | CYP253A1 | 5.27e-91 | 282 |
| gb6_bin.52.strict_NODE_30_length_20515_cov_7.543987.region001_haloAMP_p450_14_15346_16690 | K21164 | 1.9e-110 | 370.5 | CYP253B2 | 1.13e-86 | 271 |
| DOM015_c02570_NODE_25...region001_haloAMP_p450_2_266_1178 | K21164 | 1.5e-28 | 100.6 | CYP286A1 | 1.95e-43 | 152 |
| DOM044_c01110_NODE_11...region001_haloAMP_p450_22_13908_15156 | K16593 | 1.6e-127 | 426.3 | CYP107AZ1 | 1.34e-113 | 337 |

| | | | | | | |
|---|---|---|---|---|---|---|
| DOM33_c01507_NODE_15...region001_haloAMP_p450_21_26204_27473 | K16593 | 2.4e-108 | 363.2 | CYP1011B1 | 1.11e-103 | 312 |
| DOM33_c01988_NODE_19...region001_haloAMP_p450_23_19387_20641 | K16593 | 5.6e-127 | 424.5 | CYP107AZ1 | 7.73e-115 | 340 |
| PLPH_bmp7_AHA34040.1_BGC0000890 | K17814 | 7.2e-102 | 342.2 | Cyp2r1 | 2.06e-59 | 202 |
| PL2TA16_bmp7_ESP90850.1_BGC0000891 | K17814 | 3.9e-98 | 329.9 | CYP1A1 | 9.37e-59 | 201 |
| MARME_bmp7_WP_013663197.1_BGC0001465 | K17814 | 1.9e-92 | 311.1 | CYP1A1 | 1.37e-56 | 195 |
| GUM202_bmp7_MBO1347623.1 | K23139 | 2.9e-180 | 600.1 | CYP110C6 | 0.0 | 514 |
| GUM202_bmp12_MBO1347626.1 | K17814 | 4.3e-120 | 402.3 | CYP1C2 | 2.43e-71 | 234 |
| sp\|A0R4Q6\|CP142_MYCS2 | K16046 | 3.3e-216 | 718.0 | CYP142A2 | 0.0 | 819 |
| sp\|A0R4Y3\|CP125_MYCS2 | K15981 | 2.4e-212 | 705.4 | CYP125A3 | 0.0 | 883 |
| sp\|A4F7P2\|ERYC2_SACEN | K15997 | 7.4e-228 | 756.0 | CYP131A1 | 2.49e-24 | 102 |
| sp\|B4XY99\|AZIB1_STREG | K21376 | 4.4e-291 | 964.7 | CYP1019A1 | 1.23e-49 | 171 |
| sp\|C4B644\|CPVDH_PSEAH | K24389 | 3.7e-240 | 796.9 | CYP107BR1 | 0.0 | 808 |
| sp\|C9K1X6\|COTB3_STRMJ | K22997 | 9,00E-249 | 826.1 | CYP183B1 | 2.90e-129 | 380 |
| sp\|C9K1X7\|COTB4_STRMJ | K22998 | 0 | 1016.7 | CYP183A2 | 5.24e-133 | 389 |
| sp\|D5E3H2\|CP107_PRIM1 | K16593 | 1.4e-115 | 387.1 | CYP267B1 | 2.70e-108 | 323 |
| sp\|E3VWI3\|PNTM_STRAE | K17476 | 2.9e-259 | 859.8 | CYP161C2 | 0.0 | 798 |
| sp\|E3VWJ9\|PENM_STREX | K17476 | 1.5e-258 | 857.4 | CYP161C3 | 0.0 | 809 |
| sp\|I3DZK9\|CYPC_BACMM | K15629 | 5.4e-190 | 632.9 | CYP152A1 | 1.15e-162 | 462 |
| sp\|O08469\|CPXY_BACSU | K16593 | 5.6e-124 | 414.7 | CYP107J1 | 0.0 | 845 |
| sp\|O31440\|CYPC_BACSU | K15629 | 1.6e-192 | 641.2 | CYP152A1 | 0.0 | 870 |
| sp\|O31785\|PKSS_BACSU | K15468 | 6.3e-269 | 891.9 | CYP107K1 | 0.0 | 765 |
| sp\|O34374\|YJIB_BACSU | K23138 | 3.7e-185 | 616.0 | CYP109B1 | 0.0 | 821 |
| sp\|O34926\|CYPX_BACSU | K17474 | 2.8e-221 | 734.9 | CYP134A1 | 0.0 | 840 |
| sp\|O87605\|PIKC_STRVZ | K16006 | 1.5e-295 | 979.9 | CYP107L1 | 0.0 | 829 |
| sp\|P00183\|CPXA_PSEPU | K21569 | 9.9e-216 | 716.3 | CYP101A1 | 0.0 | 863 |
| sp\|P0A513\|CP51_MYCBO | K05917 | 3.1e-176 | 588.1 | CYP51B1 | 0.0 | 930 |

| | | | | | | |
|---|---|---|---|---|---|---|
| sp\|P0A515\|CP121_MYCBO | K17483 | 6.7e-191 | 634.7 | CYP121A1 | 0.0 | 800 |
| sp\|P0DPQ7\|GCOA_AMYS7 | K23526 | 6.9e-245 | 812.7 | CYP255A2 | 0.0 | 552 |
| sp\|P14762\|CPXI_PRIM2 | K21113 | 6.8e-213 | 707.5 | CYP106A1 | 0.0 | 853 |
| sp\|P18326\|CPXE_STRGO | K21146 | 1.7e-248 | 824.8 | CYP105A1 | 0.0 | 824 |
| sp\|P18327\|CPXF_STRGO | K17876 | 6.6e-142 | 473.5 | CYP105B1 | 0.0 | 803 |
| sp\|P23296\|CPXG_STRSQ | K17876 | 3.9e-113 | 378.7 | CYP105C1 | 0.0 | 756 |
| sp\|P24466\|CPXC_RHIRD | K21033 | 4.5e-175 | 582.8 | CYP103A1 | 0.0 | 863 |
| sp\|P24467\|CPXD_RHIRD | K21034 | 1.3e-214 | 712.8 | CYP104A1 | 0.0 | 840 |
| sp\|P27632\|CPXM_BACSH | K23138 | 5.6e-167 | 556.2 | CYP109A1 | 0.0 | 837 |
| sp\|P29980\|CPXN_NOSS1 | K23139 | 1.1e-191 | 637.7 | CYP110A1 | 0.0 | 947 |
| sp\|P33006\|CPXL_PSESP | K24391 | 7.5e-230 | 763.6 | CYP108A1 | 0.0 | 894 |
| sp\|P33271\|CPXK_SACEN | K21115 | 2.9e-287 | 952.3 | CYP107B1 | 0.0 | 811 |
| sp\|P43492\|THCB_RHOER | K16046 | 7.5e-69 | 232.7 | CYP116A1 | 0.0 | 908 |
| sp\|P46373\|FAS1_RHOFA | K21035 | 8.4e-283 | 937.5 | CYP 1050,00 | 0.0 | 796 |
| sp\|P48635\|ERYK_SACEN | K14370 | 2,00E-213 | 709.1 | CYP113A1 | 0.0 | 793 |
| sp\|P53554\|BIOI_BACSU | K16593 | 1.8e-210 | 699.5 | CYP107H1 | 0.0 | 825 |
| sp\|P55540\|CPXU_SINFN | K21118 | 9.7e-270 | 895.2 | CYP117A2 | 0.0 | 893 |
| sp\|P55543\|CPXR_SINFN | K21117 | 1.8e-257 | 854.2 | CYP114A2 | 0.0 | 937 |
| sp\|P55544\|CPXP_SINFN | K21116 | 7.2e-268 | 888.3 | CYP112A2 | 0.0 | 808 |
| sp\|P59954\|CP132_MYCBO | K21164 | 1,00E-100 | 338.4 | CYP132A1 | 0.0 | 940 |
| sp\|P63708\|CP123_MYCBO | K23138 | 2.1e-99 | 333.7 | CYP123A1 | 0.0 | 814 |
| sp\|P63710\|CP125_MYCBO | K15981 | 2.9e-217 | 721.6 | CYP125A1 | 0.0 | 901 |
| sp\|P63712\|CP126_MYCBO | K15981 | 3.2e-112 | 375.7 | CYP126A1 | 0.0 | 836 |
| sp\|P63714\|CP128_MYCBO | K21199 | 1,00E-223 | 743.5 | CYP128A1 | 0.0 | 988 |
| sp\|P63716\|C135B_MYCBO | K24241 | 1.6e-199 | 663.4 | CYP135B1 | 0.0 | 939 |
| sp\|P63718\|CP138_MYCBO | K24192 | 2.1e-235 | 781.7 | CYP138A1 | 0.0 | 892 |
| sp\|P63720\|CP139_MYCBO | K23139 | 2.8e-77 | 260.6 | CYP139A1 | 0.0 | 867 |

| | | | | | | |
|---|---|---|---|---|---|---|
| sp\|P63722\|CP140_MYCBO | K20789 | 1.5e-189 | 631.0 | CYP140A1 | 0.0 | 878 |
| sp\|P9WPL0\|CP144_MYCTO | K21200 | 2.2e-204 | 679.5 | CYP144A1 | 0.0 | 882 |
| sp\|P9WPL1\|CP144_MYCTU | K21200 | 2.2e-204 | 679.5 | CYP144A1 | 0.0 | 882 |
| sp\|P9WPL4\|CP142_MYCTO | K16046 | 1.2e-208 | 693.0 | CYP142A1 | 0.0 | 757 |
| sp\|P9WPL5\|CP142_MYCTU | K16046 | 6.3e-220 | 730.2 | CYP142A1 | 0.0 | 811 |
| sp\|P9WPL6\|CP141_MYCTO | K13074 | 6.7e-81 | 272.7 | CYP141A1 | 0.0 | 814 |
| sp\|P9WPL7\|CP141_MYCTU | K13074 | 6.8e-81 | 272.7 | CYP141A1 | 0.0 | 816 |
| sp\|P9WPL8\|CP140_MYCTO | K20789 | 1.5e-189 | 631.0 | CYP140A1 | 0.0 | 878 |
| sp\|P9WPL9\|CP140_MYCTU | K20789 | 1.5e-189 | 631.0 | CYP140A1 | 0.0 | 878 |
| sp\|P9WPM0\|CP139_MYCTO | K23139 | 2.8e-77 | 260.6 | CYP139A1 | 0.0 | 867 |
| sp\|P9WPM1\|CP139_MYCTU | K23139 | 2.8e-77 | 260.6 | CYP139A1 | 0.0 | 867 |
| sp\|P9WPM2\|CP138_MYCTO | K24192 | 2.1e-235 | 781.7 | CYP138A1 | 0.0 | 892 |
| sp\|P9WPM3\|CP138_MYCTU | K24192 | 2.1e-235 | 781.7 | CYP138A1 | 0.0 | 892 |
| sp\|P9WPM4\|CP137_MYCTO | K24241 | 2,00E-129 | 432.4 | CYP137A1 | 0.0 | 938 |
| sp\|P9WPM5\|CP137_MYCTU | K24241 | 2,00E-129 | 432.4 | CYP137A1 | 0.0 | 938 |
| sp\|P9WPM6\|CP136_MYCTO | K24392 | 9.4e-86 | 288.7 | CYP136A1 | 0.0 | 1021 |
| sp\|P9WPM7\|CP136_MYCTU | K24392 | 9.4e-86 | 288.7 | CYP136A1 | 0.0 | 1021 |
| sp\|P9WPM8\|C135B_MYCTO | K24241 | 1.6e-199 | 663.4 | CYP135B1 | 0.0 | 939 |
| sp\|P9WPM9\|C135B_MYCTU | K24241 | 1.6e-199 | 663.4 | CYP135B1 | 0.0 | 939 |
| sp\|P9WPN0\|C135A_MYCTO | K24241 | 1.2e-187 | 624.2 | CYP135A1 | 0.0 | 912 |
| sp\|P9WPN1\|C135A_MYCTU | K24241 | 1.2e-187 | 624.2 | CYP135A1 | 0.0 | 912 |
| sp\|P9WPN2\|CP132_MYCTO | K21164 | 1,00E-100 | 338.4 | CYP132A1 | 0.0 | 940 |
| sp\|P9WPN3\|CP132_MYCTU | K21164 | 4.3e-100 | 336.4 | CYP132A1 | 0.0 | 942 |
| sp\|P9WPN4\|CP130_MYCTO | K21119 | 8.4e-229 | 760.0 | CYP130A1 | 0.0 | 829 |
| sp\|P9WPN5\|CP130_MYCTU | K21119 | 8.4e-229 | 760.0 | CYP130A1 | 0.0 | 829 |
| sp\|P9WPN6\|CP128_MYCTO | K21199 | 1,00E-223 | 743.5 | CYP128A1 | 0.0 | 988 |
| sp\|P9WPN7\|CP128_MYCTU | K21199 | 1,00E-223 | 743.5 | CYP128A1 | 0.0 | 988 |

| | | | | | | |
|---|---|---|---|---|---|---|
| sp\|P9WPN8\|CP126_MYCTO | K15981 | 3.2e-112 | 375.7 | CYP126A1 | 0.0 | 836 |
| sp\|P9WPN9\|CP126_MYCTU | K15981 | 3.2e-112 | 375.7 | CYP126A1 | 0.0 | 836 |
| sp\|P9WPP0\|CP125_MYCTO | K15981 | 2.9e-217 | 721.6 | CYP125A1 | 0.0 | 901 |
| sp\|P9WPP1\|CP125_MYCTU | K15981 | 2.9e-217 | 721.6 | CYP125A1 | 0.0 | 901 |
| sp\|P9WPP4\|CP123_MYCTO | K23138 | 2.1e-99 | 333.7 | CYP123A1 | 0.0 | 814 |
| sp\|P9WPP5\|CP123_MYCTU | K23138 | 2.1e-99 | 333.7 | CYP123A1 | 0.0 | 814 |
| sp\|P9WPP6\|CP121_MYCTO | K17483 | 6.7e-191 | 634.7 | CYP121A1 | 0.0 | 800 |
| sp\|P9WPP7\|CP121_MYCTU | K17483 | 6.7e-191 | 634.7 | CYP121A1 | 0.0 | 800 |
| sp\|P9WPP8\|CP51_MYCTO | K05917 | 3.1e-176 | 588.1 | CYP51B1 | 0.0 | 930 |
| sp\|P9WPP9\|CP51_MYCTU | K05917 | 3.1e-176 | 588.1 | CYP51B1 | 0.0 | 930 |
| sp\|Q00441\|CPXJ_SACEN | K14366 | 2.6e-248 | 824.0 | CYP107A1 | 0.0 | 804 |
| sp\|Q06069\|CPXM_PRIMG | K21114 | 1,00E-281 | 934.1 | CYP106A2 | 0.0 | 846 |
| sp\|Q0S7M1\|CP125_RHOJR | K15981 | 1.4e-220 | 732.5 | CYP125A14P | 0.0 | 975 |
| sp\|Q53215\|Y4VG_SINFN | K21569 | 9.3e-90 | 301.4 | CYP127A1 | 0.0 | 846 |
| sp\|Q54823\|DNRQ_STRPE | K15952 | 7.4e-304 | 1007.5 | CYP131A1 | 0.0 | 845 |
| sp\|Q59203\|CPXP_BRADU | K21116 | 1,00E-266 | 884.5 | CYP112A1 | 0.0 | 803 |
| sp\|Q59204\|CPXR_BRADU | K21117 | 2.3e-257 | 853.8 | CYP114A3v1 | 0.0 | 793 |
| sp\|Q59205\|CPXU_BRADU | K21118 | 2,00E-264 | 877.6 | CYP117A3 | 0.0 | 816 |
| sp\|Q59723\|CPXO_PSEPU | K05525 | 1,00E-223 | 742.8 | CYP111A1 | 0.0 | 835 |
| sp\|Q59831\|CPS2_STRCC | K21146 | 4,00E-239 | 793.9 | CYP105A3 | 0.0 | 828 |
| sp\|Q59971\|DOXA_STRS5 | K15955 | 1.4e-295 | 980.1 | CYP129A1 | 0.0 | 840 |
| sp\|Q59990\|CP120_SYNY3 | K24392 | 1.4e-196 | 654.0 | CYP120A1 | 0.0 | 902 |
| sp\|Q5IZM4\|CP51_MYCVP | K05917 | 6.5e-179 | 597.0 | CYP51B1 | 0.0 | 939 |
| sp\|Q5J1R4\|NOCL_NOCUT | K19106 | 3.2e-262 | 869.6 | CYP107AC1 | 1.69e-144 | 415 |
| sp\|Q6N8N2\|CYPA2_RHOPA | K22553 | 1.4e-192 | 640.4 | CYP199A2 | 0.0 | 838 |
| sp\|Q825I8\|CYP28_STRAW | K17876 | 8.1e-212 | 703.7 | CYP105D7 | 0.0 | 804 |
| sp\|Q82IY3\|PTLI_STRAW | K15907 | 5.7e-247 | 820.1 | CYP183A1 | 0.0 | 911 |

| | | | | | | |
|---|---|---|---|---|---|---|
| sp\|Q83WF5\|MYCCI_MICGR | K24292 | 5.5e-233 | 772.8 | CYP105L2 | 0.0 | 768 |
| sp\|Q84HB6\|NCSB3_STRCZ | K20420 | 2,00E-167 | 559.7 | CYP154J1 | 0.0 | 837 |
| sp\|Q87AV9\|C1332_XYLFT | K16593 | 4.1e-93 | 313.0 | CYP133B2v2 | 0.0 | 803 |
| sp\|Q87AX5\|C1331_XYLFT | K16593 | 2,00E-101 | 340.4 | CYP133B1v2 | 0.0 | 830 |
| sp\|Q8RN03\|C5C4_AMYOR | K16447 | 3.1e-258 | 856.6 | CYP165C4 | 0.0 | 827 |
| sp\|Q8RN04\|C5B3_AMYOR | K16446 | 1.4e-258 | 857.9 | CYP165B3 | 0.0 | 806 |
| sp\|Q8RN05\|C5A3_AMYOR | K16445 | 2.3e-249 | 827.4 | CYP165A3 | 0.0 | 794 |
| sp\|Q8VQF6\|CINA_CITBR | K21120 | 1.3e-288 | 956.5 | CYP176A1 | 0.0 | 830 |
| sp\|Q93MI2\|DOXA_STRC0 | K15955 | 1,00E-277 | 921.2 | CYP129A2 | 0.0 | 756 |
| sp\|Q9FCA6\|C1582_STRCO | K13074 | 3.3e-212 | 705.2 | CYP158A2 | 0.0 | 719 |
| sp\|Q9KIZ4\|C167_SORCE | K16400 | 1.3e-260 | 864.7 | CYP167A1 | 0.0 | 840 |
| sp\|Q9KZF5\|C1581_STRCO | K19628 | 1.7e-250 | 831.0 | CYP158A1 | 0.0 | 811 |
| sp\|Q9L4U5\|AKNT_STRGJ | K15947 | 0 | 1020.6 | CYP131A1 | 3.61e-79 | 251 |
| sp\|Q9L9F9\|NOVI_STRNV | K12702 | 2.1e-281 | 932.8 | CYP163A1 | 0.0 | 839 |
| sp\|Q9PGC5\|C1331_XYLFA | K16593 | 5.4e-101 | 339.0 | CYP133B1v1 | 0.0 | 832 |
| sp\|Q9PGE6\|C1332_XYLFA | K16593 | 8.3e-95 | 318.6 | CYP133B2v2 | 0.0 | 803 |
| sp\|Q9ZAU3\|DOXA_STRPE | K15955 | 3.7e-294 | 975.4 | CYP129A2 | 0.0 | 828 |
| sp\|Q9ZGH8\|DES8_STRVZ | K16005 | 6.7e-277 | 918.6 | CYP131A1 | 3.21e-64 | 210 |
| sp\|Q9ZHQ1\|TYLH1_STRFR | K15991 | 1.1e-230 | 765.7 | CYP105L1 | 0.0 | 830 |
